# γδ17 T cell-stromal networks modulate matrix composition and vascularity in foreign body response

**DOI:** 10.1101/2025.09.30.679608

**Authors:** Anna Ruta, Kavita Krishnan, JiWon Woo, Joscelyn C. Mejías, Elise F. Gray-Gaillard, David R. Maestas, Helen Hieu Nguyen, Alexandra N. Rindone, Christopher Cherry, Michael Patatanian, Frank Haoning Yu, Brenda Yang, Connor Amelung, Christina D. King, Birgit Schilling, Sharon Gerecht, Elana J. Fertig, Locke Davenport Huyer, Drew M. Pardoll, Jennifer H. Elisseeff

## Abstract

Immune-stromal crosstalk governs tissue fibrosis, which is marked by dysregulated extracellular matrix (ECM) production and aberrant vasculature. Here, we investigate how γδ T cell interactions with stromal cells shape fibrosis in the foreign body response. During the acute reaction, type-1 (γδIFNγ) and type-17 (γδ17) effector subsets accumulated at the implant. While γδIFNγ decreased as fibrosis progressed, activated γδ17 persisted as dominant interleukin-17 producers. The γδ17 increased with aging and high-fat diet, both factors associated with chronic inflammation and fibrosis. Co-culture with γδ17 stimulated fibroblast expression of collagen genes and intercellular communication inference linked γδ T cell ligands to activation of ECM remodeling and vascular development programs in fibroblasts and endothelial cells. Finally, genetic deletion of γδ T cells altered expression of ECM components and increased vessel size within the fibrotic matrix. Altogether, our findings implicate γδ T cells in regulating stromal behavior to modulate composition and vascularity of fibrotic tissues.

## Introduction

γδ T cells regulate tissue structure and function by reacting to diverse environmental stimuli and interacting with neighboring cells under homeostatic and pathophysiological conditions^1^. As unconventional T cells with unique γδ T cell receptors (TCRs), their activation is not restricted to peptide antigen presentation in classic MHC molecules. Instead, γδ T cells broadly survey their surroundings and mount rapid innate-like responses upon recognition of antigens such as stress-liable ligands on damaged or transformed cells, pathogenic and commensal microbes, lipids, phosphoantigens, neuropeptides, and carbohydrates^2,3^. This versatility allows γδ T cells to serve critical roles in barrier surveillance and host defense^4^, while also regulating physiological processes ranging from body temperature control^5–7^ and dietary nutrient absorption^8^ to anxiety-like behaviors^9^ and short-term memory development^10^. Moreover, γδ T cells support tissue development, including new blood vessel formation^11–13^ and sympathetic innervation of peripheral tissues^6^, which are crucial processes for restoration of tissue integrity upon injury.

γδ T cells play pivotal roles in orchestrating tissue injury responses that depend on complex immune-stromal dynamics to determine outcomes of productive repair versus dysregulated fibrosis. γδ T cells enhance repair of numerous tissue types by secreting immunomodulatory and tissue growth factors^1^. For example, skin γδ T cells that sense damage signals from injured keratinocytes produce insulin-like and fibroblast growth factors to support re-epithelialization^14–17^. Likewise, gingival γδ T cells supply amphiregulin, an epidermal growth factor involved in repair, to prevent periodontal bone loss^18^. Activated γδ T cells that secrete immunomodulatory factors, such as interferon-γ (IFNγ) and interleukin (IL)-22, exert protective functions following lung injury^19–23^ and mitigate liver fibrosis^24,25^. Furthermore, γδ T cells secreting IL17 promote osteogenesis and bone fracture healing^26,27^, muscle repair following myotoxic injury^28^, lung epithelium recovery after viral infection^29^, and skin wound re-epithelialization by activating hypoxia-inducible transcription factors (TFs) in injured epithelial cells^30^.

While γδ T cells respond rapidly to tissue injury and support productive repair, depending on the context and nature of their interactions with the stromal compartment, they can also contribute to fibrosis. Fibrosis is a pathological process characterized by excessive extracellular matrix (ECM) deposition and abnormal vascular structure, with resulting tissue stiffening and dysfunction contributing to morbidity across a range of diseases. Chronic inflammation can drive and exacerbate fibrosis, however immune regulation of fibroblast and endothelial cell (EC) behavior, in particular by γδ T cells, is less understood. Persistent and elevated levels of IL17 are associated with fibrosis across multiple tissue types^31^. As potent IL17 producers, γδ T cells contribute to inflammation and fibrosis in preclinical models of arthritis^32,33^, psoriasis^34,35^, silicosis-induced lung fibrosis^36^, ischemic and traumatic brain injuries^37–39^, liver fibrosis^40,41^, and glomerulonephritis^42^. Further exploration of mechanisms by which γδ T cells engage with resident stromal populations, and the downstream effect of these interactions on shaping tissue structure in fibrosis, is needed.

In this study, we investigate how γδ T cells interact with stromal cells to regulate repair and fibrosis using a foreign body response (FBR)-induced model of soft tissue fibrosis. The FBR remains a significant clinical challenge that leads to dysfunction or failure in 10-30% of medical implants and devices^43–45^. The implantation of synthetic biomaterials reliably induces immune reactions that drive fibroblast activation and fibrotic encapsulation^46,47^, thus enabling mechanistic studies of fibrosis development and progression. Previously, we found that implant-associated fibrosis is linked to elevated IL17, senescent stromal cell accumulation, fibrotic macrophage activation, and prolonged granulocyte infiltration^48–50^. Here, we show that while multiple γδ T cell subtypes, including type-1 (γδIFNγ) and type-17 (γδ17), accumulated during acute wound healing, γδ17 persisted along with FBR-induced fibrosis development. Although γδ17 were a relatively small T cell subset in the fibrotic environment, they accounted for a third of the activated IL17-producing T cells and produced higher levels of IL17a than conventional CD4^+^ T_H_17 cells. The γδ17 further expanded with obesity and aging, both systemic conditions associated with increased fibrotic predisposition. Computational and *in vitro* experimental analyses revealed that γδ T cells expressed bioactive ligands (e.g., cytokines, chemokines, and growth factors) that can activate transcriptional programs in fibroblasts and ECs associated with ECM remodeling and vascular development. Mice lacking γδ T cells differentially expressed various ECM components, including collagens and proteoglycans associated with fibrocartilage, at transcriptomic and proteomic levels. Furthermore, the absence of γδ T cells significantly increased EC numbers and blood vessel size, leading to altered vascular architecture within the fibrotic tissue. Collectively, we demonstrate that signaling from γδ T cells can influence stromal cell phenotypes, identifying potential pathways through which they may impact fibrotic tissue structure and composition. These insights into the complex cellular and molecular interactions between γδ T cells and stromal populations may inform new therapeutic strategies to target fibrotic pathologies.

## Results

### γδ T cells accumulate in fibrotic tissue of a biomaterial implant model

Tissue injury is marked by acute immune infiltration and activation, typically followed by repair and resolution. However, sustained injury and inflammation, such as in the presence of biomaterial implants, can impede healing and result in fibrosis driven by stromal cell accumulation and excessive ECM deposition^48,49^. To characterize the γδ T cell response in tissue injury and fibrosis, we performed bilateral volumetric muscle loss (VML) injury in murine quadricep muscles and implanted synthetic materials into the defect sites^48,49^. This preclinical model induces robust fibrotic implant encapsulation and enables investigation of cellular and molecular mechanisms contributing to tissue fibrosis. The implantation of polycaprolactone (PCL) particulate, a well-established inducer of local tissue fibrosis^49,51,52^, significantly increased the number of γδ T cells (**Fig 1A**). In saline-treated injury only (no implant) controls, γδ T cell numbers peaked at 1 week (wk) during the acute inflammatory phase of wound healing (203.3±73.3 cells/muscle, mean±SD) and were largely absent by 6 wks as inflammation resolved (31.6±14.6 cells/muscle), limiting characterization at later time points (**Fig S1A**). While γδ T cells still expanded early with the PCL implant (1 wk: 1620.8±348.8 cells/muscle), their high levels persisted at later time points associated with implant fibrosis (12 wk: 1239.5±664.2 cells/muscle) (**Fig S1A**). The PCL implant environment had significantly more γδ T cells (5-fold cell count increase at 6 wks) than other polymeric materials, polyethylene (PE) and silicone, also commonly used in biomedical applications (**Fig 1A**, **S1B**).

**Fig 1.**
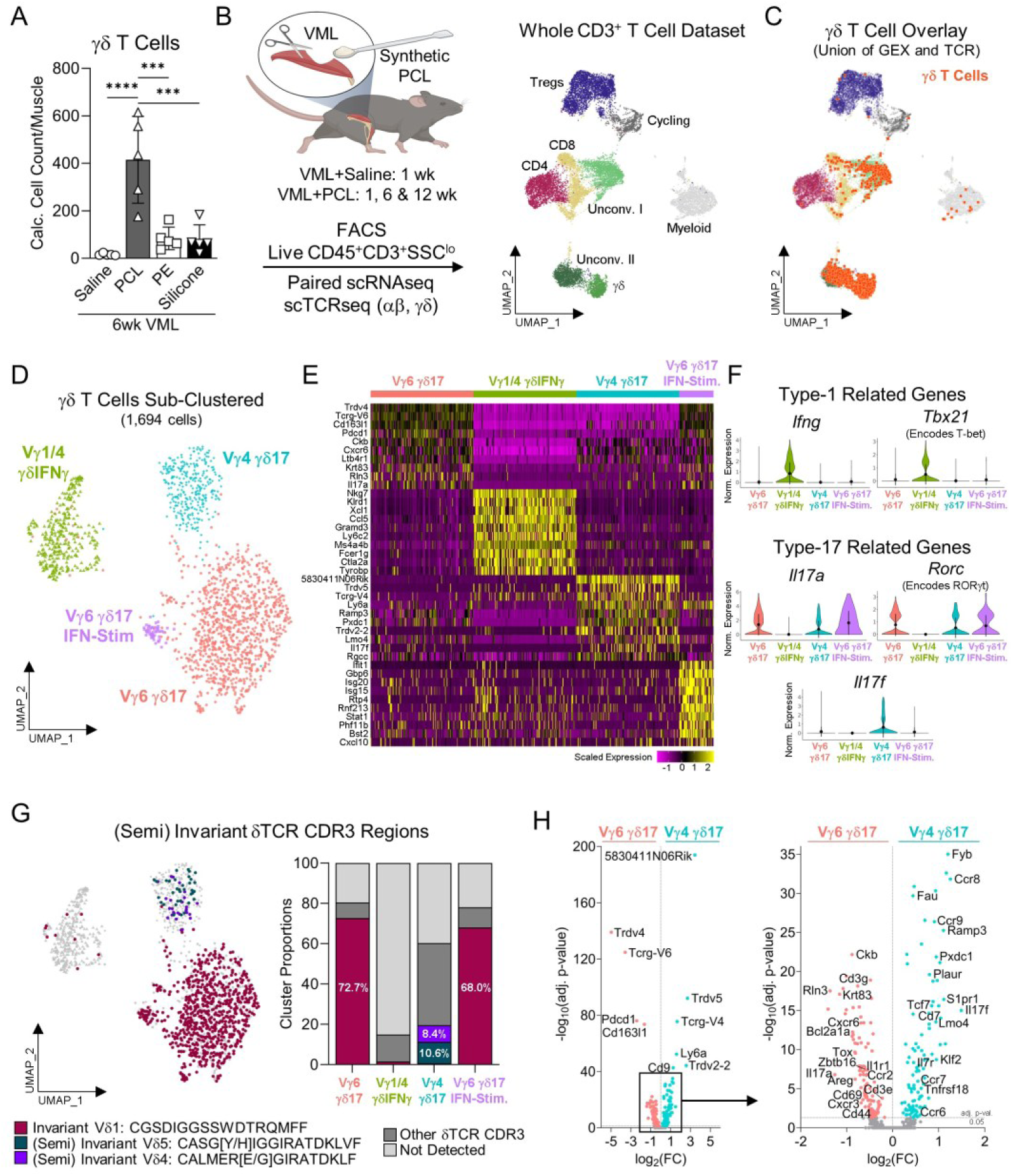
Murine γδ T cells found in peri-implant fibrotic tissue exhibit multiple effector phenotypes. **(A)** Flow cytometric quantification of γδ T cells in murine quadricep muscles 6 weeks (wks) after volumetric muscle loss (VML) injury with synthetic biomaterial implantation. PCL: polycaprolactone, PE: polyethylene. **(B)** Experimental schematic of paired, single cell RNA- and T cell receptor-sequencing of FACS-isolated CD3^+^ T cells at various time points following VML injury, with and without PCL implants (n=3-5 per PCL time point, n=18 pooled for 1 wk injury only) (left). UMAP visualization of whole CD3^+^ T cell dataset (14,995 T cells) depicting CD8, CD4, Tregs, and multiple unconventional (uncov.) T cells (right). **(C)** Overlay of γδ T cells (orange) on CD3^+^ T cell UMAP. γδ T cells computationally identified by expression of γ and/or δ TCR variable region genes and/or TCRs with an absence of α/β counterparts. **(D)** UMAP of γδ T cell clusters (1,694 cells). **(E)** Scaled expression heatmap of top 10 differentially expressed genes per γδ T cell cluster. **(F)** Violin plots of canonical type-1 (*Ifng* and *Tbx21*) and type-17 (*Il17a*, *Il17f*, and *Rorc*) gene expression per γδ T cell cluster. **(G)** Top 3 most abundant (semi)invariant δTCR sequences overlayed on γδ T cell UMAP (left). Proportion of cells within the γδ T cell clusters that express select δTCR sequences (right). **(H)** Volcano plot of differential gene expression between Vγ6 γδ17 and Vγ4 γδ17 clusters. **(Statistics)** Bar and violin plots: mean±SD. Data analyzed using ordinary one-way ANOVA with Tukey’s multiple comparisons test (A). NS: Not significant p>0.05, * p<0.05, ** p<0.01, *** p<0.001, **** p<0.0001.

### γδ T cells in peri-implant fibrotic tissue exhibit multiple effector phenotypes

To comprehensively phenotype the γδ T cells enriched in the PCL implant environment, we isolated T cells (Live CD45^+^ CD3^+^) via fluorescent-activated cell sorting (FACS) at 1, 6, and 12 wks post VML injury with PCL implantation and performed paired single-cell RNA- and TCR-sequencing (scRNA/TCRseq) (n=3-5 per time point). An analogous analysis of T cells from 1 wk injury only controls was also performed, for which we pooled 18 biological replicates due to low T cell numbers in this condition. The collective dataset captured multiple T cell subtypes (14,995 cells) including CD8, CD4, regulatory T cells, and mixed unconventional T cells (**Fig 1B**, **S1C-D**). We computationally identified γδ T cells based on expression of gamma or delta TCR variable region genes (*Trgv*, *Tcrg-V,* or *Trdv*) and/or TCRs (γ or δ chains) in the absence of alpha or beta TCR counterparts (**Fig 1C**, **S1E-G**). Small populations of myeloid cells, cycling cells, and doublets were detected and excluded in subsequent analyses (**Fig S2**). We identified four distinct γδ T cell subsets (1,694 cells) marked by unique Vγ chains and effector phenotypes that reportedly correspond with distinct developmental waves^1^, including one γδIFNγ subset and three γδ17 subsets (**Fig 1D-F**, **S3**). These γδ T cell subsets were detected at varying proportions at all PCL implant time points as well as in the injury only controls (**Fig S3A-C**).

The Vγ1/4 γδIFNγ subset expressed *Tcrg-V1* and *Tcrg-V4* Vγ chains (**Fig S3D**) and hallmark type-1 genes including *Ifng*, *Tbx21* (encodes TF T-bet), and *Eomes* (**Fig 1E-F**, **S4A-B**). This subset had high expression of the co-stimulatory receptor *Cd27*, which is a critical functional regulator of γδIFNγ development, as well as the common γδIFNγ markers *Ly6c2, Klrb1c* (encodes NK1.1), and *Il2rb*^53–58^ (**Fig S4A-B**). Additionally, the subset’s significant differentially expressed genes considerably overlapped (8 out of 13 genes) with a recently generated transcriptomic signature of activated peripheral γδIFNγ^59^. These cells originally clustered with natural killer (NK) T cells, likely driven by shared expression of effector- and killer cell lectin-like receptor-related genes (*Nkg7, Klrc1*, *Klrd1*) (**Fig S3E, S4B**). Accordingly, ranked gene set enrichment analysis (GSEA) revealed that the Vγ1/4 γδIFNγ subset was significantly enriched for pathways involved in cell killing, NK cell-related immunity, IFNγ response, and lymphocyte activation (**Fig S4C-D**).

The three γδ17 subsets expressed canonical type-17 genes including *Il17a*, *Il23r*, *Rorc* (encodes TF RORγt), and *Maf* as well as the common γδ17 markers *Cd44*, *Il7r* (encodes CD127), and *Ccr6*^58,60–63^ (**Fig 1E-F**, **S5A**). The largest subset, Vγ6 γδ17, consisted of Vγ6Vδ1 cells marked by *Tcrg-V6* and *Trdv4* (encodes Vδ1) expression (**Fig 1E**, **S3D**). Vγ6Vδ1 cells are known to develop in the fetal thymus, where they are “pre-programmed” towards a γδ17 phenotype with a highly restricted TCR V(D)J gene usage, and subsequently disseminate to various peripheral tissues where they persist as self-renewing populations^1,64–71^. Accordingly, 90.4% of the Vγ6 γδ17 cells with a detected δTCR displayed the invariant Vδ1Dδ2Jδ2 sequence, though limited γTCR capture prevented detection of the corresponding invariant Vγ6Jδ1 sequence (**Fig 1G**, **S6**). A small Vγ6 γδ17 IFN-Stimulated subset shared a similar transcriptional profile with the main Vγ6 γδ17 subset and also expressed the invariant Vδ1Dδ2Jδ2 sequence, but distinctly expressed genes induced by type I IFNs (*Isg15*, *Ifit1*, *Gbp6*, *Cxcl10*) and was enriched in pathways related to type I IFN response, IFNγ response, and TNFα signaling via NFκβ (**Fig S5B-D**). The other main γδ17 subset, Vγ4 γδ17, largely expressed *Tcrg-V4* and *Trdv5* (**Fig 1E-F**, **S3D**) and is typically enriched in secondary lymphoid organs^1,68,71,72^. Although this subset exhibited greater TCR diversity, 21.3% and 13.9% of the Vγ4 γδ17 cells with a detected δTCR expressed the semi-invariant Vδ5Dδ2Jδ1 and Vδ4Dδ2Jδ1 sequences, respectively^63,68,71,72^ (**Fig 1G**, **S6**).

While the Vγ6 γδ17 and Vγ4 γδ17 subsets displayed overlapping γδ17 features, their differential expression of IL17 cytokines (*Il17a* by Vγ6 γδ17 and *Il17f* by Vγ4 γδ17) and chemokine receptors (*Cxcr6*, *Cxcr3* and *Ccr2* by Vγ6 γδ17 and *Ccr9*, *Ccr8* and *Ccr7* by Vγ4 γδ17) suggests distinct responses within the fibrotic implant environment (**Fig 1H**). The Vγ4 γδ17 subset exhibited higher expression of T cell migration genes (*S1pr1, Klf2*), which corresponds with their reported trafficking to sites of inflammation^73^. Additional transcriptional patterns that differentiate the two subsets, including expression of scavenger receptors (*Cd163l1* encoding Scart1 and *5830411N06Rik* encoding Scart2), co-inhibitory receptors (*Pdcd1* encoding PD1), *Cd9*, and anti-apoptotic Bcl2 family members appear to be conserved across different peripheral tissues, including lung and skin^70,74,75^. Collectively, the local PCL implant environment consists of a heterogeneous γδ T cell population, including distinct γδIFNγ and γδ17 effector subsets.

### Acute tissue injury activates both γδIFNγ and γδ17 subsets with persistence of expanded γδ17 in chronic implant fibrosis

We temporally profiled γδ T cells to assess whether their response changes during fibrotic tissue development. Proportional analysis of scRNAseq clusters suggested dynamic shifts in γδ T cell composition (**Fig 2A**, **S3B**). The Vγ6 γδ17 consistently made up approximately 60% of γδ T cells, while the proportion of Vγ1/4 γδIFNγ decreased (1 wk: 18.7±13.1%, 12 wk: 2.5±1.8%) and Vγ4 γδ17 increased (1 wk: 11.2±3.4%, 12 wk: 34.4±26.6%) over time. The small Vγ6 γδ17 IFN-Stimulated subset was mainly present at 1 wk. We validated these temporal shifts with high-parameter flow cytometry performed at acute (3 day, 1 wk), intermediate (3 wk), and chronic (6 wk, 12 wk) time points following PCL implantation. The γδ T cell effector subsets were defined by established surface markers corroborated by our scRNAseq results, specifically NK1.1 and Ly6C to mark γδIFNγ and CD127 and CD44^hi^ to mark γδ17 (**Fig 2B-C**, **S7A-B**). Dimensional reduction of γδ T cells across all time points (30,242 cells) revealed three main subsets: Vγ1/4^+^ γδIFNγ, Vγ4^+^ γδ17, and Vγ1/4^-^ γδ17, which corresponded to the primary scRNAseq clusters (**Fig 2D**, **S7C-D**). We inferred that the Vγ1/4^-^ γδ17 were likely Vγ6^+^ based on scRNAseq clusters and brighter CD3 expression relative to Vγ4^+^ cells^76^ (**Fig S7D**), which was confirmed in subsequent studies with an anti-Vγ6 antibody (**Fig S7E**).

**Fig 2.**
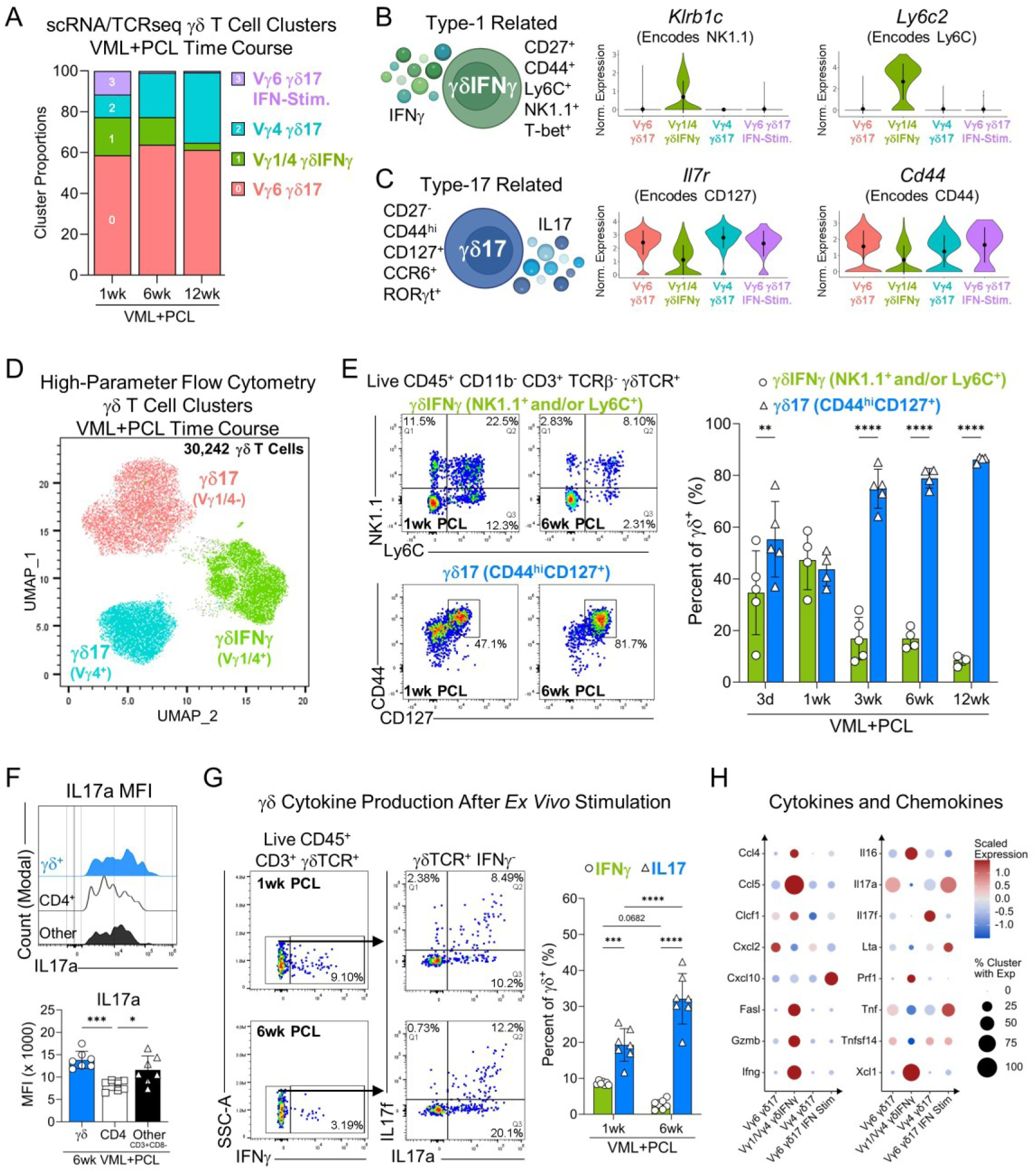
Acute wound healing activates both γδIFNγ and γδ17, but γδ17 persist as peri-implant fibrosis progresses. **(A)** Proportion of each scRNAseq γδ T cell cluster per VML+PCL time point (1, 6, and 12 wk). **(B)** Schematic and violin plots of γδIFNγ marker genes (*Klrb1c* and *Ly6c2*). **(C)** Schematic and violin plots of γδ17 marker genes (*Il7r* and *Cd44*). **(D)** UMAP of main 3 γδ T cell subsets (30,242 cells) from VML+PCL time course (3 days, 1, 3, 6 and 12 wk) based on high-parameter spectral flow cytometry (n=4-5 per time point). **(E)** Representative flow plots (left) and quantification (right) of γδIFNγ (NK1.1^+^ and/or Ly6C^+^) and γδ17 (CD44^hi^CD127^+^) in VML+PCL time course. **(F)** Representative histogram (top) and median fluorescence intensity (MFI) quantification (bottom) for IL17a by various IL17a^+^ T cells (γδ T cells, CD4^+^ T cells, and Other: CD3^+^γδ^-^CD4^-^CD8^-^) at 6 wk VML+PCL. **(G)** Representative flow plots (left) and quantification (right) of γδ T cell cytokine production (IFNγ, IL17a, and IL17f) following *ex vivo* stimulation at 1 and 6 wk following VML+PCL. IL17 production is characterized as IL17a^+^ and/or IL17f^+^. **(H)** Row-scaled expression of secreted cytokines and chemokines by scRNAseq γδ T cell clusters (>10% of cells in at least one cluster and >0.1 average expression in at least one cluster). **(Statistics)** Bar and violin plots: mean±SD. Data analyzed using repeated measures one-way ANOVA (F) or two-way ANOVA (E and G) with Tukey’s multiple comparisons test. Only displaying statistics within time point for E. NS: Not significant p>0.05, * p<0.05, ** p<0.01, *** p<0.001, **** p<0.0001.

The acute inflammatory phase of wound healing consisted of both γδIFNγ (NK1.1^+^ and/or Ly6C^+^) and γδ17 (CD44^hi^CD127^+^); however, these subsets exhibited divergent kinetics as implant-associated fibrosis progressed (**Fig 2E**, **S7F**). The γδIFNγ significantly decreased over time in terms of proportion and counts (1 wk: 47.3±11.5%, 12 wk: 8.4±1.8%), whereas the expanded γδ17 persisted and ultimately comprised the majority of γδ T cells at chronic time points (1 wk: 43.7±6.6%, 12 wk: 86.1±1.1%) (**Fig 2E**, **S7F**).This prominent γδ17 response was conserved across different polymer material implants. The PE and silicone implants similarly showed a significant enrichment of γδ17 relative to γδIFNγ and a comparable Vγ chain distribution (**Fig S7G-H**). In contrast, a biologically derived scaffold composed of decellularized ECM, which promotes muscle regeneration by inducing a type-2 immune response^77^, exhibited a balanced ratio of γδIFNγ to γδ17 (**Fig S7G**).

### γδ T cells are a prominent source of IL17 in fibrotic implant environments

Next, we functionally assessed T cell cytokine production by performing intracellular cytokine staining following *ex vivo* stimulation. γδ T cells accounted for less than 5% of IFNγ-producing and over 30% of IL17-producing T cells in the PCL environment at 1 and 6 wks (**Fig S8A**). The γδ T cells produced less IFNγ than CD8^+^ or CD4^+^ (T_H_1) cells based on median fluorescence intensity (**Fig S8B**), but equal amounts of IL17f and significantly more IL17a than CD4^+^ (T_H_17) cells (**Fig 2F, S8C-D**), indicating they are a notable IL17 source in the fibrotic implant environment. IFNγ was not co-expressed with either IL17 cytokine, confirming strict distinction of γδ T cell effector phenotypes (**Fig S8E**). Consistent with the γδIFNγ and γδ17 kinetics (**Fig 2E**), the percentage of IFNγ^+^ γδ T cells decreased (1 wk: 8.6±0.5%, 6 wk: 2.7±1.3%), whereas the percentage of IL17^+^ (IL17a^+^ and/or IL17f^+^) γδ T cells significantly increased (1 wk: 19.3±4.5%, 6 wk: 32.1±7.50%) over time (**Fig 2G**). The significant decrease in IFNγ^+^ γδ T cell counts and consistency in IL17^+^ γδ T cell counts (**Fig S8F**) signifies that the proportional shift is due to contraction of IFNγ^+^ γδ T cells and maintenance, rather than expansion, of IL17^+^ γδ T cells at chronic implant time points.

The cytokines IL17a and IL17f share considerable structural homology, resulting in shared and distinct bioactivity^78–80^. They can both signal through the IL17RA-IL17RC receptor complex, but IL17a exhibits a stronger binding affinity to IL17RA^81–84^. In addition to functioning as homodimers, IL17a and IL17f can signal as an IL17a/f heterodimer^85,86^. The distribution of Vγ chains differed based on cytokine profile: IFNγ and IL17f/f were mainly produced by Vγ4^+^, IL17a/a by Vγ1/4^-^ (inferred Vγ6^+^), and IL17a/f evenly by Vγ4^+^ and Vγ1/4^-^ (**Fig S8G**). The majority of IL17^+^ γδ T cells produced the stronger binding IL17a, either as IL17a/a or IL17a/f, with the percentage of IL17a/a^+^ γδ T cells significantly increasing over time in the fibrotic PCL environment (**Fig S8H**).

In addition to IFNγ and IL17, our scRNAseq analysis detected a myriad of other cytokines and chemokines expressed by the γδ T cells through which they can interact with and influence other cell types in the local PCL environment (**Fig 2H**, **S9A**). For example, the Vγ1/4 γδIFNγ subset exhibited strong expression of potent chemoattractants and pro-inflammatory factors, such as *Ccl5* and *Il16,* that promote immune cell recruitment and activation (**Fig S9B**). Similarly, the Vγ6 γδ17 and Vγ4 γδ17 subsets displayed high, scaled expression of *Cxcl2*, a known chemoattractant for neutrophils and macrophages, whereas the Vγ6 γδ17 IFN-Stimulated subset highly expressed *Cxcl10*. Beyond secreted factors mediating chemotaxis and activation, the Vγ1/4 γδIFNγ subset expressed factors associated with cytotoxicity including *Xcl1*, *Gzmb* (encodes Granzyme B), *Prf1* (encodes Perforin), and *Fasl* that aligned with enrichment in cell killing pathways (**Fig S4D**, **S9C**). Further, all four γδ T cell subsets expressed *Tnf*, which encodes the pro-inflammatory and cytotoxic tumor necrosis factor-alpha (TNFα), albeit at varying levels. Taken together, γδ T cells express a broad range of secreted immunomodulatory factors to actively shape the inflammatory landscape associated with tissue fibrosis.

### γδ T cells in the fibrotic implant environment are a mix of migratory and tissue-resident cells

We expanded our γδ T cell profiling beyond the local implant environment to consider draining inguinal lymph nodes (iLNs) that serve as important hubs for immune surveillance and activation. VML injury, both with and without PCL implants, significantly increased γδ T cells in the iLNs during the acute inflammatory phase of wound healing in comparison to uninjured controls (**Fig S10A**). The iLNs consisted primarily of Vγ1^+^ and Vγ4^+^ cells as opposed to the Vγ6^+^ cells found in the local PCL environment (**Fig S10B**). The amount of γδIFNγ in iLNs significantly increased with VML injury irrespective of PCL implant, while the amount of γδ17 did not change (**Fig S10C-E**). In contrast to the dominant IL17^+^ γδ T cell response within the local PCL environment, the iLNs had significantly more IFNγ^+^ γδ T cells at 6 wks (**Fig S10F-G**). Collectively, the γδ T cell response observed at the local implant site is distinct from that in the regional draining iLNs.

To discern whether γδ T cells present in the local PCL environment are tissue-resident or infiltrating from draining LNs, we pharmacologically antagonized the sphingosine-1-phosphate 1 receptor (S1P1R) using Fingolimod (FTY720 HCl) to inhibit T cell egress from LNs (**Fig S11A-B**). The FTY720 significantly decreased CD4^+^ and CD8^+^ T cells in the local PCL environment at 6 wks, but the γδ T cells were broadly unchanged (**Fig S11C-D**). However, FTY720 differentially impacted γδ T cell subsets by significantly decreasing the count and percentage of Vγ4^+^ cells, while not affecting Vγ6^+^ cell counts (**Fig S11E-F**). These results support that Vγ6^+^ cells are tissue-resident, whereas Vγ4^+^ cells that have higher *S1p1r* expression (**Fig 1H**) are migratory and can home to sites of inflammation from lymphoid tissues.

### High fat diet and age expand γδ17 in implant-associated fibrosis

Host intrinsic and extrinsic factors can influence tissue fibrosis. For example, obesity is associated with increased tissue fibrosis (e.g., adipose, liver, and lung) and elevated risk of many fibrosis-related diseases^87–92^. Interestingly, γδ17 can uptake and store lipids, are programmed to utilize oxidative metabolism, and increase systemically in mice on a high-fat diet (HFD)^93–95^. We investigated whether HFD-induced obesity can impact implant-associated fibrosis and its accompanying γδ17 response. We placed mice on HFD (Fat: 60 kcal%) or low-fat diet control (LFD, Fat: 10 kcal%) regimens for 8 wks and performed VML injury with PCL implantation 2 wks after diet initiation (**Fig S12A**). Mice on HFD gained significantly more weight (57.7±14.3% change from baseline) than LFD controls (14.0±4.3% change from baseline) over the 8 wk period (**Fig 3A**, **S12B**). In addition to systemic weight gain and accumulation of adipose tissue (**Fig S12C**), mice on HFD had ectopic fat deposition both within the peri-implant fibrotic tissue and at the muscle-implant interface (**Fig 3B**). HFD heightened chronic inflammation within the 6wk PCL environment as indicated by increased gene expression of pro-inflammatory factors such as *Tnf*, *Il1b*, and *Nos2* (**Fig 3C**, **S12D**). However, expression of fibrosis (*Col1a1*, *Col3a1*, *Col5a3*, *Fap, S100a4*) and growth factor (*Pdgfa*, *Pdgfb*, *Vegfa, Tgfb1*) genes within the PCL environment was largely unaffected by HFD, except for a significant decrease in *Tgfb2* and increase in the senescence marker *Cdkn2a* (**Fig S12E-G**).

**Fig 3.**
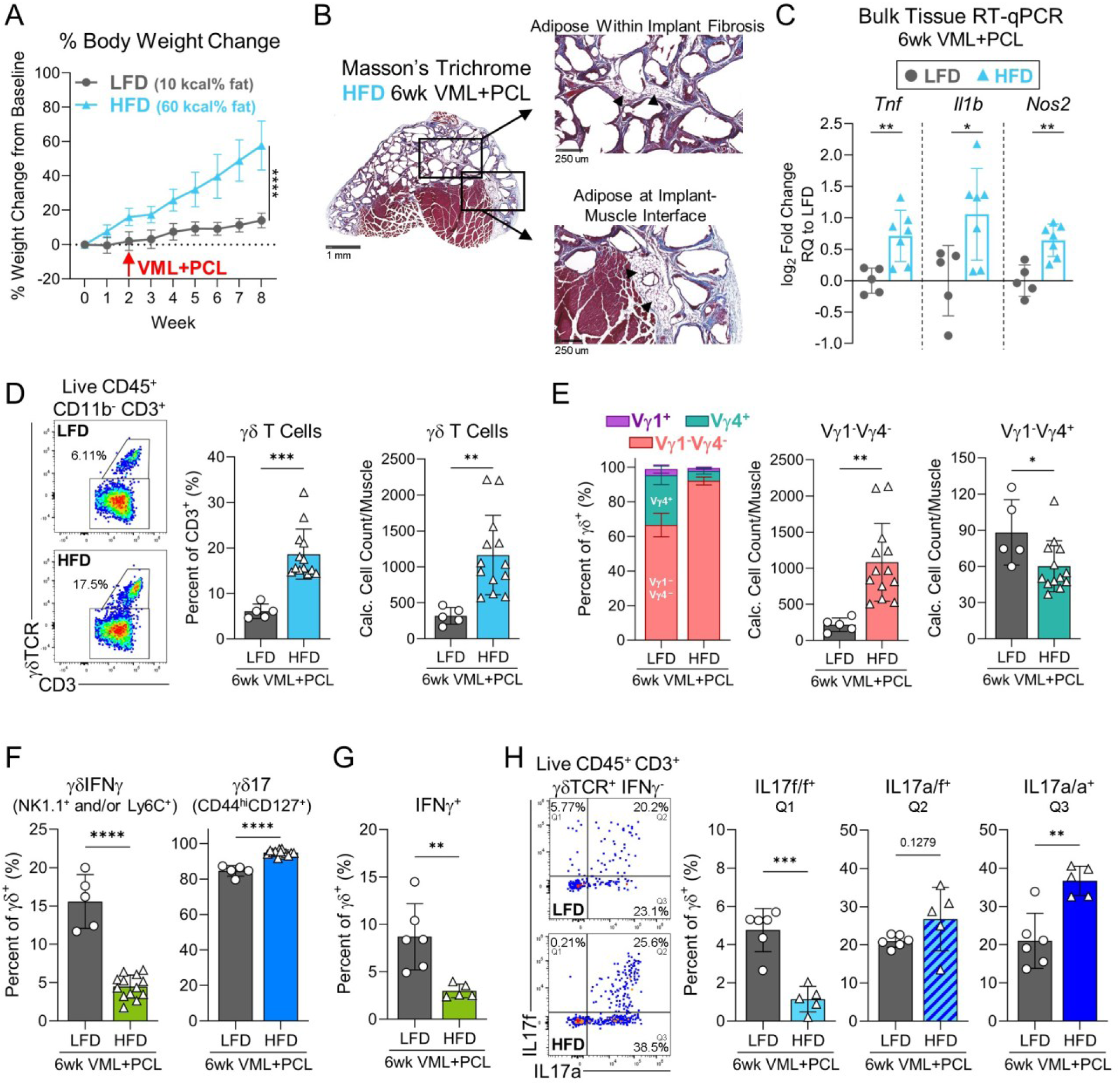
High fat diet increases γδ17 in peri-implant fibrotic tissue. **(A)** Longitudinal change in body weight relative to baseline (0 wk) of mice fed low-fat diet (LFD, 10 kcal% fat) (n=10) or high-fat diet (HFD, 60 kcal% fat) (n=20). VML+PCL was performed 2 wks after initiation of diet regimens. **(B)** Representative Masson’s Trichrome staining of fibrotic muscle tissue 6 wks after VML+PCL in HFD-fed mouse. Ectopic adipose deposition (black triangle) located within fibrotic tissue and at muscle-implant interface (scale bar: left 1 mm, right 250 μm). **(C)** Pro-inflammatory gene expression (*Tnf*, *Il1b*, and *Nos2*) in muscle-implant bulk tissue of HFD-fed mice relative to LFD-fed controls using RT-qPCR. **(D)** Representative flow plots (left) and quantification (middle, right) of γδ T cells in quadricep muscles of LFD- and HFD-fed mice 6 wks after VML+PCL. **(E)** Distribution and quantification of Vγ chain expression by muscle γδ T cells. **(F)** Quantification of γδIFNγ (NK1.1^+^ and/or Ly6C^+^) and γδ17 (CD44^hi^CD127^+^) in muscles of LFD- and HFD-fed mice 6 wks after VML+PCL. **(G)** Quantification of IFNγ production by γδ T cells following *ex vivo* stimulation. **(H)** Representative flow plots (left) and quantification (right) of IL17 production by γδ T cells following *ex vivo* stimulation including only IL17a^+^ (IL17a/a), only IL17f^+^ (IL17f/f), or dual IL17a^+^ and IL17f^+^ (IL17a/f) in muscles of LFD- and HFD-fed mice 6 wks after VML+PCL. **(Statistics)** Line and bar graphs: mean±SD. Data analyzed using unpaired two-tailed student t-test (A: only wk 8, C-H). NS: Not significant p>0.05, * p<0.05, ** p<0.01, *** p<0.001, **** p<0.0001.

γδ T cells significantly increased within the fibrotic PCL environment in mice fed with HFD, both in terms of cell count (LFD: 318.9±117.5 cells/muscle, HFD: 1165±551.2 cells/muscle) and proportion of CD3^+^ T cells (LFD: 6.1±1.6%, HFD: 18.7±5.5%) (**Fig 3D**). This expansion was unique to γδ T cells and not observed in other T cell subsets including NKTs, CD4^+^, or CD8^+^ T cells (**Fig S13A**). The percentage of γδ T cells positively correlated with mouse body weight (**Fig S13B**). HFD altered the γδ T cell composition by significantly expanding the Vγ1/4^-^ (inferred Vγ6^+^) subset, while simultaneously contracting the Vγ1^+^ and Vγ4^+^ subsets (**Fig 3E**, **S13C**). Further, the proportion of γδIFNγ significantly decreased in mice fed with HFD (LFD: 15.6±3.5%, HFD: 4.5±1.5%), whereas γδ17 significantly increased in both proportion (LFD: 84.6±3.0%, HFD: 94.8±1.5%) and cell count (**Fig 3F**, **S13D**). Intracellular cytokine staining following *ex vivo* cell stimulation revealed a corresponding decrease in the percentage of IFNγ^+^ γδ T cells with HFD, but differential expression of IL17 cytokines (**Fig 3G-H**, **S13E**). Specifically, HFD increased the percentages and counts of γδ T cells that produced IL17a/a and IL17a/f, but decreased γδ T cells that produced IL17f/f (**Fig 3H**, **S13E**). Accordingly, the dominant producer of IL17a/a was the Vγ1/4^-^ subset that expanded with HFD, whereas IL17f/f was primarily produced by the Vγ4^+^ subset that decreased with HFD (**Fig S13F**). In addition to Vγ1/4^-^ expansion, HFD altered this subset’s phenotype by increasing the percentage of Vγ1/4^-^ cells that expressed IL17a/a and CCR6 (**Fig S13G-H**). Lastly, HFD significantly increased the relative contribution of γδ T cells to the overall IL17a production by T cells (**Fig S13I**), emphasizing their unique reactivity with HFD and active immunomodulatory role within the fibrotic implant environment.

Dysregulated wound healing and increased tissue fibrosis are further exasperated with aging^96^. Age-related factors such as ECM component modifications (e.g., glycosylation and crosslinking) and cellular senescence contribute to matrix accumulation and stiffening^96^. Murine γδ T cell responses also change across lifespan with progressive accumulation of γδ17 observed in numerous tissues including adipose, lymph nodes, lung, intestinal lamina propria, and injured muscle treated with pro-regenerative biomaterials^5,63,97–100^. We performed VML injuries with PCL implants in young 8 wk (standard), 16 wk, and early middle-aged 35 wk old mice to assess if age alters γδ T cell implant responses (**Fig S14A**). γδ T cells composed a significantly greater percentage of T cells in the PCL environment of middle-aged mice compared to young controls at 6 wks (**Fig S14B**). The distribution of Vγ chains shifted in middle-aged mice with a significant increase in Vγ1/4^-^ and decrease in Vγ4^+^ percentage (**Fig S14C**). Further, middle-aged mice had a significantly higher percentage and cell count of γδ17 compared to young controls (young: 64.1±15.6%, middle-aged: 88.3±9.6%) (**Fig S14D-E**). Collectively, γδ T cell responses within the fibrotic PCL environment can be affected by host factors that are independent of the local wound and implant properties.

### γδ17 stimulate fibroblast expression of pro-inflammatory factors and ECM components

Given γδ T cells that persist at chronic implant time points predominantly display a γδ17 effector phenotype, we investigated whether γδ17 can interact with fibroblasts to affect implant-associated fibrosis. We generated a CD45^+^-enriched scRNAseq dataset that captured both immune and stromal cell fractions at the 6 wk PCL implant time point (18,201 cells, n=3) (**Fig S15A-B**). The fibroblast cluster expressed genes related to IL17 signaling^101^ including IL17R subunits (*Il17ra* and *Il17rc*), adaptor proteins (*Traf3ip2* encoding Act1, *Traf6*, *Map3k7*), TFs and regulators (*Nfkb1* encoding NFκβ, *Nfkbiz*, *Mapk1*, *Cebpb*), and downstream targets (*Il6*) (**Fig 4A**, **S15C**). Additionally, the fibroblast cluster exhibited a significant positive correlation (R=0.36) between *Il17ra* expression and an ‘IL17 Response Score’ generated with UCell^102^ using the Gene Ontology (GO)^103,104^ ‘Response to Interleukin-17’ gene set further supporting that fibroblasts in the fibrotic implant environment actively engage in IL17-mediated signaling (**S15D**).

**Fig 4.**
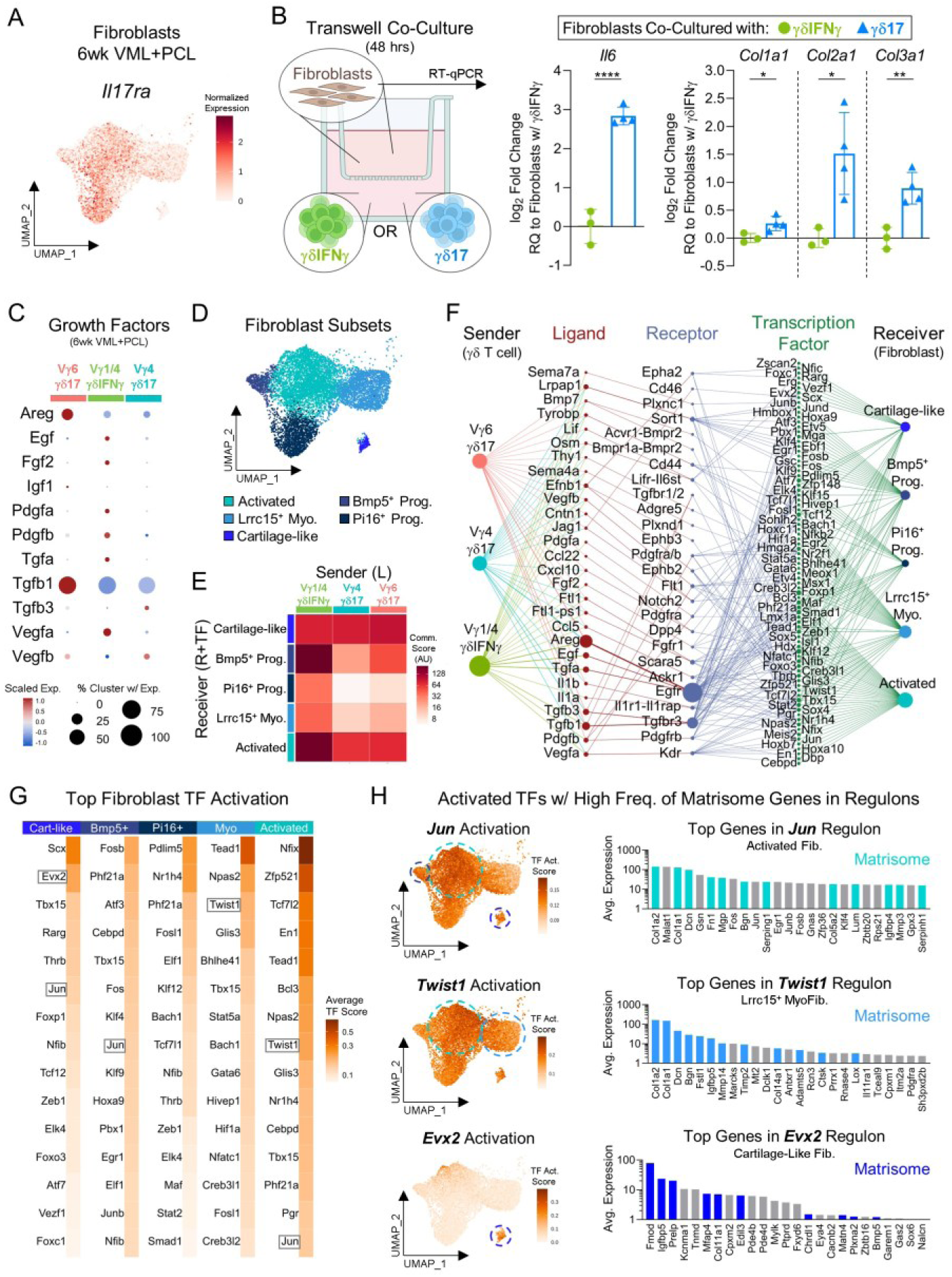
Signaling from γδ T cells can promote fibroblast expression of ECM-related genes. **(A)** Feature plot displaying *Il17ra* normalized expression in fibroblast clusters from a 6 wk VML+PCL CD45^+^-enriched scRNAseq dataset (n=3). **(B)** Schematic of *in vitro* co-culture assay of primary murine fibroblasts (50,000 cells) seeded in Transwell and phenotypically skewed γδ T cells (γδIFNγ or γδ17, 200,000 cells) in plate wells (left). Pro-inflammatory (*Il6*) and collagen (*Col1a1*, *Col2a1*, and *Col3a1*) gene expression in fibroblasts after 48-hour co-culture with γδ17 relative to γδIFNγ using RT-qPCR (right). **(C)** Row-scaled expression of a curated list of secreted growth factors by scRNAseq γδ T cell clusters in VML+PCL at 6 wks. **(D)** UMAP of 5 fibroblast clusters (10,085 cells) in VML+PCL at 6 wks. **(E)** Global communication score heatmap of signaling from “sender” γδ T cell clusters (L: ligand) to “receiver” fibroblast clusters (R: Receptor, TF: Transcription Factor) in VML+PCL at 6 wks. **(F)** Inferred dominoSignal network connecting γδ T cell ligands to fibroblast receptors and activated TFs in VML+PCL at 6 wks (L and R average expression >0.01, TF activation scores >0.05). **(G)** Top 15 activated TFs present in γδ T cell signaling network per fibroblast cluster ranked by average TF activation score. Gray boxes indicate TFs highlighted in H. **(H)** Select activated fibroblast TFs (*Jun*, *Twist1*, and *Evx2*) with high frequency of matrisome-related genes in regulon. Feature plot of TF activation score (left) and average expression of top 25 genes present in respective TF regulon (right) with colored bars indicating matrisome association. **(Statistics)** Bar graphs: mean±SD. Data analyzed using unpaired two-tailed student t-test (B). Communication and TF activation scores are presented on log scale (E, G, and H). NS: Not significant p>0.05, * p<0.05, ** p<0.01, *** p<0.001, **** p<0.0001.

To interrogate this potential interaction, we established an *in vitro* Transwell co-culture system with murine dermal fibroblasts and splenic γδ T cells. The γδ T cells were skewed via plate-bound αCD3 with soluble IL2 for γδIFNγ or soluble IL1β, IL7, IL21 and IL23 for γδ17 for 72 hours prior to the 2-day co-culture in basal media. Enzyme-linked immunosorbent assay (ELISA) for IFNγ and IL17a/a performed on the co-culture media confirmed cytokine secretion by the skewed γδ T cells (**Fig S16A**). Co-culture with γδ17 significantly increased fibroblast expression of inflammatory genes *Il6* and *Ptgs2* (encodes cyclooxygenase-2, an enzyme involved in prostaglandin synthesis) (**Fig 4B**, **S16B**). Notably, γδ17 co-culture also stimulated fibroblasts to upregulate expression of ECM components including collagens (*Col1a1*, *Col2a1*, *Col3a1*) and proteoglycans (*Prg4* encoding lubricin), suggesting that γδ17 may modulate ECM composition through their interactions with fibroblasts (**Fig 4B**, **S16C**).

Next, we introduced an IL17a neutralizing antibody (αIL17a) or isotype control into the co-culture media to isolate the effects of IL17a-mediated signaling. Expectedly, addition of αIL17a resulted in decreased fibroblast expression of IL17 signaling-related genes (*Nfkbiz*, *Il6, Ccl2*)^105^ (**Fig S16D**). Interestingly, fibroblast expression of *Col3a1* also significantly decreased with αIL17a, however expression of other ECM components (*Col1a1, Col2a1*, *Col5a1*, *Col9a2*, *Acan*, *Prg4*) was not significantly changed (**Fig S16E-F**). These results indicate that while γδ17 can modulate fibroblast gene expression of ECM components that contribute to tissue fibrosis, these transcriptional changes are only partly mediated by IL17a signaling and motivate investigation of other γδ17 secreted factors.

### γδ T cells contribute to activation of fibroblast transcription factors involved ECM remodeling

Growth factors can stimulate fibroblast proliferation, collagen production, and tissue remodeling. At the fibrotic 6 wk PCL time point, γδ T cells expressed a variety of growth factors including ones belonging to the epidermal (EGF), fibroblast (FGF), insulin-like (IGF), platelet-derived (PDGF), transforming (TGF), and vascular endothelial (VEGF) growth factor families, albeit some were detected at low abundance in only a subset of cells (**Fig 4C**, **S17A**). We leveraged our two scRNAseq datasets to further explore potential interactions between γδ T cells and fibroblasts beyond IL17-mediated signaling. First, computational sub-clustering of the heterogeneous fibroblast population was performed to improve resolution and revealed 6 distinct fibroblast subsets: an activated subset (*Sfrp1, Sfrp2, Mmp3*)^106,107^, a *Lrrc15*^+^ myofibroblast subset (*Lrrc15, Tnc, Acta2*)^108^, a cartilage-like subset (*Fmod, Tnmd, Comp*), and 2 progenitor-like subsets (*Bmp4, Bmp5, Penk; Pi16, Dpp4, Igfbp5*)^108–110^, in addition to a small proliferating subset (*Mki67, Top2a*) that was excluded (**Fig 4D, S17B**). All of the fibroblast subsets ubiquitously expressed common ECM components including collagens (*Col1a1*, *Col1a2*, *Col3a1*, *Col5a1*), fibronectin (*Fn1*), and proteoglycans (*Dcn* encoding decorin) (**Fig S17C**), suggesting they actively contribute to and shape the matrix composition. However, only the cartilage-like and a small portion of the activated fibroblast subsets expressed prototypical cartilage ECM components including *Col2a1, Col9a1*, *Col11a1*, *Prg4*, *Comp* (encodes cartilage oligomeric matrix protein), *Acan* (encodes aggrecan), and *Fmod* (encodes fibromodulin) (**Fig S17D**).

Next, we identified putative interactions between γδ T cells and fibroblasts in the 6 wk PCL implant environment by applying dominoSignal^111,112^, an algorithm that predicts intercellular communication networks. These are networks are constructed based on ligand expression in “sender” clusters and corresponding receptor expression with correlated downstream TF activation, which is inferred using the SCENIC algorithm, in “receiver” clusters. The global signaling heatmap reflects the intensity of potential signaling interactions between the “sender” γδ T cells and “receiver” fibroblasts by integrating ligand, receptor and TF activity (**Fig 4E**). The network revealed specific signaling pathways that originate from γδ T cells and activate fibroblasts, with the number of interactions indicated by node size and edge thickness (**Fig 4F**, **S18A**). The Vγ6 γδ17 IFN-Stimulated subset was omitted from this intercellular communication analysis due to low cell count at 6 wks. We identified multiple growth factor-related networks with predicted signaling mediated through EGF, FGF, PDGF, TGF, and VEGF receptors. For instance, *Tgfb1* and *Tgfb3* expressed by γδ T cells primarily signaled through *Tgfbr3* expressed by fibroblasts and predominantly activated TFs in the cartilage-like fibroblasts (**Fig S19A-B**). While *Pdgfa* and *Pdgfb* signaling through *Pdgfra* and *Pdgfrb* activated the most TFs in the *Lrrc15*^+^ myofibroblasts (**Fig S19C-D**), *Areg* and *Egf* signaling through *Egfr* activated many TFs across all fibroblast subsets (**Fig S19E-F**).

We ranked fibroblast TFs involved in the inferred γδ T cell communication network by calculating the mean TF activation score by cluster to discern which TFs appear to have the greatest downstream activity (**Fig 4G**). Several highly activated TFs across the various fibroblast subsets regulate tissue development and pro-fibrotic fibroblast differentiation. For example, *Tead1* was one of the highest activated TFs in the *Lrrc15*^+^ myofibroblast and activated fibroblast subsets with signaling networks mediated by various growth factors (*Areg, Egf, Tgfa, Vegfa, Vegfb*) (**Fig S20A**). The TEAD TF family plays an important role in embryonic development^113^, however it is also involved in YAP/TAZ mechanotransduction and recently implicated in driving pathological cardiac, hepatic, and renal fibrosis^114–116^. The most activated TF in the activated fibroblasts was *Nfix* (**Fig 4G**, **S20B**), which critically controls fetal myogenesis^117^ as well as post-natal muscle regeneration after injury^118^. The TF scleraxis (*Scx*), an important developmental regulator of tendon formation^119,120^ and modulator of fibrotic tendon healing^121^, was the most activated TF in the cartilage-like fibroblasts (**Fig 4G**, **S20C**). Beyond its role in tendons, scleraxis was identified as a distinct TF expressed by myofibroblasts in lung fibrosis^122^ and found to contribute to cardiac fibrosis by driving cardiac fibroblast conversion into myofibroblasts^123,124^. Further, scleraxis is a proposed biomarker of fibrotic disease severity in patients with idiopathic pulmonary fibrosis (IPF) and systemic sclerosis^125^. The predicted γδ T cell signaling was also associated with activation of cartilage-like fibroblast TFs important for cartilage remodeling, including ones implicated in cartilage matrix degeneration (*Rarg*, *Tcf12*, *Aft, Zeb1, Jun, Foxc1*)^126–130^ as well as cartilage protection (*Foxo3*)^131,132^.

Lastly, we screened the inferred regulons of activated TFs for overlap with the matrisome^133^ to assess whether γδ T cell signaling may affect fibroblast expression of ECM components linked to tissue remodeling (**Fig S18B**). The TF *Jun*, commonly associated with tissue fibrosis^134–136^, was activated in three fibroblast subsets with many matrisome-related genes present in its regulon including ECM proteins (*Col1a2*, *Col1a1*, *Col5a2*, *Fn1*), proteoglycans (*Dcn*, *Bgn*, *Lum*), and matrix metalloproteinases (*Mmp3*) (**Fig 4H**, **S21A-C**). Similarly, many top genes in the *Twist1* regulon, which was a TF activated in *Lrrc15*^+^ myofibroblasts and activated fibroblasts, were associated with ECM components (*Col1a2*, *Col1a1*, *Dcn*, *Bgn*) and enzymatic matrix modulators (*Mmp14*, *Timp2*, *Ctsk*, *Lox*) (**Fig 4H**, **S21D-F**). Twist1 contribution to epithelial-mesenchymal transition and fibroblast activation implicates it in driving progression of fibrotic diseases^137–141^. Within the cartilage-like fibroblasts, the *Evx2* TF regulon consisted of genes related to cartilage and connective tissues (*Fmod*, *Col11a1*, *Prelp*, *Mfap4*, *Edil3*, *Matn4*) (**Fig 4H**, **S21G-I**). Collectively, signaling from γδ T cells is predicted to activate TFs in the fibroblast populations that are linked to tissue development and remodeling processes involved in fibrosis.

### γδ T cells modulate fibroblast phenotype and ECM composition in implant-associated fibrosis

Since our findings suggest that γδ T cells may regulate fibroblast expression of matrisome-related genes, we assessed their functional contribution to the tissue fibrosis by implanting PCL into muscle defects in γδ T cell-deficient mice (TCRδ^-/-^, B6.129P2-*TCRd*^tm1Mom^/J). Bulk RNAseq analysis of FACS-isolated stromal cells (Live CD45^-^ CD29^+^ CD31^-^) harvested from C57BL/6J wildtype (WT) and TCRδ^-/-^ mice 6 wks after PCL implantation revealed 365 significant differentially expressed genes, many of which corresponded to ECM components (**Fig 5A**). Specifically, stromal cells from TCRδ^-/-^ mice significantly increased expression of numerous collagens (*Col1a1*, *Col3a1*, *Col5a1*, *Col5a2*, *Col5a3*) and ECM glycoproteins (*Prg4, Comp, Acan*, *Vcan, Fbn2*, *Tnc*) (**Fig 5B**). Interestingly, TCRδ^-/-^ stromal cells only significantly reduced expression of two collagens, *Col9a2* and *Col11a2*, which are commonly associated with collagen II primarily found in articular cartilage^142^. Immunofluorescent staining of collagen XI revealed that it localized to the fibrotic tissue surrounding PCL particles and was less abundant in TCRδ^-/-^ mice upon visual inspection (**Fig 5C**, **S22**). Further, the TCRδ^-/-^ stromal cells significantly decreased expression of skeletal muscle-related genes including fast-twitch troponins (*Tnnc2*, *Tnni2*, *Tnnt3*), myosin (*Myh2*, *Myh4*, *Mylpf*), and skeletal alpha-actin (*Acta1b*) (**Fig 5B**).

**Fig 5.**
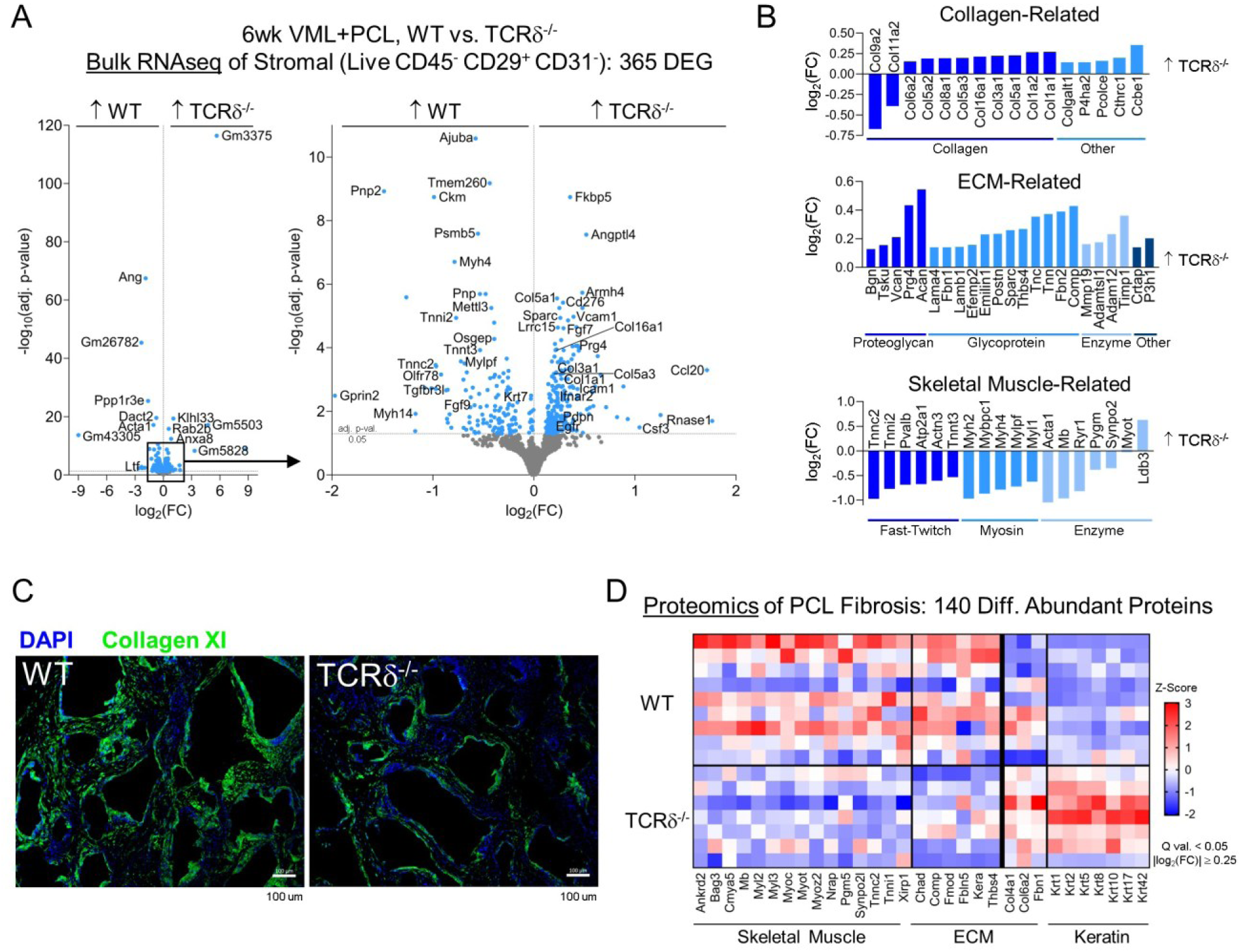
γδ T cells may influence the composition of peri-implant fibrotic tissue. **(A)** Differential gene expression of FACS-isolated stromal cells (Live CD45^-^ CD29^+^ CD31^-^) from VML+PCL at 6 wks in C57BL/6 wildtype (WT) versus TCRδ^-/-^ mice using bulk RNA sequencing. **(B)** Log fold change of significant differentially expressed genes (adj. p-val. <0.05) related to collagen, ECM, and skeletal muscle group by manual curation. **(C)** Representative immunofluorescence staining of collagen XI in fibrotic tissue surrounding PCL particulate implants in WT and TCRδ^-/-^ mice (scale bar: 100μm). **(D)** Z-score heatmap of select significant differentially abundant proteins (Q val. <0.05, |log_2_(FC)| ≥0.25) from PCL-associated fibrotic tissue in WT (n=9) versus TCRδ^-/-^ (n=7) mice using LC-MS/MS.

To explore whether these transcriptional changes correspond to protein-level changes in matrix composition, we performed quantitative proteomics using liquid chromatography-tandem mass spectrometry (LC-MS/MS) on fresh-frozen fibrotic PCL implants from WT and TCRδ^-/-^ mice harvested at 6 wks. We detected 6,446 proteins including 20 collagens (I, VI, XIV being most abundant) and numerous ECM glycoproteins (fibronectin, decorin, lumican, thrombospondin, versican) (**Fig S23A**). A total of 140 proteins met the criteria for significant differential abundance and most were decreased in fibrotic tissue of TCRδ^-/-^ mice (**Fig 5D**, **S23B**). As observed on the transcriptional level, many skeletal muscle-related proteins such as myosin light chains, troponins, and myoglobin were less abundant in the fibrotic tissue of TCRδ^-/-^ mice. In contrast, the fibrotic tissue of TCRδ^-/-^ mice had significantly greater amounts of type I and II keratins that form intermediate filaments, which serve as important cytoskeletal components for cell structure, migration, and mechanical stress responses. The fibrotic tissue of WT mice contained more ECM proteins associated with cartilaginous connective tissue including COMP, chondroadherin (CHAD), FMOD, and fibulin-5 (FBLN5, involved in elastic fiber formation), whereas the fibrotic tissue of TCRδ^-/-^ mice was enriched in collagen VI, collagen IV, and fibrillin-1 (FBN1).

### γδ T cells signal to endothelial cells and affect blood vessel size in implant-associated fibrosis

The molecular composition and scaffolding of ECM can regulate many aspects of tissue structure, including vascular development and network organization. Fibrotic tissues often display aberrant vascular remodeling and EC dysfunction that can contribute to fibrotic pathophysiology^143,144^. We indeed observed increased expression of ECM components related to vessel wall structure and elasticity (collagens I, III, VI, VIII, fibrillins, emilin) as well as vascular basement membrane (collagen IV, laminins) in TCRδ^-/-^ mice that motivated further vascular profiling (**Fig 5A-D**). The TCRδ^-/-^ mice had significantly more CD31^+^ ECs compared to WT controls by flow cytometry and immunofluorescent staining (**Fig 6A-B, S24A-D**). Quantitative histological evaluation of blood vessel size revealed that TCRδ^-/-^ mice had significantly more large blood vessels (luminal surface area >3000 μm^2^) in the fibrotic PCL environment compared to WT controls (**Fig S24E**). These results suggest that γδ T cells contribute to shaping the vascular architecture in fibrosis by modulating the size and number of blood vessels.

**Fig 6.**
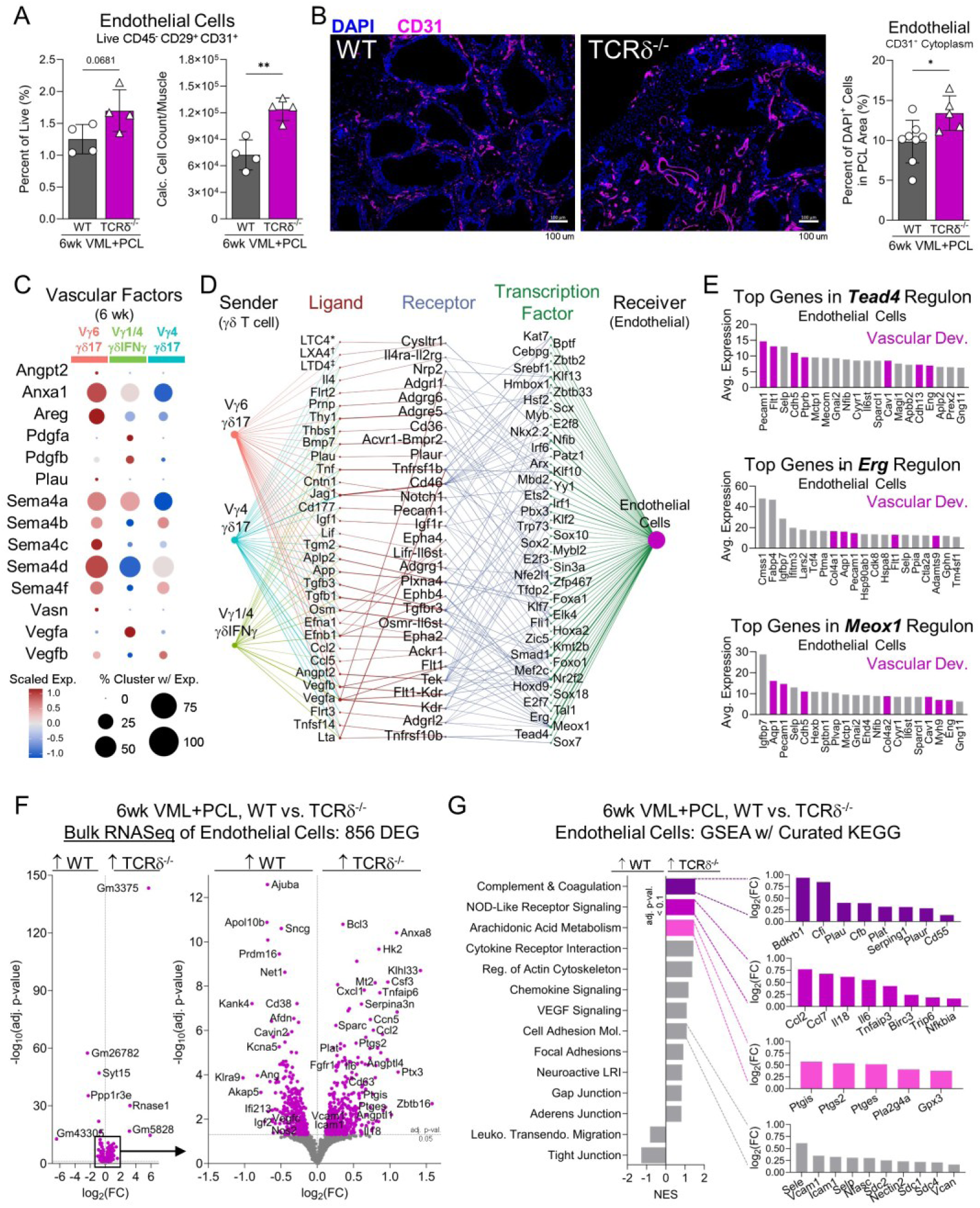
γδ T cells communicate with endothelial cells to modulate vascularity of peri-implant fibrotic tissue. **(A)** Flow cytometric quantification of endothelial cells (ECs) (Live CD45^-^ CD29^+^ CD31^+^) from VML+PCL at 6 wks in C57BL/6 wildtype (WT) versus TCRδ^-/-^ mice. **(B)** Representative immunofluorescence staining (left) and histological quantification (right) of CD31^+^ ECs in fibrotic tissue surrounding PCL in WT and TCRδ^-/-^ mice (scale bar: 100μm). **(C)** Row-scaled expression of a curated list of secreted vascular factors by scRNAseq γδ T cell clusters in VML+PCL at 6 wks. **(D)** Inferred dominoSignal network connecting γδ T cell ligands to EC receptors and activated TFs in VML+PCL at 6 wks (L and R average expression >0.01, TF activation scores >0.05). **(E)** Average expression of top 20 genes in regulons of select EC activated TFs (*Tead4*, *Erg*, and *Meox1*) with colored bars indicating vascular development association. **(F)** Differential gene expression of FACS-isolated ECs from VML+PCL at 6 wks in WT versus TCRδ^-/-^ mice using bulk RNA sequencing. **(G)** Gene set enrichment analysis (GSEA) of WT versus TCRδ^-/-^ ECs using a curated subset of vascular-related KEGG gene sets. Colored bars indicate significant pathways (adj. p <0.1) with expression of significant leading edge genes (adj. p <0.05). NES: normalized enrichment score. **(Statistics)** Bar graphs: mean±SD. Data analyzed using unpaired two-tailed student t-test (A and B). NS: Not significant p>0.05, * p<0.05, ** p<0.01.

Since γδ T cells affected the number of ECs and blood vessel size, we investigated whether they interact with ECs and impact EC phenotype in fibrosis. γδ T cells in the PCL environment expressed secreted factors that can influence EC behavior and vessel properties including amphiregulin (*Areg*), angiopoietin-2 (*Angpt2*), annexin1 (*Anxa1*), pro-inflammatory cytokines, urokinase-type plasminogen activator (*Plau*), semaphorins, vasocrin (*Vasn*), and VEGFs (**Fig 6C**, **S25A**). The γδIFNγ subset had the highest scaled expression of *Vegfa*, which is a central angiogenic factor that activates EC migration and proliferation, whereas the two γδ17 subsets had greater expression of *Vegfb*, whose role in angiogenesis is less understood but is implicated maintenance and survival of new blood vessels^145,146^. We applied dominoSignal^111,112^ to identify putative interactions in which ligands expressed by γδ T cells were linked to receptors and activated TFs in the EC cluster derived from the CD45^+^-enriched scRNAseq dataset (**Fig 6D**, **S25B**). This revealed that γδ T cells may interact with ECs via adhesion molecules (*Cd177-Pecam1*) and pro-inflammatory factors such as TNF family members (*Tnf*, *Lta*, *Tnfsf14*) and IL6 family members (*Osm*, *Lif*). γδ T cell expression of *Vegfa* and *Vegfb* is inferred to activate many EC TFs by signaling through VEGFR-1 (*Flt1*), VEGFR-2 (*Flt1-Kdr* complex) and neuropilin (*Nrp2)*. Additionally, γδ T cell expression of leukotrienes (*LTC4*, *LTD4*) may stimulate EC activation of *Nfib* by *Cysltr1*. Leukotrienes are arachidonic acid derivates involved in increasing vascular permeability, vasoconstriction, and leukocyte adhesion to blood vessels^147–149^.

γδ T cell signaling was predicted to activate EC TFs that regulate vessel properties and contain genes in their regulons associated with vascular development (**Fig 6E**, **S25C**). For example, TEAD TF family members form complexes with YAP/TAZ to facilitate EC nutrient acquisition that supports angiogenesis (**Fig 6E, S25D**)^150^, while the TF *Erg* promotes VEGF-induced angiogenesis and vascular stability (**Fig 6E, S25E**)^151,152^. Krüppel-like factors (*Klf2*, *Klf7, Klf10, Klf13*) also play important roles in vascular biology and their dysregulation is associated with various cardiovascular pathologies^153–157^. The TF *Meox1,* also activated in the fibroblast clusters and increasingly implicated in fibrotic pathologies^158–160^, is involved in modulating vascular smooth muscle cell phenotype to control vascular tone and remodeling (**Fig 6E, S25F**)^161^. Collectively, these results support that γδ T cells may signal to and influence EC behavior, ultimately impacting vascular network properties in fibrotic environments.

To determine how γδ T cell interactions with ECs may functionally alter blood vessel properties and tissue vascularity, we performed bulk RNAseq on FACS-isolated ECs (Live CD45^-^ CD29^+^ CD31^+^) from WT and TCRδ^-/-^ mice 6 wks after PCL implantation that resulted in 856 significant differentially expressed genes (**Fig 6F**). GSEA with a collection of KEGG gene sets relevant to vasculature found that ECs from TCRδ^-/-^ mice are enriched in pathways related to innate immunity such as ‘complement and coagulation’ and ‘NOD-like receptor signaling’ (**Fig 6G**). In regard to inflammation, TCRδ^-/-^ ECs increased expression of numerous genes related to neutrophil and monocyte chemoattractants including *Il34*, *Cxcl1*, *Ccl2*, *Ccl7*, and *Csf3*. TCRδ^-/-^ ECs also displayed significantly higher expression of adhesion molecules involved in leukocyte attachment and transendothelial migration, namely *Sele* (encodes E-selectin), *Selp* (encodes P-selectin), *Icam1*, and *Vcam1*. However, flow cytometric immune cell profiling did not reveal an increase in the counts or frequency of myeloid subsets (Ly6G^+^ neutrophils, Ly6C^hi^ monocytes, or F4/80^+^ macrophages) within the 6 wk PCL environment of TCRδ^-/-^ mice compared to WT controls (**Fig S26**). Additionally, the TCRδ^-/-^ ECs were enriched in the ‘arachidonic acid metabolism’ pathway with significantly higher expression of prostaglandin synthases and receptors genes (*Ptgir*, *Ptges*, *Ptgs2, Ptgis*) (**Fig 6G**). Prostaglandins are another type of arachidonic acid derivatives implicated in promoting inflammation, vessel permeability, and vasodilation^162,163^. In contrast, TCRδ^-/-^ ECs had significantly lower expression of *Edn1*, which encodes the potent vasoconstrictor endothelin. These findings suggest that γδ T cells play a functional role in modulating EC response to inflammation and shaping the vasculature within fibrotic tissues.

## Discussion

γδ T cells are increasingly recognized for their involvement in regulating tissue physiology, including reparative and fibrotic outcomes following tissue injury^1^. Their capacity to broadly survey the environment and mount rapid responses with rich bioactive secretomes and myriad effector functions supports further investigation of how γδ T cells interact with neighboring stromal cells to affect tissue structure. Here, we modeled tissue fibrosis by implanting synthetic biomaterials, primarily PCL particulate, into critical-sized murine muscle defects. In addition to serving as a robust system for studying mechanisms underlying pathological fibrosis, this model also recapitulates clinically relevant features of biomedical implants, including implantation in the presence of injury and implant interfacing with multiple tissue types (muscle, adipose, and skin). Biomedical implants are widely used for medical and cosmetic indications; however, their susceptibility to fibrotic encapsulation often limits clinical efficacy by causing implant malfunction that can pose patient safety risk and require invasive removal. While prior studies detect γδ T cells in fibrotic tissue surrounding biomaterial implants in clinical and preclinical contexts^49,51,164^, their functional role in the biomaterial implant-associated fibrosis remains largely unexplored.

By coupling scRNA/TCRseq and high-parameter flow cytometry, we identified multiple γδ T cell effector subsets, specifically γδIFNγ and several γδ17, that are present in fibrotic tissue responses induced by injury and PCL implants. The two main γδ17 subsets were distinguished by their Vγ chains, (semi)invariant δTCR sequences, unique transcriptional profiles, and biased expression of IL17a versus IL17f. As previously described^73^, the Vγ6^+^ γδ17 is tissue-resident, while the Vγ4^+^ γδ17 migrated to the inflamed implant site from secondary lymphoid organs. The identified Vγ6^+^ γδ17 appears comparable to the main γδ T cell population present in the early inflammatory response to cardiotoxin-induced muscle injury^28^. Moreover, this γδ T cell characterization aligns with, and considerably expands upon, our prior work that broadly identified IFNγ and IL17 producing γδ T cells in the FBR^49^. Both γδIFNγ and γδ17 expanded during the acute inflammatory phase of wound healing and were drastically increased with PCL implantation relative to injury only controls. However, the two subsets displayed divergent kinetics with γδIFNγ contracting and γδ17 persisting as fibrosis progressed at longer time points. These persistent γδ17 were a prominent producer of IL17a relative to other T cell subtypes, supporting that their high bioactivity may influence the fibrotic PCL environment.

Tissue-resident γδ17 expanded in the fibrotic PCL environment following a HFD regimen and as mice reached middle-age (35 wks), revealing that γδ T cells change in response to environmental stimuli and host conditions. These results align with prior studies that find elevated γδ17 with HFD and old age across numerous tissue types (e.g., adipose, liver, skin, and LNs)^63,94,97,100,165,166^. While γδ T cells increase by middle adulthood^5,63,99^, future studies with PCL implants should include old or geriatric mice to adequately capture age-related tissue stiffening and fibrosis. Notably, old mice (72 wks) have greater γδ T cell expression of IL17a than young controls (6 wks), which contributed to age-related immune dysfunction and impaired muscle repair following treatment with pro-regenerative biological scaffolds^100^. The impact of diet and aging on γδ T cells presents a potential mechanism that may contribute to the observed heterogeneity in patient responses to biomedical implants. We investigated body weight and age due to their known positive correlation with fibrotic pathologies^89,96,167,168^; however, other host factors may also impact γδ T cell responses during tissue fibrosis. For instance, manipulation of the gut microbiome using germ-free conditions, broad-spectrum antibiotics depletion, and enteric infection-induced dysbiosis is shown to perturb γδ T cell responses to cardiotoxin-injured muscles and biomaterial implants^28,51^.

Since γδ T cells respond to an extensive array of antigen types, future work must be done to elucidate what they are sensing in fibrotic implant environments, as well as in HFD and aging conditions, that drives their activation. A hypothesis that warrants further consideration are signals produced by senescent cells (SnCs), which are cells marked by proliferative growth arrest that accumulate with fibrosis^49,169^, HFD-induced obesity^170,171^, and aging^172^. SnCs secrete bioactive lipids as part of their inflammatory senescence associated secretory phenotype (SASP) that may be sensed by γδ17, which can uptake lipids and utilize oxidative metabolism^93^. SnCs also upregulate stress-like molecules, including MICA and ULBP2, that can be recognized by killer cell lectin-like receptors expressed by NK and γδ T cells^173–175^. Further, SnCs upregulate transcription of endogenous retroviruses (ERVs) that stimulate anti-viral IFN responses^176–178^. Interestingly, HFD is found to potentiate γδ17 and promote skin inflammation to commensal microbiota by heightening keratinocyte ERV expression and cGAS/STING/IFN signaling^165^. γδ17 may interact in a similar positive feed-forward manner with SnCs as is observed with T_H_17 cells in fibrotic conditions^49,179^.

The promotion of collagen expression by fibroblasts co-cultured with γδ17 and computational interactions linking putative γδ T cell ligands to activation of fibroblast TFs involved in ECM remodeling suggest that the persistent γδ17 may play an active role in shaping peri-implant fibrosis. This complements prior co-culture studies that found human γδ T cells isolated from patients with systemic sclerosis significantly increase fibroblast *COL1A2* expression relative to ones isolated from healthy controls^180^. We perturbed IL17 signaling using neutralizing antibodies; similar mechanistic studies should be performed to investigate how other γδ T cell secreted ligands (Amphiregulin, TNFα, TGFβ) impact fibroblast expression of matrix components. To further capture γδ T cell-to-fibroblast interactions, intercellular communication analysis performed on our 6wk VML+PCL scRNAseq datasets predicted signaling (*Areg*-*Egfr* and *Tgfb1/3*-*Tgfbr3*) and fibroblast TF activation (*Jun*, *Tead1*, *Twist1*, *Nfix*, *Scx*) commonly observed in tissue development and fibrotic pathologies. These interactions may have valuable therapeutic implications for mitigating implant-associated fibrosis. Notably, inhibition of TGFβ3^181–183^, TWIST1^140,184,185^, and YAP/TAZ-TEAD^186–188^ are being actively investigated as cancer and fibrotic disease treatments.

The absence of γδ T cells during fibrosis progression altered not only fibroblast expression of prevalent ECM collagens and glycoproteins, but also, surprisingly, prototypical cartilage matrix components. The physiological role of cartilage-like ECM and the biological processes that stimulate its production in fibrotic contexts are still largely unknown; however, cartilage exhibits unique properties (e.g., viscoelasticity and avascularity) that may substantially impact the fibrotic microenvironment. Emerging evidence suggests that “cartilage-like” fibroblasts, similar to those identified in our scRNAseq dataset, may be present in other fibrotic pathologies. For instance, myofibroblasts in fibrotic scars of porcine split-thickness skin grafts upregulated *COL11A1*, *ACAN*, *COMP*, and a ‘chondrocyte differentiation’ signature in response to the high mechanical stresses^189^. Moreover, mesenchymal fibroblasts in human keloids differentially expressed *COL11A1*, *COMP*, and *ASPN*^190^. The expression of these canonical cartilage genes is also reported in liver fibrosis^191^, IPF^192,193^, fibrotic skin diseases^194^, abdominal adhesions^195^, and pancreatic fibrosis^196^. The cartilage-like tissue detected in the PCL environment may be comparable to synovial metaplasia or lubricin-containing pseudomembranes that form around biomedical implants (e.g., breast tissue expanders, meshes, and orthopedic joint protheses)^197–200^. The TCRδ^-/-^ mice also had decreased expression of skeletal muscle components at mRNA and protein levels, suggesting γδ T cell involvement in regulating muscle repair. Similarly, ablation of γδ T cells in a cardiotoxin muscle injury model reduced proliferation of muscle precursor cells, decreased expression of a ‘filament structure/contractile function’ gene signature, and delayed muscle regeneration^28^.

TCRδ^-/-^ mice demonstrated an altered EC phenotype and blood vessel structure within the fibrotic PCL environment. This expands upon prior findings that γδ T cells support angiogenesis^11–13^, enhance vascular permeability to promote acute inflammation during wound healing^28^, and influence EC behavior to modulate vascular tone^201,202^. Computationally predicted intercellular communication revealed numerous signaling networks by which γδ T cells may interact with ECs to influence downstream vascular development including pro-inflammatory cytokines, pro/anti-angiogenic factors, and leukotrienes. Since sufficient tissue perfusion is essential for effective repair and often dysregulated in fibrotic contexts, a continued investigation of how γδ T cells impact EC behavior and vascular properties (e.g., sprouting, permeability, maturity, 3D network structure, and contractility) is warranted.

We acknowledge that compensatory responses by innate-like lymphocytes and CD4^+^ T cells can confound interpretation of results obtained using the TCRδ^-/-^ strain, thus motivating follow-up validation using conditional γδ T cell depletion models (e.g., knock-in diphtheria toxin strains^69^ and anti-γδTCR) or selective Vγ knockout strains. Additionally, murine γδ T cells are recognized to be considerably different than human γδ T cells, including their development and effector phenotypes^203,204^; thus, future work is needed to discern how these findings translate to human fibrotic pathologies. In all, our data suggest that γδ T cells and their diverse secretome can actively shape the composition and vascularity of fibrotic implant environments by interacting with resident stromal cells, ultimately emphasizing the critical role of immunoregulation in tissue fibrosis.

## Materials and Methods

### Mice

All animal procedures were performed in accordance with Johns Hopkins University (JHU) Animal Care and Use Committee (ACUC) approved protocols MO21M80/MO24M66. Mice were housed in JHU animal facilities under standard conditions and sterile facility feed (except for special diet studies described below). All strains were obtained directly from Jackson: Wildtype (C57BL/6J, JAX stock #000664) and TCRδ^-/-^ (B6.129P2-*TCRd*^tm1Mom^/J, JAX stock #002120)^205^. All experiments were performed using female mice at 7-10 wks of age. The aging study also included 16- and 35-wk old wildtype, female mice.

### Volumetric Muscle Loss (VML) Injury with Biomaterial Implants

Mice were fully anesthetized (1.5-3% isoflurane with 100% O_2_), the hindlimb fur shaved, and the surgical site thoroughly disinfected. Carprofen (5mg/kg) (Rimadyl 54771-8507-1 or OstiFen 510510) was administered via subcutaneous injection for analgesia. VML injury was performed as previously described^49,77,206^. In short, longitudinal incisions were made in the skin and fascia overlaying the quadriceps femoris muscle. A critical-sized defect (3 mm x 3 mm x 3 mm) was excised from the quadriceps muscle using sterile micro dissecting scissors. The defect was directly filled with either sterile Dulbecco’s phosphate-buffered saline (PBS, no implant control), polycaprolactone particulate (PCL, MW 50,000, <600 μm mean particle size) (Polysciences 25090), polyethylene powder (PE, ultra-high MW, 40-80 μm particle size) (Sigma Aldrich 434272), poly(dimethylsiloxane) (PDMS/silicone, viscosity 18,000-22,000 cST) (Sigma-Aldrich 432997), or decellularized porcine urinary bladder matrix (ECM, reconstituted at 200 mg/mL in dPBS) (Integra, MicroMatrix MM1000). The PBS, silicone and ECM were administered through a syringe (50 μL/defect), while the PCL and PE were placed using a sterile surgical spoon (1 scoop/defect) (spoon dimensions: 8 mm diameter, 1.5 mm depth) (Roboz RS-6161 or Moria Surgical 1121b). The skin incision was closed using sterile veterinary nylon sutures. The VML injury with material implantation was repeated on the other hindlimb to complete the bilateral procedure. Mice were placed on a heat pad and closely monitored to ensure full recovery.

### High Fat Diet (HFD) Experiments

After a 1-2 wk acclimatization period to the JHU animal facility following delivery, mice were weighed (baseline) and randomly placed on either an *ad libitum* high-fat diet (HFD, 60 kcal% Fat) (Research Diets D12492i, irradiated) or low-fat diet (LFD, 10 kcal% Fat) (Research Diets D12450Ji, irradiated) for 8 total wks. Mice were weighed and feed was replenished weekly. After 2 wks on the new diets, mice received bilateral VML injuries with PCL implants as described above. Feed was replenished and mice body weight was measured on a weekly basis. Injured mice were then maintained on LFD/HFD regimens for 6 wks prior to euthanasia and tissue harvest for immunophenotyping using techniques described below.

### Inhibition of T Cell Egress from Lymph Nodes (LNs) with Fingolimod

T cell egress from LNs was pharmacologically blocked using fingolimod hydrochloride (FTY720 HCl) (Selleckchem S5002) delivered via intraperitoneal injection at 25 μg/mouse prepared in sterile saline (Sodium Chloride 0.9%) (Intermountain Life Sciences Z1376). Sterile saline served as the corresponding vehicle control. C57BL/6J wildtype mice received bilateral VML injures with PCL implants and randomized between vehicle and FTY720 groups. FTY720 was administered on the day of surgery (day 0), again on day 1, and then every 2-3 days (3 doses per week) for the duration of the 6 wk long experiment. T cell sequestration was confirmed via flow cytometric (Cytek Aurora) profiling of peripheral blood using αCD45, αCD11b, αCD3, and αTCRγδ fluorophore-conjugated antibodies at the 6 wk VML+PCL harvest time point.

### Tissue Processing for Flow Cytometry and Sequencing

#### Muscles with Biomaterial Implants

Following euthanasia, VML-injured quadricep muscles along with any implanted material were carefully harvested. The tissues were mechanically dissociated by manual dicing, then enzymatically digested using Liberase TL (0.5 mg/mL) (Roche 05401020001) and DNAse I (0.2 mg/mL) (Roche 10104159001) in RMPI-1640 with L-Glutamine (Gibco 11875-093) and 25 mM HEPES (Quality Biological 118-089-721) at 37°C for 45 mins under agitation. Enzymatic digestion was quenched using RMPI-1640 supplemented with 1% w/v bovine serum albumin (BSA) (Sigma-Aldrich A9647) and placing samples on ice. Digested samples were filtered through 70 μm cell strainers (Falcon 352350) followed by either 40 μm cell strainers (Falcon 352340) or a Percoll density gradient (GE Healthcare 17-0891-01) for lymphocyte enrichment.

#### Inguinal Lymph Nodes (iLNs)

Following euthanasia, iLNs were carefully separated from inguinal fat pads and manually mashed through 70 μm cell strainers (Miltenyi Biotec 130-110-916) with blunt pestles. Cell strainers were washed with RMPI-1640 containing L-Glutamine and HEPES.

#### Blood

Mice were anesthetized using isoflurane and peripheral blood drawn from the submandibular vein was collected into K_2_EDTA-coated tubes (BD 365974) to prevent clotting. A standardized blood volume of 150 μL was analyzed for all samples. Red blood cell lysis was performed with 1x Pharm Lyse Lysing Buffer (BD 555899) and washed with dPBS.

### Multi-Parameter Flow Cytometry

#### Viability and Surface Staining

Single-cell suspensions from digested and/or filtered tissues were plated into 96-well U-bottom plates. After pelleting, cells were washed with dPBS prior to viability staining with either Zombie NIR Fixable Viability kit (BioLegend 423106) or LIVE/DEAD Fixable Aqua Dead Cell Stain kit (ThermoFisher Scientific L34957) for 30 mins covered on ice. Cells were then washed with FACS buffer consisting of dPBS (Gibco 14190-144) supplemented with 1% w/v BSA and 1mM EDTA (Invitrogen 15575-038) prior to surface marker staining for 45 mins covered on ice (**Mat. Tables 1-5**). All surface marker antibodies were prepared in FACS buffer with TruStain FcX (1:20) (BioLegend 101320), True-Stain Monocyte Blocker (1:50) (BioLegend 426102), and Super Bright Complete Staining Buffer (1:50) (eBioscience SB-4401-75). Cells were washed with FACS buffer and immediately sorted (for bulk or single-cell RNAseq) or fixed using FluoroFix Buffer (BioLegend 422101) for 15 mins covered at room temperature.

#### *Ex Vivo* Stimulation and Intracellular Cytokine Staining (ICS)

Lymphocyte-enriched single-cell suspensions were plated into 96-well U-bottom plates. Cells were incubated in 1x eBioscience Cell Stimulation Cocktail Plus Protein Transport Inhibitors (eBioscience 00-4975-93) diluted in Iscove’s Modified Dulbecco’s Media (IMDM) with L-Glutamine and 25mM HEPES (Gibco 21056-023) supplemented with 10% v/v heat-inactivated fetal bovine serum (Gibco 26140-079) for 3.5 hrs at 37°C with 5% CO_2_. After pelleting, cells were washed with dPBS prior to viability and surface staining as detailed above. Stained cells were fixed and permeabilized using Cyto-Fast Fix/Perm solution (BioLegend 426803) for 20 mins covered at room temperature, followed by washing with 1x Cyto-Fast Perm Wash solution, and intracellular cytokine staining (ICS) for 20 mins covered at room temperature (**Mat. Table 3**). All ICS antibodies were prepared in 1x Cyto-Fast Perm Wash solution with TruStain FcX (1:20), True-Stain Monocyte Blocker (1:50), and Super Bright Complete Staining Buffer (1:50). In addition to single stains and fluorescence minus one (FMO) controls, isotype controls were prepared for all ICS antibodies. Cells were washed with 1x Cyto-Fast Perm Wash solution prior to storage in FACS buffer.

#### Flow Cytometry and Data Analysis

All panels were run on the spectral 4-laser Cytek Aurora flow cytometer (Lasers: 405 nm Violet, 488 nm Blue, 561 nm Yellow/Green, and 640 nm Red) with automated sample loader. Cytek Aurora data acquisition and spectral unmixing was performed using SpectroFlo software. Autofluorescence signatures were extracted from unstained samples. Manual flow gating to identify cell populations of interest was performed using FlowJo software (Tree Star, v10.8) (**Suppl Fig 27-29**). Gate placement for cell population delineation was guided by FMO and isotype controls. Samples for different timepoints in the time course analysis (**Fig 2E**) were processed and acquired on different days, but spectrally unmixed and gated together. Calculated cell counts were determined using the following formula: CalcCount=(TotalVol/AcqVol)*(DilFac)*(Count) where “TotalVol” was total sample volume, “AcqVol” was volume of sample acquired as recorded by the cytometer’s volumetric measurement, “DilFac” was the dilution factor by which the sample was split prior to staining, and “Count” was the number of events recorded by cytometer within the desired cell population gate. To visualize γδ T cells subsets using high-parameter flow data (**Fig 2D**), all γδ T cells across independent samples and timepoints were selected using manual gating and concatenated into a single .fsc file, followed by dimensional reduction performed using the FlowJo Uniform Manifold Approximation and Projection (UMAP) plugin^207^. Median fluorescence intensity (MFI) for cytokine expression was measured considering only cells in the positive gate. All data is presented as mean±SD. Significant outliers based on Vγ chain distribution were identified using ROUT method (Q=0.2%) and removed from subsequent analysis in HFD experiment. For experiments with 2 groups, data was analyzed using an unpaired, two-tailed student t-test. For experiments with > 2 groups, data was analyzed with either one-way or two-way ANOVA (specified in corresponding figure captions) with Tukey’s multiple comparison test. All statistical analysis and data graphing were performed in GraphPad Prism (v9.0.0 - v10.3.0).

### Single-Cell RNA/TCR Sequencing

Sample Preparation CD3^+^ T Cell Dataset: Data for scRNA/TCRseq was collected in 3 independent experiments: (1) 1 wk VML+Saline (n=18, pooled) and 1 wk VML+PCL (n=2, pooled), (2) 1 wk VML+PCL (n=5, hashed) and 12 wk VML+PCL (n=5, hashed), and (3) 6 wk VML+PCL (n=3, hashed). Quadricep muscles, and fibrotic PCL implants if present, were harvested at the abovementioned time points and processed as described above. After lymphocyte enrichment with a Percoll density gradient, samples were stained for viability and surface markers (**Mat. Table 4**) as described above. During the last 30 minutes of surface staining, hashed samples had TotalSeq-C antibodies (**Mat. Table 4**) added to their staining mixture. Cell sorting was performed by the JHU SKCCC High Parameter Flow Core on a 4-laser BD FACSAria Fusion Cell Sorter. T cells (LiveSSC^lo^CD45^+^CD3^+^) were sorted into dPBS with 1% BSA, centrifuged, and re-suspended in dPBS with 0.01% BSA. Sorted T cells were counted using the Countess (Thermo Fisher Scientific) and processed for 5’ paired scRNA/TCRseq with feature barcodes (10x Genomics, v1.1 chemistry) by the JHU Transcriptomics & Deep Sequencing Core. Library preparations for gene expression, feature barcodes, and αβTCRs were performed using standard Chromium Single Cell 5’ library Construction kit v1.1 (10x Genomics PN 1000020) as per manufacturer’s protocols. Library preparation for γδTCRs was performed using previously published^208^ custom primers (**Mat. Table 6**) (Integrated DNA Technologies) and methods. Briefly, the standard VDJ protocol (10x Genomics v1.1) was followed until section 4.0, at which point the following custom T cell mixes were used: “T cell mix 1” (10X-FP-1 and GDM-RP-1-#) and “T cell mix 2” (10X-FP-2 and GDM-RP-2-#), with only 12 cycles of amplification per thermocycler step. Libraries were sequenced using the NovaSeq platform (Illumina) with S2 100 flow cell at a target depth of 100,000 unique reads/cell.

Sample Preparation CD45^+^ Enriched Dataset: Data for CD45^+^-enriched scRNAseq was collected at 6 wk VML+PCL (n=3, each replicate run on independent lane). Quadricep muscles and fibrotic PCL implants were harvested at the 6 wk time point and processed as described above (no density gradient). Dead cells were removed with Magnetic-Activated Cell Sorting (MACS) using the Dead Cell Removal kit (Miltenyi Biotec 130-090-101). MACS with the mouse CD45 MicroBeads (Miltenyi Biotec 130-052-301) was used to separate and collect the CD45^+^ immune cells (positive fraction) and CD45^-^ stromal cells (negative fraction). Cells were counted in both fractions using a hemocytometer and pooled at in a 1:1 ratio. The CD45^+^-enriched samples were processed for 3’ scRNAseq (10x Genomics, v3.1 chemistry) by the JHU Transcriptomics & Deep Sequencing Core. Libraries were sequenced using the NovaSeq platform (Illumina) with S2 100 flow cell at a target depth of 100,000 unique reads/cell.

### Computational Data Analysis of Single-Cell RNA/TCR Sequencing

#### Data Pre-Processing

FASTQ files were aligned to GENCODE vM23 annotations and genome using Cell Ranger (10x Genomics, v6.1.2) with default parameters^209^. To demultiplex the hashed samples in the CD3^+^ T cell data and pair both αβ TCR sequencing and γδ TCR sequencing with the aligned gene expression, the multi-function was used along with hashtag oligo sequences to separate samples, after which the bamtofastq function was used to create sample specific FASTQ files. The multi-function was run once more to align the αβ TCR sequences, then the VDJ function was used to additionally align the γδ TCR sequences.

#### Quality Control

Automatically called cells were further filtered to isolate high quality captured cells and remove empty droplets. Quality control parameters were selected separately for analysis of the CD45^+^ enriched data, CD3^+^ T cell data, and subset γδ T cell data based on cells with a unique molecular identifier (UMI) count above a threshold, a feature count above a threshold, and less than a specified percentage of mitochondrial genes (%MT). For CD45^+^ enriched data: UMI >500, feature count >350, and %MT < 25%. For CD3^+^ T cell data: UMI >2000, feature count >1000, and %MT <5%. For γδ T cell data: UMI >1000, feature count >750, and %MT <10%. Only features detected in at least 0.1% of cells were included in downstream analyses.

#### TCR Analysis

The paired scTCRseq data was integrated with gene expression data using scRepertoire (v2.0.0)^210^. Based on the associated CDR3 sequence for a cell, an α, β, γ, and/or δ chain was identified. Following selection of the γδ T cell population (see below), clones were identified using the amino acid sequences of the γ and δ chains, requiring a match in both chains for a clone to be called. To compare TCR sequences to known (semi)invariant sequences, cells with the correct δ chain, regardless of γ chain status, were selected.

#### γδ T Cell Selection

γδ T cells were identified using both gene expression and TCR data. Cells exhibiting gene expression consistent with the regular expression patterns “^Tr[gd]v” or “^TCR[gd]-V\\d” without expression of genes matching the pattern “^Tr[ab]v”, were classified as γδ T cells based on gene expression. Genes matching “^Tr[ab]v.dv” were disregarded due to the inability to discern recombination with α or δ components. TCR-based γδ T cell selection required a γ or δ chain sequence associated with a cell without α or β chain sequences. Following subset isolation, processing and analysis steps were performed. After the initial clustering analysis of the γδ T cells, 4 clusters of contamination/doublets, cycling, or low-quality cells were removed before re-processing the remaining γδ T cells.

#### Data Processing

Seurat (v4.4.0) was used for normalization, identification of highly variable genes, scaling (including regressing out the percentage of mitochondrial genes and total UMI count), principal component analysis, dimensionality reduction via UMAP, shared nearest neighbor graph construction, and graph-based community detection using the Louvain (CD3^+^ T cell data, γδ T cell data) or Leiden (CD45^+^ enriched data) algorithms for clustering^207,211–215^. An elbow plot guided the selection of the number of principal components for downstream analyses or corrected principal components for analyses that were batch corrected to reduce technical artifacts. Data pertaining to CD3^+^ T cells or the subset γδ T cells underwent batch correction using Harmony (v1.2.0)^216^, based on sample collection group.

Cluster resolutions were determined by considering silhouette scores, canonical marker gene expression, cluster stability visualized using Clustree (v0.5.1), and differential expression analysis conducted with the Wilcoxon rank-sum test implemented in Presto (v1.0.0)^217,218^. Unless otherwise indicated, differential expression analysis was performed by comparing each cluster against all remaining cells. For further analysis, clusters annotated as fibroblasts from the CD45^+^ enriched data were combined and isolated for dedicated investigation, using the same analysis approach described.

#### Gene Set Enrichment Analysis (GSEA)

Ranked GSEA using the Wilcoxon rank-sum differential expression results ranked by -log_10_(p-adj)*sign(logFoldChange) was performed using the fgsea package^219^. Gene ontology biological processes^103,104^ and Hallmark^220^ gene sets were sourced from MSigDB. Pathways with p-adj <0.05 were determined to be significantly enriched.

#### IL17 Signaling Correlation

IL17 response scores were generated using UCell (v2.6.2)^102^ with a modified version of Gene Ontology “Response to Interleukin-17” gene set (GO:0097396) acquired from the Molecular Signatures Database. The only modification of the gene set was removal of *Il17ra*. We used Pearson correlation to evaluate the relationship between expression of *Il17ra* with the IL17 response scores in the fibroblast cluster with statistical significance evaluated with cor.test in R.

#### Secreted Ligand Selection

Secreted ligands were identified using the protein annotations provided with CellPhoneDB (v4.1.0)^221^ which were subsequently matched to their associated genes. Gene orthologs between provided human gene symbols and mouse gene symbols were determined using Ensembl 111, GRCh38.p14^222^. The resulting murine genes were considered secreted ligands and subsequently visualized.

#### Transcription Factor (TF) Activity Inference

TF regulons and activity scores were inferred using the pySCENIC (v0.12.1) implementation of the SCENIC algorithm^223^. ScRNAseq data were converted into loom files using loomR (v0.2.1.9) and processed with SCENIC using the command line interface, with a seed of 42 set where possible and using the mm10 genome reference provided with SCENIC. The AUC scores output by SCENIC were used for TF activity, and expression of genes within the regulon was calculated based on normalized gene expression within the scRNAseq data. Matrisome-associated genes were identified by selecting those present within the matrisome gene sets^224^. Genes associated with vascular development were selected from the gene ontology biological process gene sets^103^ containing “angiogenesis,” “venous_blood,” “vasodilation,” “vascular_wound_healing,” or “blood_vessel” in their names.

#### Intercellular Communication Analysis

Cell-cell communication inference was conducted with dominoSignal (v0.99) (see data and code availability statement) utilizing CellPhoneDB ligand-receptor pairings, SCENIC regulon and TF activity scores, and the gene expression data and cluster assignments from analysis of each dataset^111,112^. Parameters included enabling complexes, p-adj <0.001 to determine TF inclusion, a minimum receptor to TF correlation of 0.25, a minimum receptor expression threshold of 10% within the relevant cell population, and no limitations on TF or receptor numbers. Analyses were performed independently for each 6 wk time point sample. Pathways consistently identified across all samples within each analysis data set (CD45^+^ enriched data or subset γδ T cell data) were retained after merging. Ligand and receptor pairings from CellPhoneDB linked populations in the CD45^+^ enriched data to populations within the subset γδ T cell data to create a network with connections between the separate datasets. To maintain robust connections, networks were filtered to include only receptor-TF correlations >0.25 with a p-value <0.05 calculated for each cluster. To enhance the identification of biologically relevant signaling pathways, thresholds were set for network connections based on ligand and receptor expression levels, as well as TF activation. Average ligand and receptor normalized expression levels were >0.01, while TF activation score was >0.05. Network plots include nodes and edges weighted by the number of pathways in which they participate. A global communication score was developed to quantify interaction strength. Ligand expression, receptor expression, and TF activation were each scaled within the network to range from 1 to 10 using the scales package (v1.3.0)^225^. These scaled values were multiplied to equally weigh each pathway component. For legibility, the resulting scores were adjusted by order of magnitude as the units are arbitrary.

#### Data Visualization

Experiment schematics were created with BioRender. Volcano plots for differential gene expression and GSEA plots were created in GraphPad Prism (v9.0.0 - v10.3.0). Heatmaps were created using ComplexHeatmap (v2.14.0)^226,227^ or ggplot2 from the tidyverse (v2.0.0)^228^. Network communication plots were created in igraph^229,230^. Additional plotting tools such as ggnewscale (v0.4.10)^231^, ggrepel (v0.9.5)^232^, data manipulation tools such as stringdist (v0.9.12)^233^, reshape2 (v1.4.4)^234^, plyr (v1.8.9)^235^, and graphics device svglite (v2.1.3)^236^ were also used for data visualization.

### Bulk RNA Sequencing

#### Sample Preparation

PCL was implanted into bilateral hindlimb VML defects in wildtype and TCRδ^-/-^ female mice. After 6 wks, mice were euthanized and quadriceps muscles with fibrotic PCL implants were harvested and processed as outlined above (excluding Percoll density gradient). Single-cell suspensions were stained for viability and surface markers (**Mat. Table 5**). To achieve sufficient cell yields, tissues sourced from 2 independent mice (4 total quads) were pooled together prior to sorting to form a “biological replicate”, resulting in 4 “biological replicates” per experimental group. Cell sorting was performed by the JHU SKCCC High Parameter Flow Core on a 4-laser BD FACSAria Fusion Cell Sorter. Stromal cells (LiveCD45^-^CD31^-^ CD29^+^) and endothelial cells (LiveCD45^-^CD31^+^CD29^+^) were sorted directly into DNA LoBind Eppendorf tubes (Eppendorf 022431048) filled with sterile RLT Plus Buffer (Qiagen 1030963) and 1x 2-Mercaptoethanol (Gibco 21985023). After sorting, tubes were vortexed and left at room temperature for 1-2 mins to allow for cell lysis, followed by snap freezing on dry ice and short-term storage at -80°C.

#### Library Prep and Bulk RNAseq

All RNA isolation, QC, library prep and bulk RNAseq was performed by the JHU Transcriptomics & Deep Sequencing Core. RNA was isolated using miRNeasy Mini kit (Qiagen 217004). RNA yield was determined using NanoDrop Spectrophotometer and quality assessed using Agilent Fragment Analyzer (RQN for all samples between 8.7-9.6). cDNA synthesis and library preparation with unique dual indexes (UDI) was performed using Stranded mRNA Prep, Ligation kit (Illumina 20040532). QC was assessed with Qubit and Agilent Fragment Analyzer. Libraries were pooled and pair-end sequencing performed using the NovaSeq platform (Illumina) with a SP100 flow cell at a targeted depth of 50 million unique reads/sample.

#### Data Analysis

FASTQ files were aligned to the GENCODE vM23 mouse genome and transcriptome annotation using STAR aligner (v2.7.84)^209,237^. Gene expression quantification was performed concurrently with alignment using STAR’s built-in gene counting feature. Counts associated with the gene *Trdc* were removed from further downstream analysis due to constitutive, unproductive overexpression in the knock-out model. Differential gene expression analysis was conducted with DESeq2^238^, incorporating a design matrix that accounted for mouse strain. Statistical significance for differential gene expression was determined using a false discovery rate corrected threshold of p-adj <0.05. GSEA was performed with fgsea (v1.25.1), ranking genes by “sign(log_2_FoldChange) * −log_10_(p-adj)” using a curated subset of the KEGG gene sets^239^, selecting for pathways pertaining to endothelial cells and/or vessels. Pathways with a corrected p-adj. *<*0.1 were determined to be significantly enriched. Significant differentially expressed genes were manually grouped into biologically-relevant categories (e.g., collagen, ECM, and skeletal muscle) based on literature review. Volcano plots for differential gene expression and GSEA plots were created in GraphPad Prism (v9.0.0 - v10.3.0).

### RT-qPCR

#### Sample Preparation

VML-injured muscles with fibrotic PCL implants were carefully excised and placed into RNA*later* Stabilization Solution (Invitrogen AM7024) for at least 48 hours at 4°C, then transferred into cold TRIzol Reagent (Invitrogen 15596018). Tissues were homogenized using the Bead Ruptor 12 (OMNI International 19-050A) with 2.8 mm ceramic beads (OMNI International 19-646) for 2-3 cycles (speed: 6 m/sec, cycle duration: 15 sec). RNA was purified using chloroform extraction followed by RNeasy Plus Mini kit (Qiagen 74134) as per manufacturer’s protocol. RNA concentration was measured using NanoDrop 2000 Spectrophotometer (ThermoFisher Scientific).

#### cDNA Preparation and RT-qPCR

cDNA was synthesized using SuperScript IV VILO Master Mix (Invitrogen 11756500) and a C1000 Touch Thermocycler (BioRad). RT-qPCR was performed with 100ng cDNA/well using TaqMan Gene Expression Master Mix (Applied Biosystems 4370074) and TaqMan probes (single-plex FAM) (**Mat. Table 7**) following manufacturer’s protocol on the StepOne Plus Real-Time PCR System (Applied Biosystems). *Rer1* was selected as the endogenous reference control (housekeeper gene) based on its stability across experimental groups.

#### Data Analysis

All RT-qPCR data was analyzed using the StepOne software (Applied Biosystems, v2.3). All samples for all primers were required to have a cycle threshold (CT) below the pre-determined cut-off value of 35. For HFD experiments, samples were normalized to LFD control group. All RT-qPCR data was analyzed using the Livak Method for calculation of ΔΔCT values and fold change is 2^-ΔΔCT^. For statistical analysis, the fold change was transformed to log_2_(2^-ΔΔCT^) and presented as mean±SD on a linear scale. Data was analyzed using an unpaired, two-tailed student t-test. All statistical analysis and data graphing were performed in GraphPad Prism (v9.0.0 - v10.3.0).

### *In Vitro* Transwell Co-Culture

#### Murine Dermal Fibroblast Isolation and Culture

Dermal fibroblasts were isolated from 3 wk old wildtype C57BL/6J mice. After euthanasia, dorsal fur was removed by shaving and topical application of depilatory cream, followed by skin sterilization with 70% EtOH. A rectangular section of dorsal skin (approx. 1 cm x 2 cm) was harvested, mechanically dissociated by manual dicing, then enzymatically digested using Liberase TM (0.5 mg/mL) (Roche 5401127001) in RMPI-1640 with L-Glutamine (Gibco 11875-093) at 37°C for 45 mins under agitation. Enzymatic digestion was quenched using complete RPMI media, RMPI-1640 supplemented with 10% heat-inactivated fetal bovine serum FBS (Gibco 26140-079), 1% Penicillin-Streptomycin (Pen-Strep) (Gibo 15140-122), and 1% Sodium Pyruvate (Gibco 11360-070). Digested tissue fragments were centrifuged, media exchanged to fresh complete RPMI, tissue fragments transferred into tissue culture (TC) treated T-175 flasks, and incubated at 37°C with 5% CO_2_ for 4 days. On day 5, the adherent cells were washed with sterile PBS and culture media changed to complete MEM media, MEM (Gibco 11095080) supplemented with 10% heat-inactivated FBS and 1% Pen-Strep to selectively culture dermal fibroblasts. The dermal fibroblasts were passaged at 80% confluency in complete MEM media for 1-3 passages usage for Transwell co-culture assay.

#### Murine γδ T Cell Isolation and *In Vitro* Cytokine Skewing

γδ T cells were isolated from spleens of 12 wk old wildtype C57BL/6J mice. After euthanasia, spleens were harvested and manually mashed through 70 μm cell strainers (Falcon 352350) with blunt pestles. Red blood cell lysis was performed with 1x Pharm Lyse Lysing Buffer (BD 555899) and followed by filtration through 40 μm cell strainers (Falcon 352340). γδ T cells were purified from the single-cell suspension using MACS with the mouse TCRγ/δ+ T Cell Isolation Kit (Miltenyi Biotec 130-092-125) as per manufacturer protocol. γδ T cell purity was confirmed using flow cytometry. γδ T cells were cultured in T cell media consisting of RPMI-1640 supplemented with 10% heat-inactivated FBS, 1% Pen-Strep, 1% Sodium Pyruvate, 1% Non-Essential Amino Acids (Gibco 11140-050), and 1% HEPES. γδ T cells were seeded at 200,000 – 300,000 cells/well in a 96-well TC-treated flat bottom plate in T cell media and skewed *in vitro* using plate-bound anti-CD3 antibody (10 μg/mL) (BioLegend 100238) and either IL2 (10 ng/mL) (BioLegend 575402) for γδIFNγ or IL1β (BioLegend 575102), IL7 (BioLegend 577802), IL21 (BioLegend 574502), and IL23 (10 ng/mL each) (BioLegend 589002) for γδ17 (adapted from ^93,240,241^). γδ T cells were incubated in respective cytokine cocktail at 37°C with 5% CO_2_ for 72 hours and phenotypic skewing was confirmed via enzyme-linked immunosorbent assay (ELISA) for IFNγ (Invitrogen 88-7314) and IL17A homodimer (Invitrogen 88-7371) on supernatant as per manufacturer’s protocol.

#### Transwell Co-Culture Assay

Isolated dermal fibroblasts (50,000 cells/Transwell) were plated onto the 6.5 mm Transwell with 0.4 μm Pore Polyester Membrane Insert (Corning 3470) in complete MEM media overnight to allow for cell adhesion to the membrane. On day 0 of the co-culture, γδIFNγ or γδ17 (200,000 cells/well) were plated in the bottom well in fresh T cell media. For experiments that blocked IL17-mediated signaling, anti-mouse IL17A monoclonal antibody (70 ng/mL) (clone eBioMM17F3) (Invitrogen 16-7173-81) or corresponding mouse IgG1 kappa Isotype control (70 ng/mL) (clone P3.6.2.8.1) (Invitrogen 16-4714-82) was added to the co-culture media. The co-culture was incubated at 37°C with 5% CO_2_ for 48 hours. On day 2, the co-culture media was collected for quantification of by γδ T cells production of IFNγ and IL17A homodimer using ELISA. The fibroblasts were lysed directly in the Transwell using RLT Plus Buffer with 1x 2-Mercaptoethanol for RNA isolation and transcriptional profiling via RT-qPCR.

#### RNA Isolation and RT-qPCR of Co-Culture Fibroblasts

RNA was purified using RNeasy Plus Micro kit (Qiagen 74034) as per manufacturer’s protocol. RNA concentration was measured using NanoDrop 2000 Spectrophotometer (ThermoFisher Scientific). cDNA was synthesized as described above. cDNA was further pre-amplified using TaqMan PreAmp Master Mix (Applied Biosystems 4391128) for 10 cycles as per manufacturer’s protocol. RT-qPCR was performed on the pre-amplified cDNA as described above using TaqMan probes (**Mat. Table 7**).

#### Data Analysis

All RT-qPCR data was analyzed as described above. For γδIFNγ versus γδ17 co-culture experiments, samples were normalized to fibroblasts co-cultured with γδIFNγ. For IL17 neutralization co-culture experiments, samples were normalized to fibroblasts co-cultured with γδ17 and isotype control. For ELISA, absorbances were measured at 450 nm and 570 nm using Varioskan Lux Multimode Microplate Reader (ThermoFisher Scientific) and the difference calculated as per manufacturer’s protocol. The standard curve was generated using a four-parameter logistic curve fitting. Sample concentrations were calculated using the generated standard curve. In certain experimental conditions and samples were below the limit of detection (n.d.), which prohibited statistical analysis. Data graphing was performed in GraphPad Prism (v9.0.0 - v10.3.0).

### Histology

#### Sample Preparation

VML-injured muscles with fibrotic PCL implants were carefully excised and fixed for 48 hours in 10% neutral buffered formalin (Sigma-Aldrich HT501320) with agitation at room temperature. Tissues were washed with DPBS, dehydrated using a graded series of ethanol (EtOH) washes, transferred to Xylenes (ThermoFisher Scientific X3P-1GAL), and stored in paraffin wax overnight at 55-60°C. Tissues were bisected, embedded in paraffin blocks, and sectioned (7 μm) with a microtome in-house or at the JHU Oncology Tissue Services SKCCC core facility.

#### Masson’s Trichrome Staining

Formalin-fixed paraffin embedded (FFPE) sections of VML-injured muscles with PCL implants were stained for Masson’s Trichrome by the JHU Oncology Tissue Services SKCCC core facility. Slides were imaged with NanoZoomer-XR digital slide scanner (Hamamatsu Photonics) and viewed using NDP.view2 software (Hamamatsu Photonics, v2.9.25).

#### Immunofluorescence Staining

FFPE sections (7 μm) of VML-injured muscles with PCL implants from wildtype C57BL/6J and TCRδ^-/-^ mice were baked at 55-60°C for 1 hour, deparaffinized with xylenes and serial EtOH washes, and rinsed with type I water. Antigen retrieval was performed by adding slides into heated 1x AR6 Buffer (Akoya Biosciences AR600250ML) for 15 minutes in a steamer. The slides were cooled to room temperature, rinsed with type 1 water, and endogenous peroxidases quenched using fresh 3% H_2_O_2_ for 15 minutes. After blocking (10% BSA and 0.05% Tween20 in PBS) for 30 minutes, slides were stained with unconjugated primary antibodies: rabbit anti-mouse CD31 (1:2000) (Abcam 182981) or rabbit anti-mouse Col11a1 (1:100) (Gifted from Mei Sun at USF) for 30 minutes. Primary delete slides (i.e. slides not stained with primary antibody) served as negative controls. Slides were washed with 1x Tris Buffered Saline with Tween20 (TBST) (Cell Signaling Technology 9997S) and incubated with corresponding Rabbit-on-Rodent HRP-Polymer (Biocare Medical RMR622H) for 30 minutes. Slides were washed with 1x TBST and incubated in Opal570 Reagent (1:150) (Akoya Biosciences FP1488001KT) or Opal650 Reagent (1:150) (Akoya Biosciences FP1496001KT) in 1x Plus Amplification Diluent (Akoya Biosciences FP1498) for 10 minutes in the dark. Slides were washed with 1x TBST, rinsed with type 1 water, stained with 1x spectral DAPI (Akoya Biosciences FP1490) for 5 minutes, and rinsed with type 1 water. DAKO Fluorescence Mounting Medium (Agilent S302380-2) and cover slips were applied and slides dried overnight in the dark. Slides were imaged with Axio Imager A2 (Zeiss) and processed with ZEN software. All imaging parameters were kept consistent across groups and samples. The brightness of all presented IF images was increased by 40% in PowerPoint to aid visualization in final figures.

#### Image Analysis

The whole tissue sections were analyzed using HALO software (v3.6.4134) with cell segmentation performed using DAPI signal. The primary delete slides (missing incubation of primary antibody for marker of interest) were used as a negative control to set positive signal thresholding. After cell segmentation and annotation, the percentage and counts of CD31^+^ endothelial cells (ECs) were determined per fibrotic implant area analyzed. The QuPath image analysis platform (v0.5.1) was used to quantify the number and lumen size (surface area) of CD31^+^ blood vessels throughout the fibrotic implant area of a whole tissue section^242^. A threshold of > 500 μm^2^ lumen surface area was set to eliminate small cell clusters that also express CD31. All data is presented as mean±SD and analyzed using an unpaired, two-tailed student t-test. All statistical analysis and data graphing were performed in GraphPad Prism (v9.0.0 - v10.3.0).

### Proteomics via Liquid Chromatography-Tandem Mass Spectrometry (LC-MS/MS)

#### Sample Collection

Only fibrotic tissue encapsulating the PCL implant (i.e. no muscle tissue) was characterized using proteomics. The overlaying fibrotic tissue with PCL implant was carefully separated from the VML-injured muscle of wildtype C57BL/6J and TCRδ^-/-^ mice and immediately flash frozen.

#### Tissue Lysis and Homogenization

Samples were homogenized in 500 µL lysis buffer containing 8 M urea, 2% sodium dodecyl sulfate (SDS), 1 µM trichostatin A (TSA), 3 mM nicotinamide adenine dinucleotide (NAD), 75 mM sodium chloride, and 1X protease and phosphatase inhibitor cocktail (ThermoFisher Scientific) in 200 mM triethylammonium bicarbonate (TEAB) by adding them to 2.0 mL safe-lock tubes (VWR International) containing stainless steel beads and subjected to five intervals of high-speed shaking (25 Hz, 1 min) using a Qiagen TissueLyser II (Qiagen). Tissue homogenates were centrifuged at 15,700 x g for 10 min at 4°C, and the supernatant was collected for label-free quantitative proteomics experiments. Protein concentration was determined using Bicinchoninic Acid (BCA) assay (ThermoFisher Scientific).

#### Protein Digestion and Desalting

Aliquots of 100 µg protein lysates for each sample were brought to the same overall volume of 50 µL with water, reduced using 20 mM dithiothreitol in 50 mM TEAB at 50°C for 10 min, cooled to room temperature (RT) and held at RT for 10 min, and alkylated using 40 mM iodoacetamide in 50 mM TEAB at RT in the dark for 30 min. Samples were acidified with 12% phosphoric acid to obtain a final concentration of 1.2% phosphoric acid. S-Trap buffer consisting of 90% methanol in 100 mM TEAB at pH ∼7.1, was added and samples were loaded onto the S-Trap micro spin columns. The entire sample volume was spun through the S-Trap micro spin columns at 4,000 x g and RT, binding the proteins to the micro spin columns. Subsequently, S-Trap micro spin columns were washed twice with S-Trap buffer at 4,000 x g and RT and placed into clean elution tubes. Samples were incubated for one-hour at 47°C with sequencing grade trypsin (Promega) dissolved in 50 mM TEAB at a 1:25 (w/w) enzyme:protein ratio, and then digested overnight at 37°C. Peptides were sequentially eluted from micro S-Trap spin columns with 50 mM TEAB, 0.5% formic acid (FA) in water, and 50% acetonitrile (ACN) in 0.5% FA. After centrifugal evaporation, samples were resuspended in 0.2% FA in water and desalted with Oasis 10-mg Sorbent Cartridges (Waters). The desalted elutions were then subjected to an additional round of centrifugal evaporation and re-suspended in 0.1% FA in water at a final concentration of 1 µg/µL. Two microliters of each sample was diluted with 2% ACN in 0.1% FA to obtain a concentration of 20 ng/µL. One microliter of indexed Retention Time Standard (iRT, Biognosys) was added to each sample, thus bringing up the total volume to 21 µL^243^.

#### Mass Spectrometric Analysis

Samples (400 ng) were loaded onto EvoTips Pure (Evosep) following the manufacturer’s protocol. LC-MS/MS analyses were performed on an Evosep One liquid chromatography system (Evosep) coupled to a timsTOF HT mass spectrometer (Bruker). The solvent system consisted of 0.1% FA in water (solvent A) and 0.1% FA in ACN (solvent B). Peptides were eluted on a PepSep C18 analytical column (150 µm x 15 cm, 1.5 µm particle size; Bruker) using the 30 SPD method (44-minute gradient length, 500-nL/min flow rate). A zero-dead volume emitter was installed in the nano-electrospray source (CaptiveSpray source, Bruker Daltonics) and the source parameters were set as follows: capillary voltage 1600 V, dry gas 3 L/min, and dry temperature 180°C. Each sample was acquired in data-independent acquisition (DIA) mode coupled to parallel accumulation serial fragmentation (PASEF) or dia-PASEF mode with 1 survey TIMS MS1 scan and 24 dia-PASEF MS/MS scans per 2.03-s cycle^244^. The dual TIMS analyzer was operated in a 100% duty cycle with equal accumulation time and ramp time of 75 ms each. For each TIMS MS scan, the ion mobility ranged from 1/K0 = 0.85-1.30 Vs/cm^2^ and the mass range covered m/z 100-1,700. The dia-PASEF scheme was defined as 63 x 15 Th isolation windows from m/z 305.9 to 1,250.9 and covering the ion mobility range 1/K0 = 0.74-1.30 Vs/cm^2^ (**Supplementary Table S1**). The collision energy was defined as a linear function of mobility starting from 20 eV at 1/K0 = 0.6 Vs/cm2 to 59 eV at 1/K0 = 1.6 Vs/cm^2^. For calibration of ion mobility dimension, three ions of Agilent ESI-Low Tuning Mix ions were selected (m/z [Th], 1/K0 [Th]: 622.0289, 0.9915; 922.0097, 1.1986; 1221.9906, 1.13934).

#### Data Processing and Statistical Analysis

DIA data was processed in Spectronaut (v19.7.250203.62635) using directDIA. Data extraction parameters were set as dynamic and non-linear iRT calibration with precision iRT was selected. Data was searched against the *Mus musculus* reference proteome with 17,222 entries (UniProtKB, reviewed entries only), accessed on 03/27/2025. Trypsin/P was set as the digestion enzyme and two missed cleavages were allowed. Cysteine carbamidomethylation was set as a fixed modification while methionine oxidation and protein N-terminus acetylation were set as dynamic modifications. Identification was performed using 1% precursor and protein q-value. Quantification was based on the peak areas of extracted ion chromatograms (XICs) of 3 – 6 MS2 fragment ions, specifically b- and y-ions, with local normalization and q-value sparse data filtering applied (**Supplemental Table S2**). In addition, iRT profiling was selected. Differential protein expression analysis comparing TCRδ^-/-^ to WT was performed using an unpaired t-test and p-values were corrected for multiple testing using the Storey method^245^. Specifically, group wise testing corrections were applied to obtain q-values. Protein groups with at least two unique peptides, q-value < 0.05, and absolute log_2_(fold-change) > 0.25 are significantly-altered (**Supplemental Table S3**). A curated list of significant differentially abundant proteins was z-scored using z = (x-μ)/σ where x is the raw data, μ is population mean, and σ is population standard deviation for presentation on a heatmap. Volcano plots for differential protein expression and z-scored heatmaps were created in GraphPad Prism (v9.0.0 - v10.3.0).

## Data and Code Availability

### Single-cell RNA and TCR sequencing data

Raw FASTQ files and aligned counts files for both RNA and TCR sequences have been made available for download through NCBI GEO under the access number GSE 306147 and 306253. Processed objects are available upon reasonable request.

### Bulk RNA sequencing data

Raw FASTQ files and aligned counts files have been made available for download through NCBI GEO under the access number GSE 306146.

### Proteomics data

Raw data and complete MS data sets have been uploaded to the Mass Spectrometry Interactive Virtual Environment (MassIVE) repository, developed by the Center for Computational Mass Spectrometry at the University of California San Diego, and can be downloaded using the following link: https://massive.ucsd.edu/ProteoSAFe/private-dataset.jsp?task=6f0be9baa689478b8989ab9ec098f20b (MassIVE ID Number: MSV000098194; ProteomeXchange ID: PXD065015).

### Code

Scripts used for data analysis and visualization in this work were written in R (v4.2.1)^246^. Scripts for all analysis of scRNA/TCRseq and bulk RNAseq are publicly accessible in the GitHub repository hosted at https://www.github.com/Elisseeff-Lab/gd_t_cells_in_fibrosis. Communication inference analysis was performed using a pre-release branch of dominoSignal (v0.99) to implement experimental features not included in the package at the time (branch krishnan_analysis, commit: de6b93f). This version can be accessed from the dominoSignal Github repository at https://github.com/FertigLab/dominoSignal.

## Acknowledgements

We would like to thank the Johns Hopkins University (JHU) animal facilities and caretakers, the JHU SKCCC High Parameter Flow core facility for performing FACS, the JHU Transcriptomics & Deep Sequencing core facility for performing bulk and single-cell RNA sequencing, the JHU Oncology Tissue Services SKCCC core facility for histology sectioning and performing Masson’s trichrome staining, Maria Browne for scanning histology slides, Dr. Katlin Stiver for maintaining Cytek Auroa cytometer and providing flow cytometry expertise, and Dr. Mei Sun and Dr. David Birk for sharing the anti-mouse collagen XI histology antibody. This study was funded by the NIH Pioneer Award DP1AR076959 (J.H.E.), NSF Graduate Research Fellowship Program (A.R. and B.Y., DGE1746891), and K99AG081564 (J.C.M).

## Author Contributions

A.R., K.K., L.D.H., and J.H.E. conceptualized and designed the studies. A.R., J.W., J.C.M., H.N., C.A., C.D.K., and L.D.H. performed experiments and interpreted results. E.F.GG., D.R.M., A.N.R., and B.Y. assisted with experimental work. S.G., B.S., L.D.H., and J.H.E. advised on experimental work. K.K. performed computational analyses and interpreted results. C.C., M.P., F.H.Y., and E.J.F. assisted with and advised computational analysis. A.R., K.K., and J.H.E. wrote the manuscript. L.D.H., D.M.P., and J.H.E. supervised the study.

