## Supplementary Figures and Tables for "γδ17 T cell-stromal networks modulate matrix composition and vascularity in foreign body response"

[1] Translational Tissue Engineering Center, Department of Biomedical Engineering, Johns Hopkins School of Medicine, Baltimore, MD, USA
[2] Department of Biomedical Engineering, Duke University, Durham, NC, USA
[3] Buck Institute for Research on Aging, Novato, CA, USA
[4] Institute for Genome Sciences, Greenebaum Comprehensive Cancer Center, Department of Medicine, Department of Epidemiology, University of Maryland School of Medicine, Baltimore, MD, USA
[5] Department of Biomaterials & Applied Oral Sciences, School of Biomedical Engineering, Department of Microbiology & Immunology, Dalhousie University, Halifax, NS, Canada
[6] Bloomberg~Kimmel Institute for Cancer Immunotherapy, Sidney Kimmel Comprehensive Cancer Center, Johns Hopkins School of Medicine, Baltimore, MD, USA
[7] Department of Chemical Engineering, Johns Hopkins University, Baltimore, MD, USA
[8] Lead Contact

[+],^ These authors contributed equally

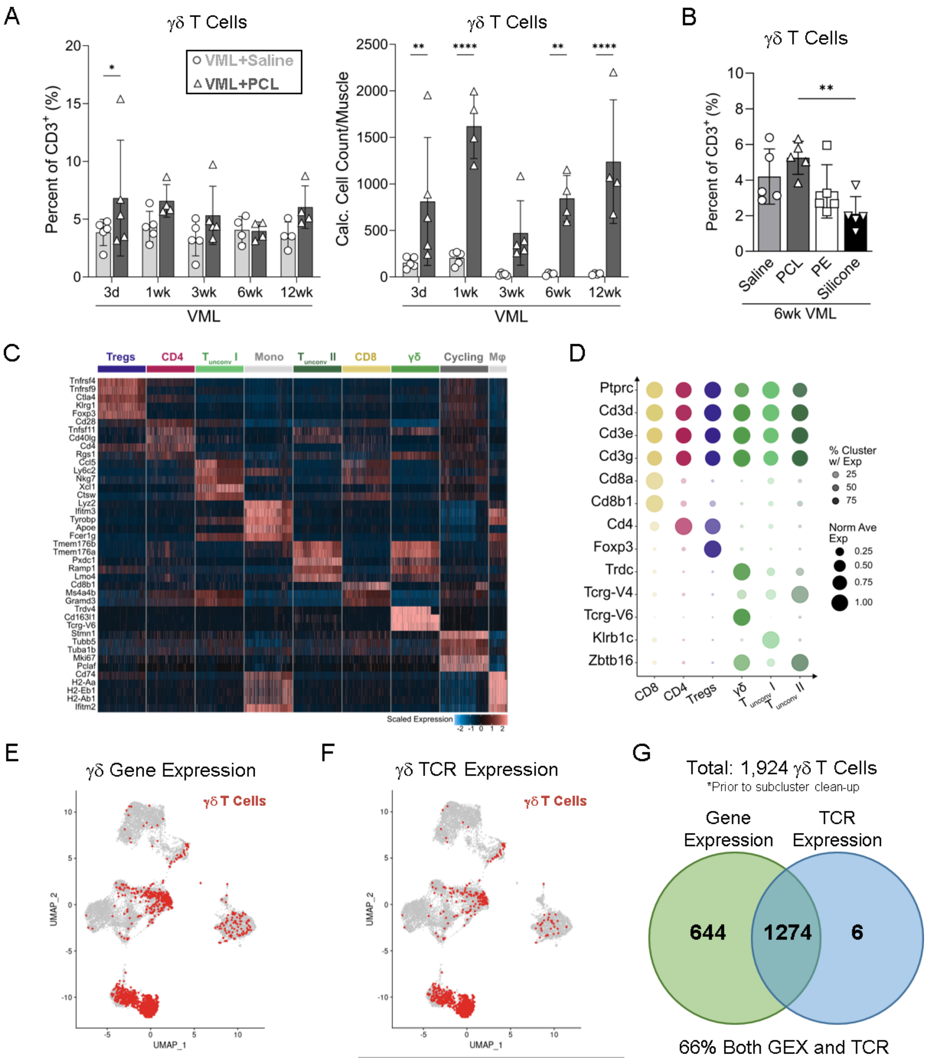

| α | β | γ | δ | Cell Type |
|---|---|---|---|---|
| Yes | ? | ? | No | **αβ T cell** |
| Yes | ? | ? | Yes | **Doublet** |
| ? | Yes | ? | No | **αβ T cell** |
| ? | Yes | ? | Yes | **Doublet** |
| **No** | **No** | **Yes** | **?** | **γδ T cell** |
| **No** | **No** | **?** | **Yes** | **γδ T cell** |

**Suppl Fig 1. Identification of γδ T cells in preclinical model of injury and biomaterial implant fibrosis.** **(A)** Flow cytometric quantification of γδ T cells in murine muscles at various time points (range: 3 days to 12 wks) after VML injury, either alone (VML+Saline) or with polycaprolactone implants (VML+PCL). **(B)** Flow cytometric quantification of γδ T cells 6 wks after VML injury with different synthetic biomaterial implants. PE: polyethylene. **(C)** Scaled expression heatmap of top differentially expressed genes by cluster in $CD3^+$ T cell scRNA/TCRseq dataset. **(D)** Dot plot of key marker genes for T cell clusters, where dot size corresponds to average expression (normalized to maximum expression) and transparency indicates percentage of cluster expressing the gene. **(E)** Overlay of γδ T cells (red) on $CD3^+$ T cell UMAP (top) identified by defined criteria for TCR variable region gene expression (bottom). **(F)** Overlay of γδ T cells (red) on $CD3^+$ T cell UMAP (top) identified by defined criteria for TCR sequences (bottom). **(G)** Quantification of and overlap between γδ T cells identified by gene and TCR expression criteria. **(Statistics)** Bar graph: mean±SD. Data analyzed using two-way ANOVA (A) or one-way ANOVA (B) with Tukey's multiple comparisons test. For A, only graphing statistics for "within time point" comparisons. NS: Not significant $p>0.05$, * $p<0.05$, ** $p<0.01$, *** $p<0.001$, **** $p<0.0001$.

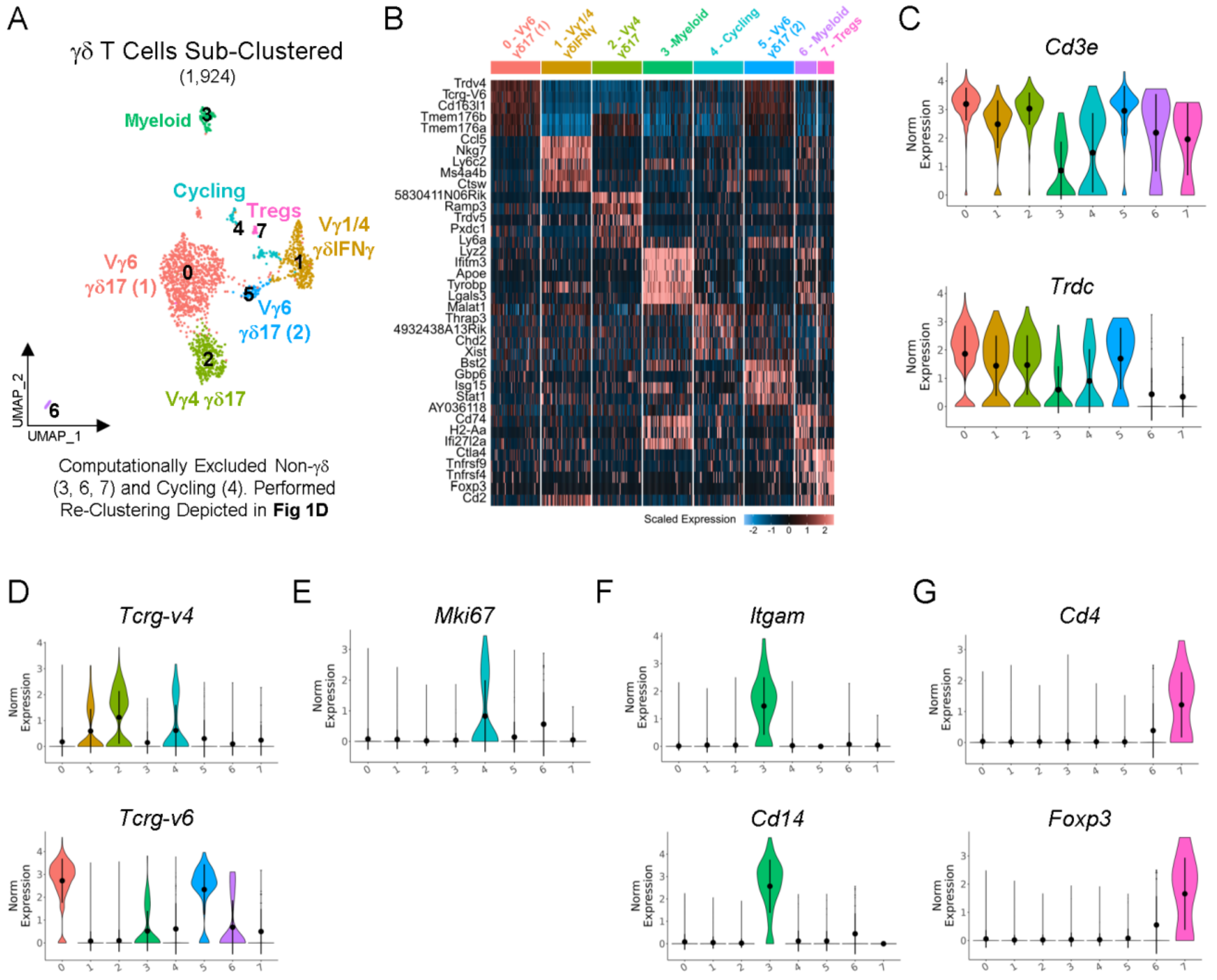

**<u>Suppl Fig 2</u>. Initial clustering of scRNAseq γδ T cells and removal of artifact clusters.**
**(A)** γδ T cells (1,924 cells) selected by gene and/or TCR criteria as outlined in **Suppl. Fig 1E-G** were processed and clustered using a standard pipeline (see Methods). UMAP visualization of the initial 8 γδ T cell clusters. **(B)** Scaled expression heatmap of top differentially expressed genes by initial γδ T cell clusters. **(C)** Violin plots of γδ T cell marker genes revealed low *Trdc* expression in clusters 3, 4, 6 and 7. **(D)** Violin plots of Vγ chains (*Tcrg-V4, Tcrg-V6*) across clusters. **(E)** Violin plot of cell proliferation marker gene (*Mki67*) identifies cluster 4 as proliferative. **(F)** Violin plots of myeloid marker genes (*Itgam, Cd14*) show high expression in cluster 3. **(G)** Violin plots of T regulatory (Treg) marker genes (*Cd4, Foxp3*) show high expression in cluster 7. Based on these results, clusters 3, 4, 6, and 7 were excluded from subsequent analyses and the remaining high-confidence γδ T cells (0, 1, 2, and 5) were re-clustered as depicted in **Fig 1D**.

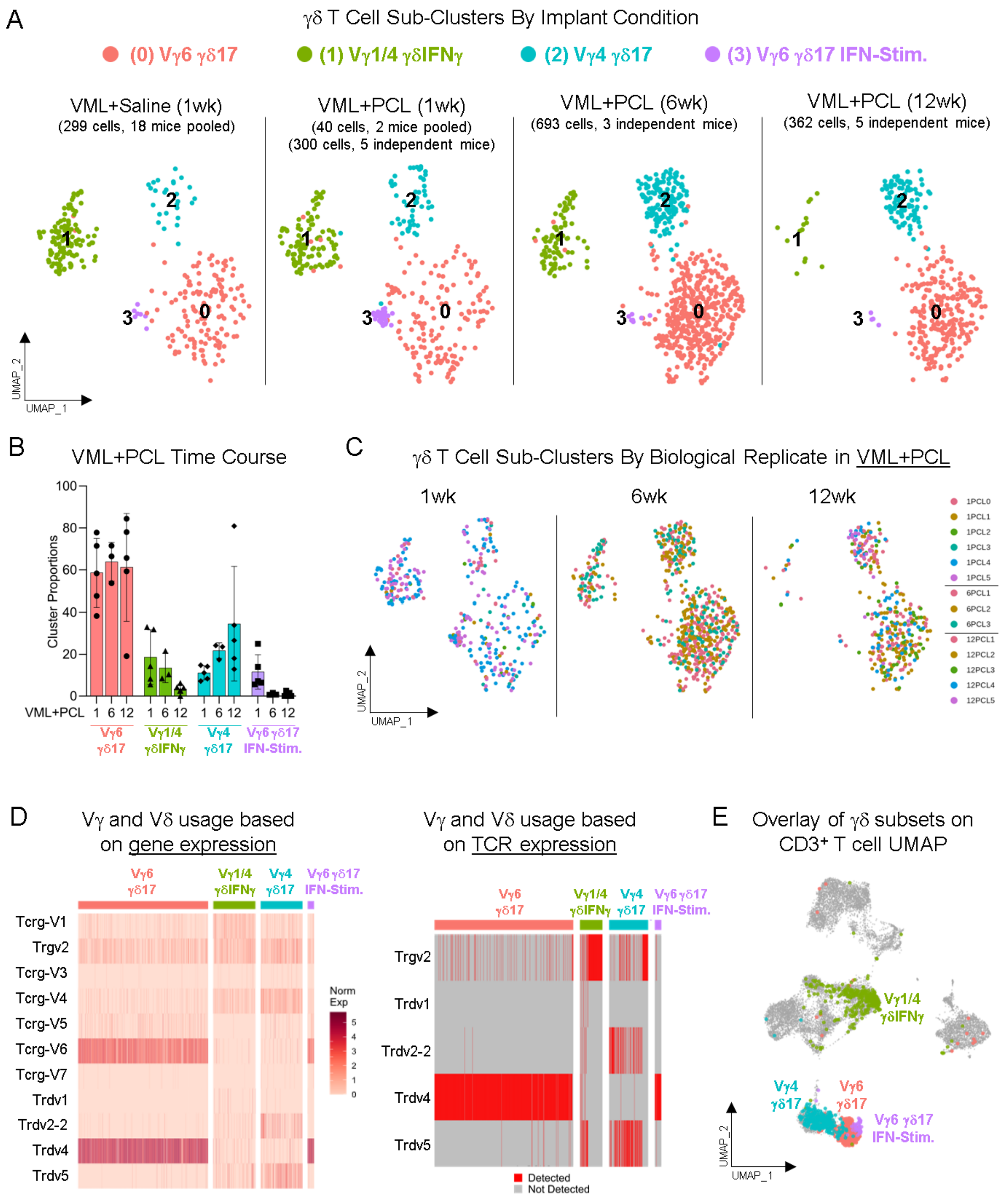

**Suppl Fig 3. Distribution of main γδ T cell clusters across experimental conditions.**
**(A)** UMAP of γδ T cell clusters separated by experimental conditions: VML+Saline (1 wk) and VML+PCL (1 wk, 6 wk, and 12 wk). **(B)** Cluster percentage of total γδ T cells per VML+PCL time point calculated by independent biological replicate; percentages across clusters sum to 100% per mouse. **(C)** γδ T cell UMAP depicting cells from independent biological replicates separated by VML+PCL time point. **(D)** Heatmap of γ and δ TCR variable region usage based on gene expression (left) and TCR expression (right) by γδ T cell clusters. **(E)** Overlay of γδ T cell clusters on CD3$^{+}$ T cell UMAP. **(Statistics)** Bar graph: mean±SD (B).

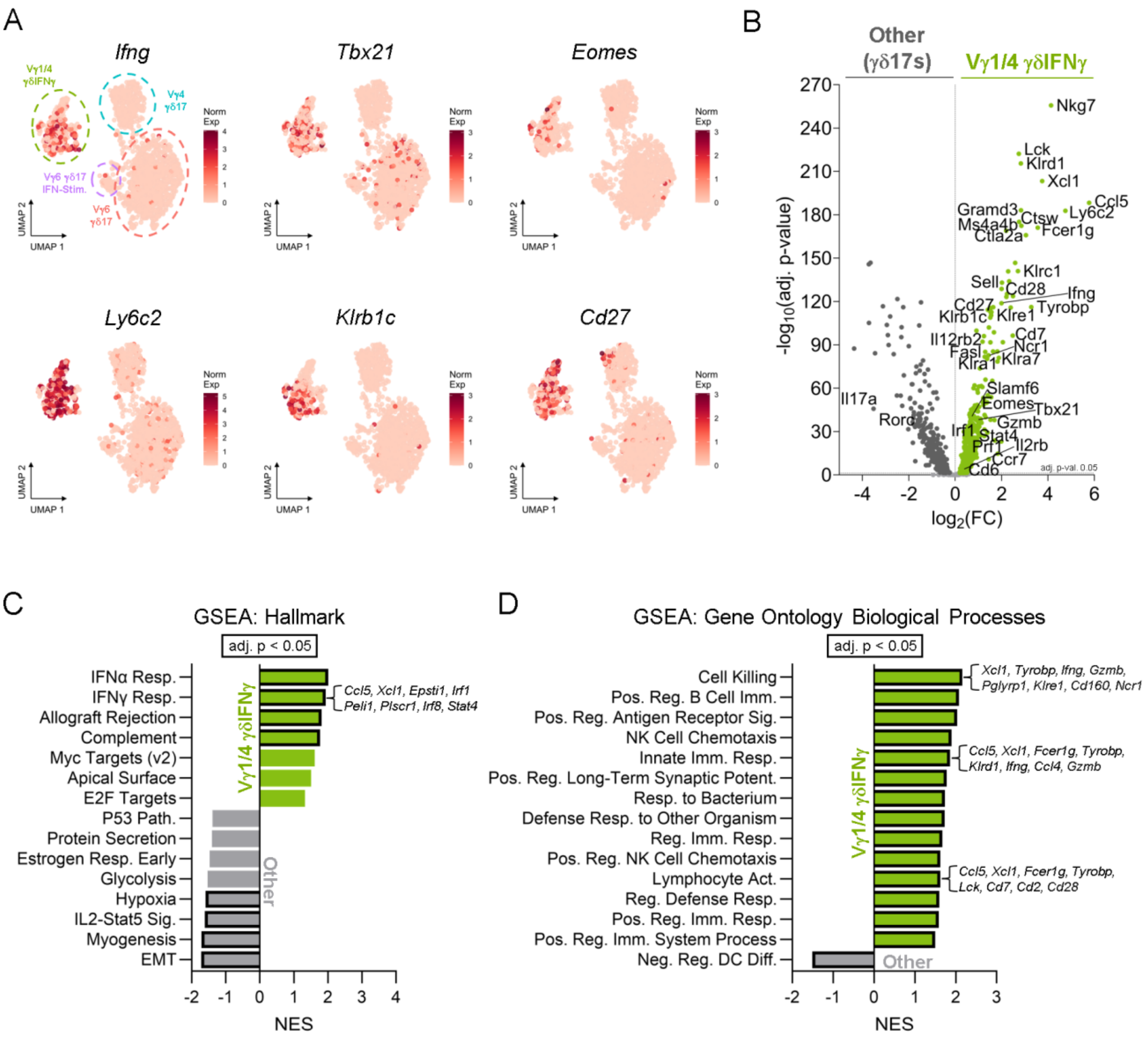

**<u>Suppl Fig 4</u>. Transcriptomic profiling of type-1 γδIFNγ scRNAseq cluster.**
**(A)** Feature plots displaying normalized expression of canonical type-1 (*Ifng, Tbx21*) and common γδIFNγ marker (*Cd27, Ly6c2, Klrb1c, Il2rb*) genes. **(B)** Volcano plot of differential gene expression (DE) between the "Vγ1/4 γδIFNγ" cluster and all other γδ T cells. **(C)** Gene set enrichment analysis (GSEA) using Hallmark gene sets based on DE results between the "Vγ1/4 γδIFNγ" cluster and all other γδ T cells with top 8 leading genes annotated for select pathways. NES: Normalized enrichment score. **(D)** GSEA using Gene Ontology Biological Processes gene sets based on DE results between the "Vγ1/4 γδIFNγ" cluster and all other γδ T cells with top 8 leading genes annotated for select pathways. **(Statistics)** For GSEA, the top 15 gene sets are selected based on lowest adj. p-value and plotted by NES with significant enrichment (adj. $p < 0.05$) indicated with a bold black outline.

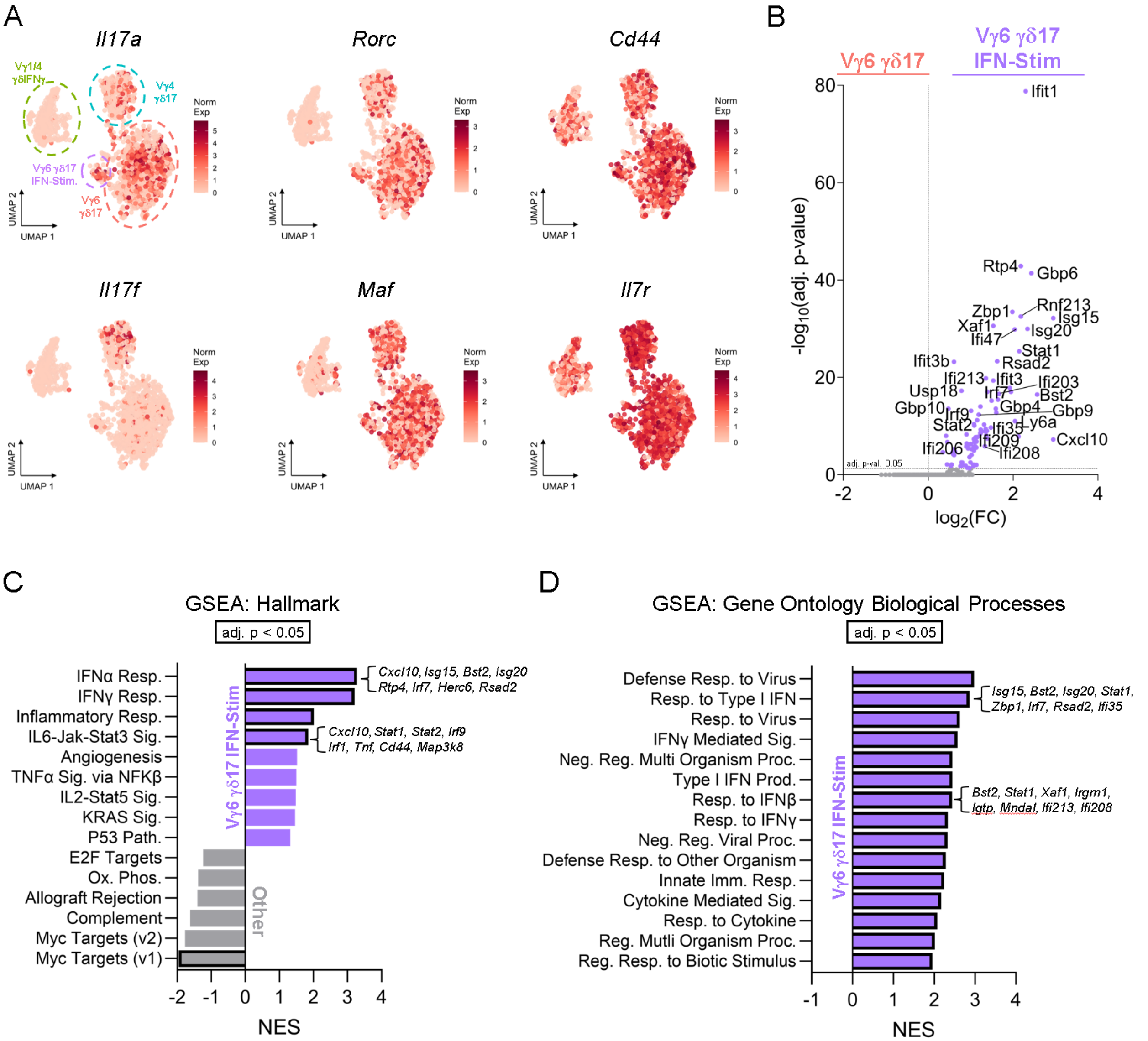

**<u>Suppl Fig 5</u>. Transcriptomic profiling of type-17 γδ17 scRNAseq clusters.**
**(A)** Feature plots displaying normalized expression of canonical type-17 (*Il17a, Il17f, Rorc, Maf*) and common γδ17 marker (*Cd44, Il7r*) genes. **(B)** Volcano plot of differential gene expression (DE) between the "Vγ6 γδ17" and the "Vγ6 γδ17 IFN-Stim." clusters. **(C)** Gene set enrichment analysis (GSEA) using Hallmark gene sets based on DE results between the ""Vγ6 γδ17" and the "Vγ6 γδ17 IFN-Stim." clusters with top 8 leading genes annotated for select pathways. NES: Normalized enrichment score. **(D)** GSEA using Gene Ontology Biological Processes gene sets based on DE results between the "Vγ6 γδ17" and the "Vγ6 γδ17 IFN-Stim." sub-clusters with top 8 leading genes annotated for select pathways. **(Statistics)** For GSEA, the top 15 gene sets are selected based on lowest adj. p-value and plotted by NES with significant enrichment (adj. p < 0.05) indicated with a bold black outline.

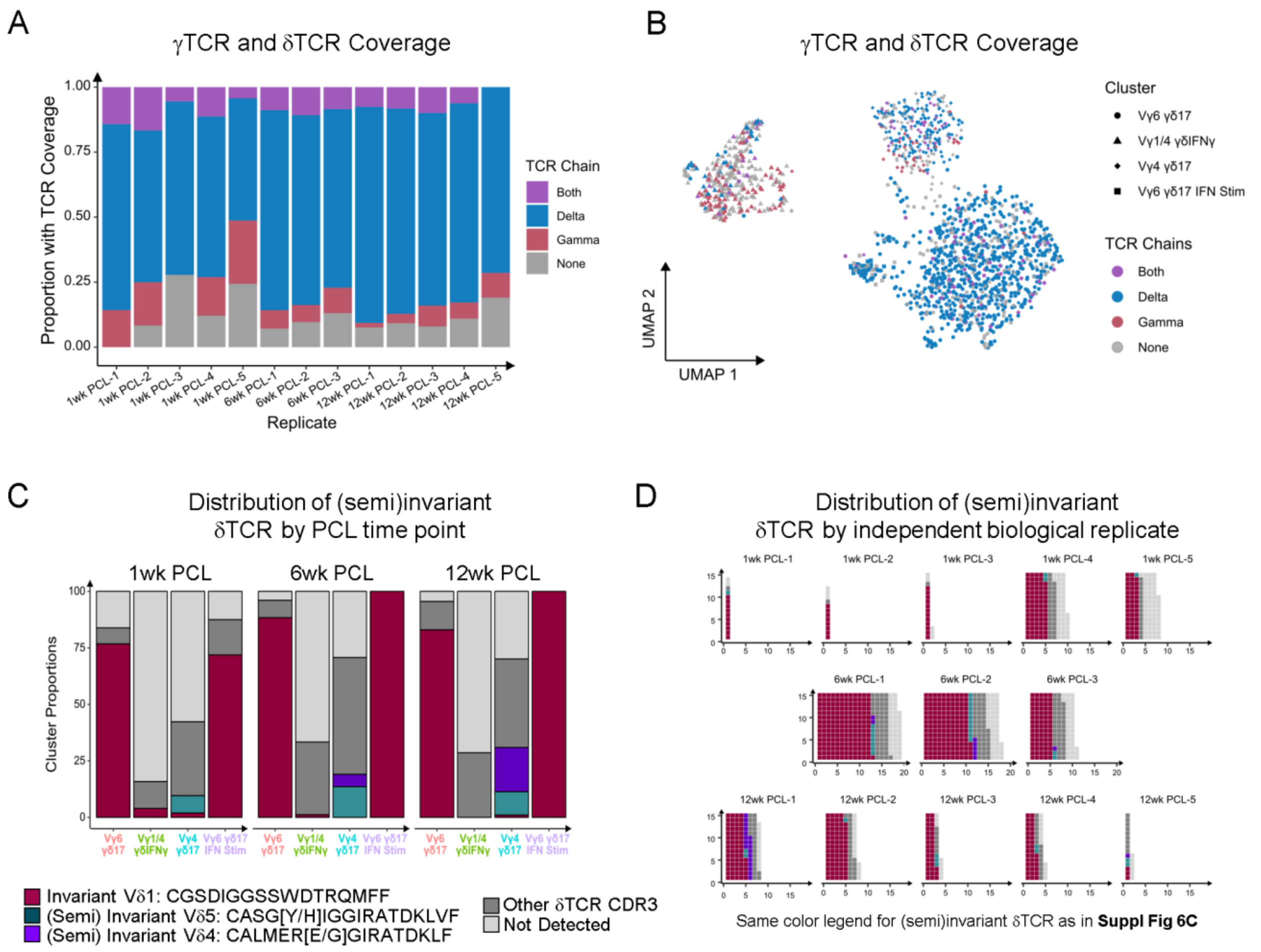

**<u>Suppl Fig 6</u>. scTCRseq analysis of γδ T cells.**

**(A)** Proportions of γδ T cell TCR sequencing coverage including detection of only γTCR (red), only δTCR (blue), both γTCR and δTCR (purple), or neither (gray) displayed by biological replicate. **(B)** Overlay of γTCR and δTCR sequencing coverage on γδ T cell cluster UMAP. **(C)** Proportions of γδ T cells with select (semi)invariant δTCR sequences displayed by PCL timepoint and individual biological replicate. **(D)** Waffle plot showing select (semi)invariant δTCR sequences across individual biological replicate; each square represents one cell.

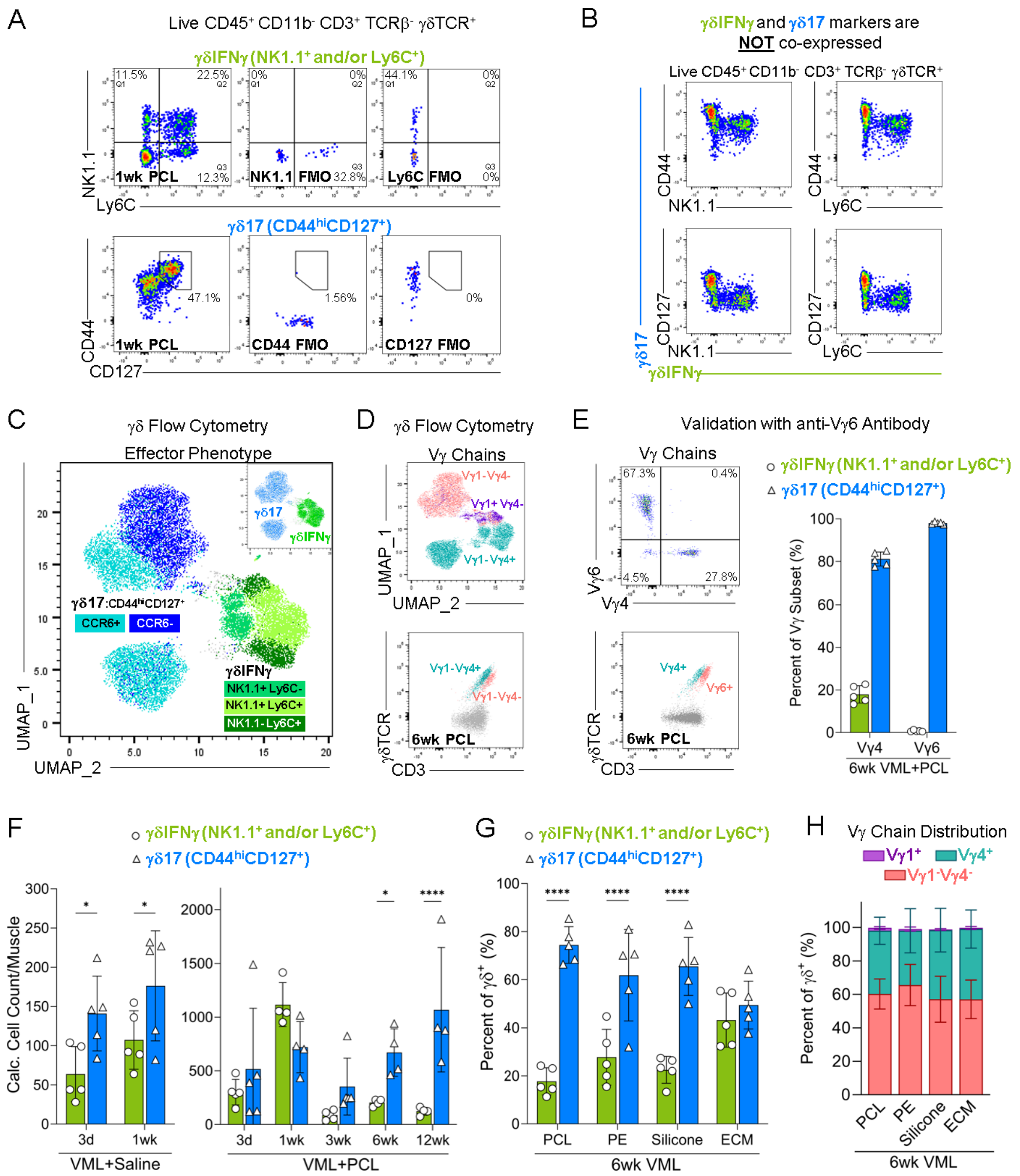

**Suppl Fig 7. Flow cytometric profiling of effector γδ T cells in tissue injury and implant fibrosis.** **(A)** Representative flow plots for γδIFNγ (NK1.1$^+$ and/or Ly6C$^+$) and γδ17 (CD44$^{hi}$CD127$^+$) with corresponding florescence minus one (FMO) controls. **(B)** Flow plots confirming lack of co-expression of γδIFNγ (x-axis) and γδ17 (y-axis) markers. **(C)** Overlay of gated γδIFNγ (green) and γδ17 (blue) subsets on flow UMAP of γδ T cells from VML+PCL time course (3 days, 1, 3, 6 and 12 wk). **(D)** Overlay of gated Vγ1+ (purple), Vγ4+ (teal), and Vγ1/4- (pink) subsets on flow UMAP of γδ T cells (top) and γδTCR versus CD3 plot (bottom). **(E)** Representative flow plot of Vγ6+ and Vγ4+ subsets (top left) and their relative CD3 expression (bottom left). Distribution of γδIFNγ and γδ17 based on Vγ expression (right). **(F)** Quantification of γδIFNγ and γδ17 counts/muscle in VML+Saline (left) and VML+PCL (right) at various time points. **(G)** Quantification of γδIFNγ and γδ17 in injured muscle with various implants at 6 wks. PE: polyethylene. ECM: extracellular matrix. **(H)** Distribution of Vγ chain expression by γδ T cells in injured muscle with various implants. **(Statistics)** Bar graphs: mean±SD. Data analyzed using two-way ANOVA with Tukey's multiple comparisons test (F, G). Only displaying statistics within time point (F) or material (G). NS: Not significant $p>0.05$, * $p<0.05$, ** $p<0.01$, *** $p<0.001$, **** $p<0.0001$.

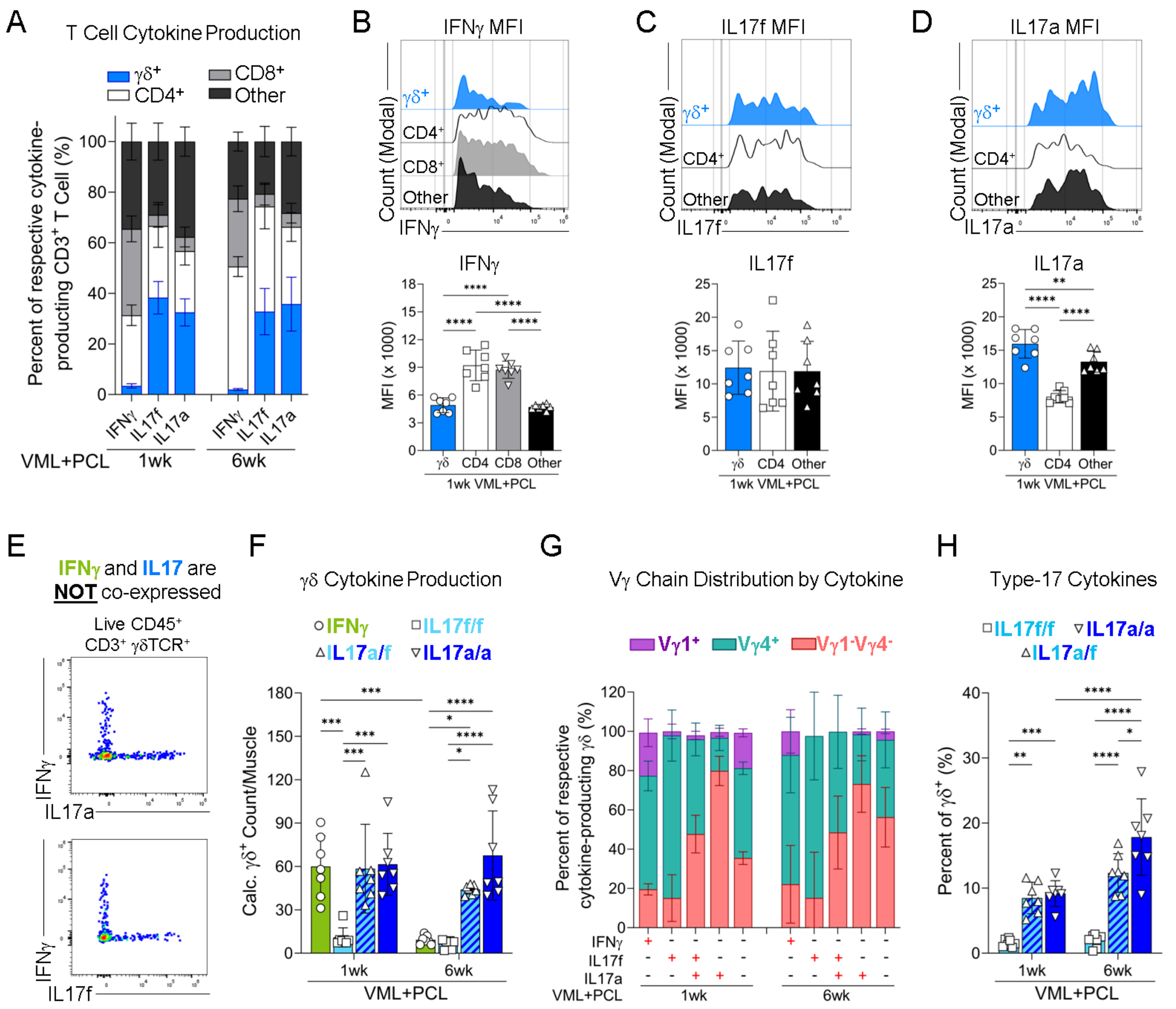

**Suppl Fig 8. Flow cytometric profiling of γδ T cell cytokine production in implant fibrosis.**
**(A)** Distribution of $CD3^+$ T cell subsets (γδ, $CD4^+$, $CD8^+$, other: $CD3^+γδ^-CD4^-CD8^-$) that produce IFNγ, IL17f, or IL17a in the VML+PCL environment at 1 and 6 wks. **(B)** Representative histogram (top) and median fluorescence intensity (MFI) quantification (bottom) for IFNγ by various $IFNγ^+$ T cells. Only displaying 1 wk due to low IFNγ production at 6 wk. **(C)** Representative histogram (top) and MFI quantification (bottom) for IL17f by various $IL17f^+$ T cells. **(D)** Representative histogram (top) and MFI quantification (bottom) for IL17a by various $IL17a^+$ T cells. **(E)** Flow plots depicting lack of co-expression of IFNγ and IL17 cytokines by γδ T cells. **(F)** Quantification of γδ T cell counts/muscle that produce cytokines (IFNγ, IL17a/a, IL17a/f, IL17f/f) following *ex vivo* stimulation in VML+PCL at 1 and 6 wks. **(G)** Distribution of Vγ chain expression by cytokine-producing γδ T cell sub-population in VML+PCL. **(H)** Distribution of $IL17^+$ γδ T cells based on their IL17 production profile (IL17a/a, IL17f/f, or IL17a/f) in VML+PCL at 1 and 6 wks. **(Statistics)** Bar graphs: mean±SD. Data analyzed using repeated measures one-way ANOVA (B-D) or two-way ANOVA (F, H) with Tukey's multiple comparisons test. NS: Not significant $p>0.05$, * $p<0.05$, ** $p<0.01$, *** $p<0.001$, **** $p<0.0001$.

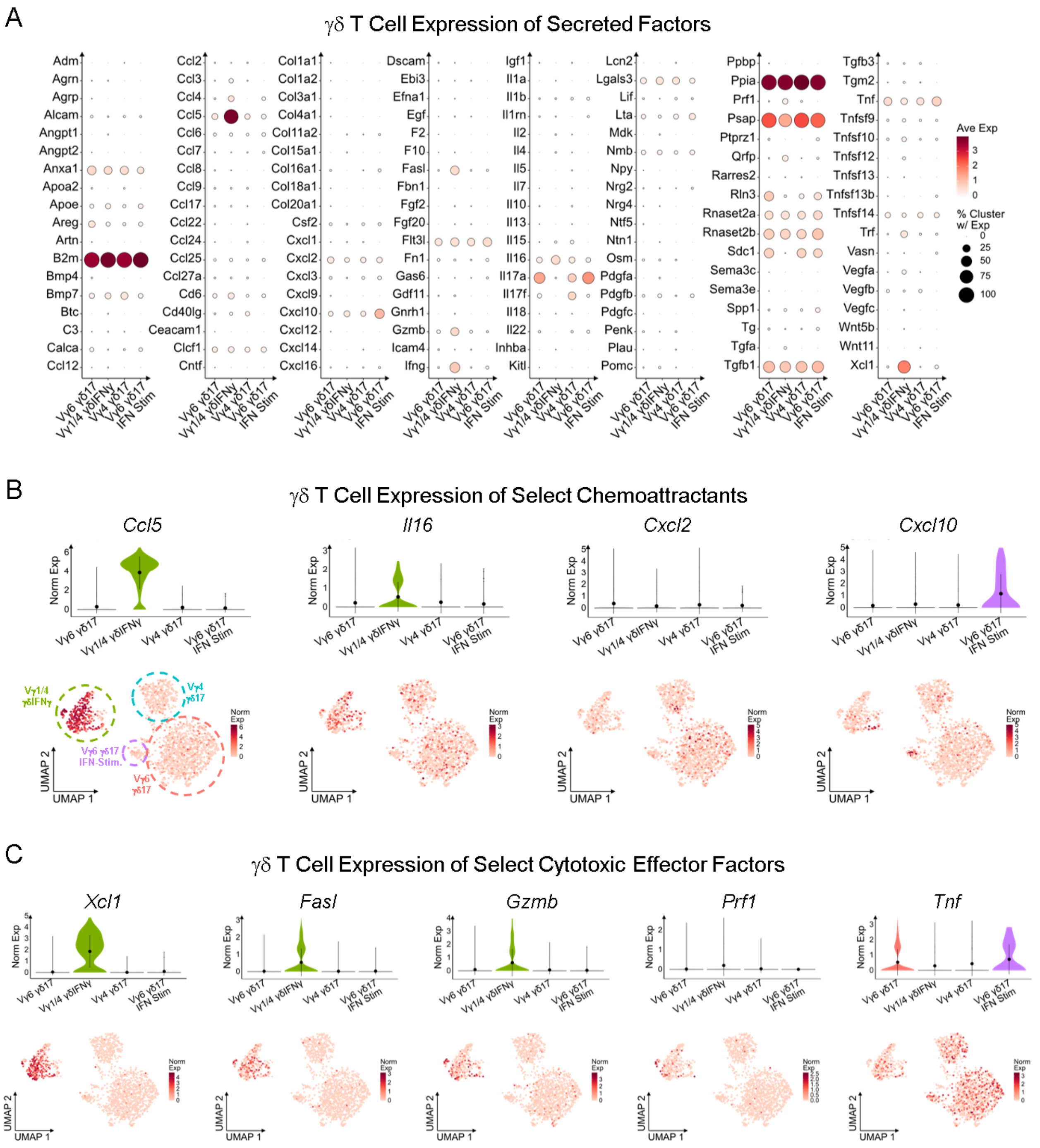

**<u>Suppl Fig 9</u>. γδ T cell subsets express a myriad of secreted immunomodulatory factors.**
**(A)** Average expression (avg. exp.) of all detected secreted factors by scRNAseq γδ T cell clusters (no thresholds for percentage of cells with detected expression or for avg. exp.). **(B)** Violin (top) and feature (bottom) plots displaying normalized expression (norm. exp.) of select cytokines (*Il16*) and chemokines (*Ccl5, Cxcl2, Cxcl10*) by scRNAseq γδ T cell clusters. **(C)** Violin (top) and feature (bottom) plots displaying norm. exp. of select cytotoxic effector factors (*Xcl1, Gzmb, Fasl, Prf1, Tnf*) by scRNAseq γδ T cell clusters. Violin plots depict data distribution; overlaid point and line indicate mean±SD.

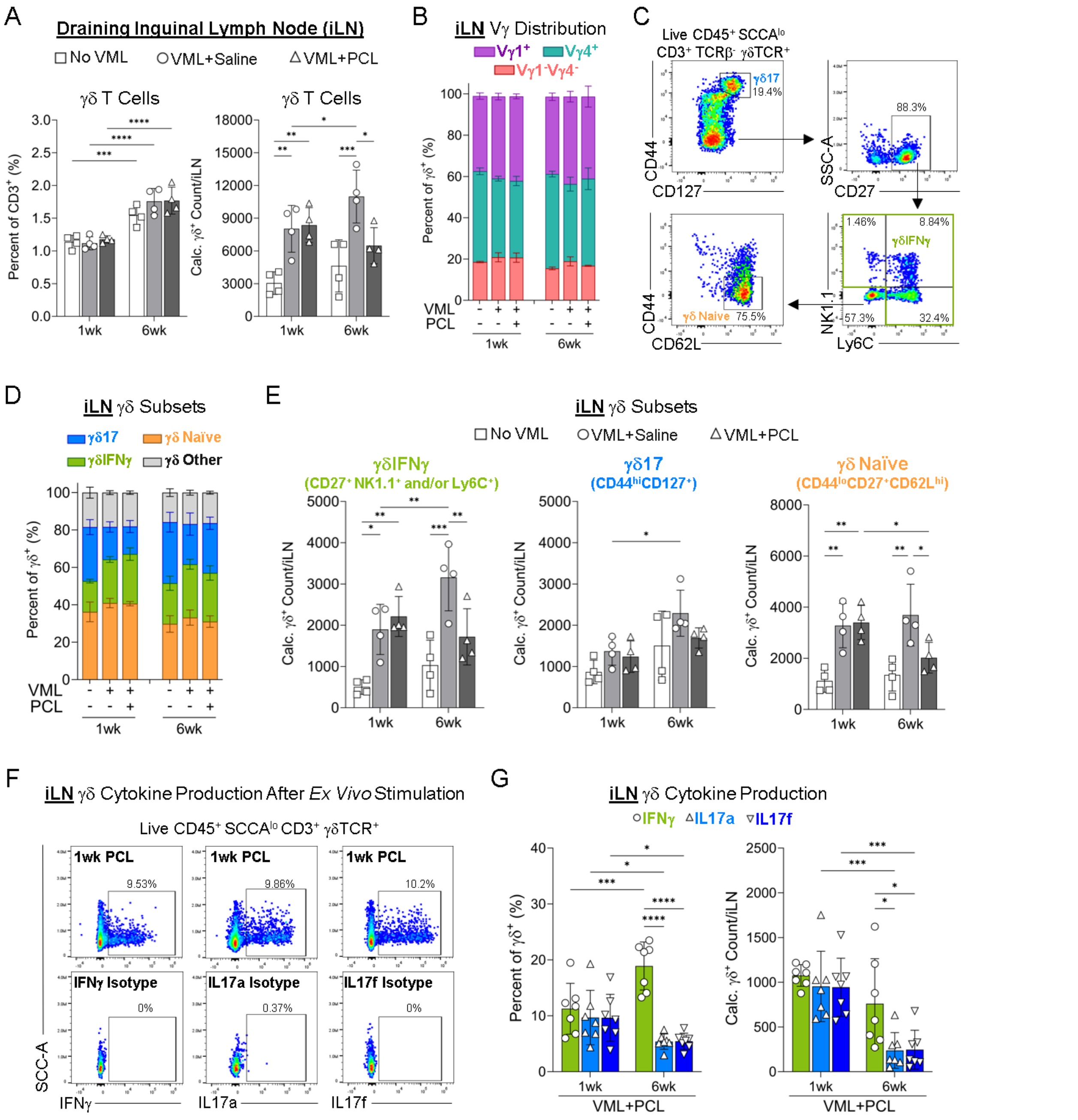

**Suppl Fig 10. Flow cytometric profiling of γδ T cell effector subsets in draining lymph nodes.** **(A)** Quantification of γδ T cell percentage (left) and counts (right) in draining inguinal LNs (iLNs) of uninjured (No VML), injured (VML+Saline), and injured with PCL implants (VML+PCL) mice at 1 and 6 wks. **(B)** Distribution of Vγ chain expression by γδ T cells in iLNs. **(C)** Representative flow gating of γδ17 (CD44$^{hi}$CD127$^{+}$), γδIFNγ (CD27$^{+}$, NK1.1$^{+}$ and/or Ly6C$^{+}$), and γδ naïve (CD44$^{-}$CD27$^{+}$CD62L$^{hi}$) subsets in iLNs. **(D)** Distribution of γδ T cell subsets in iLNs. **(E)** Quantification of γδIFNγ (left), γδ17 (middle), and γδ naïve (right) cell counts/iLN. **(F)** Representative flow plots of cytokine production by γδ T cells in iLNs following *ex vivo* stimulation with corresponding isotype controls. **(G)** Quantification of percentage (left) and cell counts (right) of cytokine production (IFNγ, IL17a, IL17f) by γδ T cells in iLN of mice with VML+PCL (1 and 6 wks). **(Statistics)** Bar graphs: mean±SD. Data analyzed using two-way ANOVA with Tukey's multiple comparisons test (A, E, G). NS: Not significant $p>0.05$, * $p<0.05$, ** $p<0.01$, *** $p<0.001$, **** $p<0.0001$.

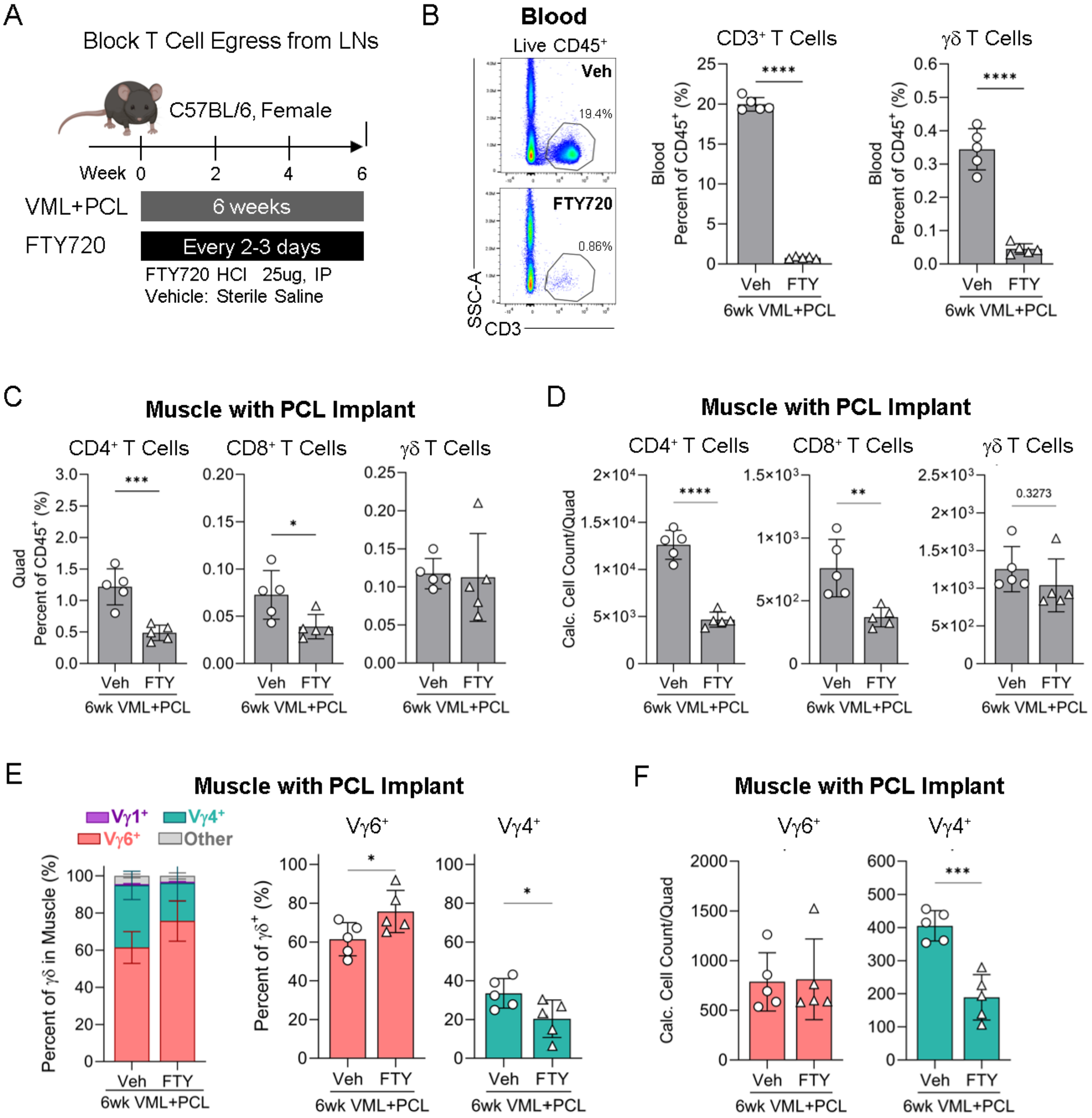

**Suppl Fig 11. Inhibition of T cell egress from lymphoid tissue using Fingolimod HCl (FTY720).**
**(A)** Experiment scheme. FTY720 HCl (25 μg/dose) or sterile saline (Veh: vehicle control) was administered via intraperitoneal injection at the time of VML injury with PCL implantation (day 0), again on day 1, and then every 2-3 days for the remainder of the 6 wk experiment. **(B)** Representative flow plots (left) and quantification of CD3$^+$ T cells (middle) and γδ T cells (right) in peripheral blood with FTY720 treatment. **(C)** Percentage of T cell subsets (CD4$^+$, CD8$^+$, γδ) of total CD45$^+$ immune cells in VML-injured muscles with PCL (VML+PCL) following FTY720 treatment. **(D)** Quantification of T cell subset (CD4$^+$, CD8$^+$, γδ) cell counts in VML+PCL following FTY720 treatment. **(E)** Distribution of Vγ chain expression (Vγ1, Vγ4, Vγ6, other) by γδ T cells in VML+PCL following FTY720 treatment. **(F)** Quantification of Vγ6$^+$ and Vγ4$^+$ γδ T cell counts in VML+PCL following FTY720 treatment. **(Statistics)** Bar graphs: mean±SD. Data analyzed using unpaired two-tailed student t-test (B-F). NS: Not significant p>0.05, * p<0.05, ** p<0.01, *** p<0.001, **** p<0.0001.

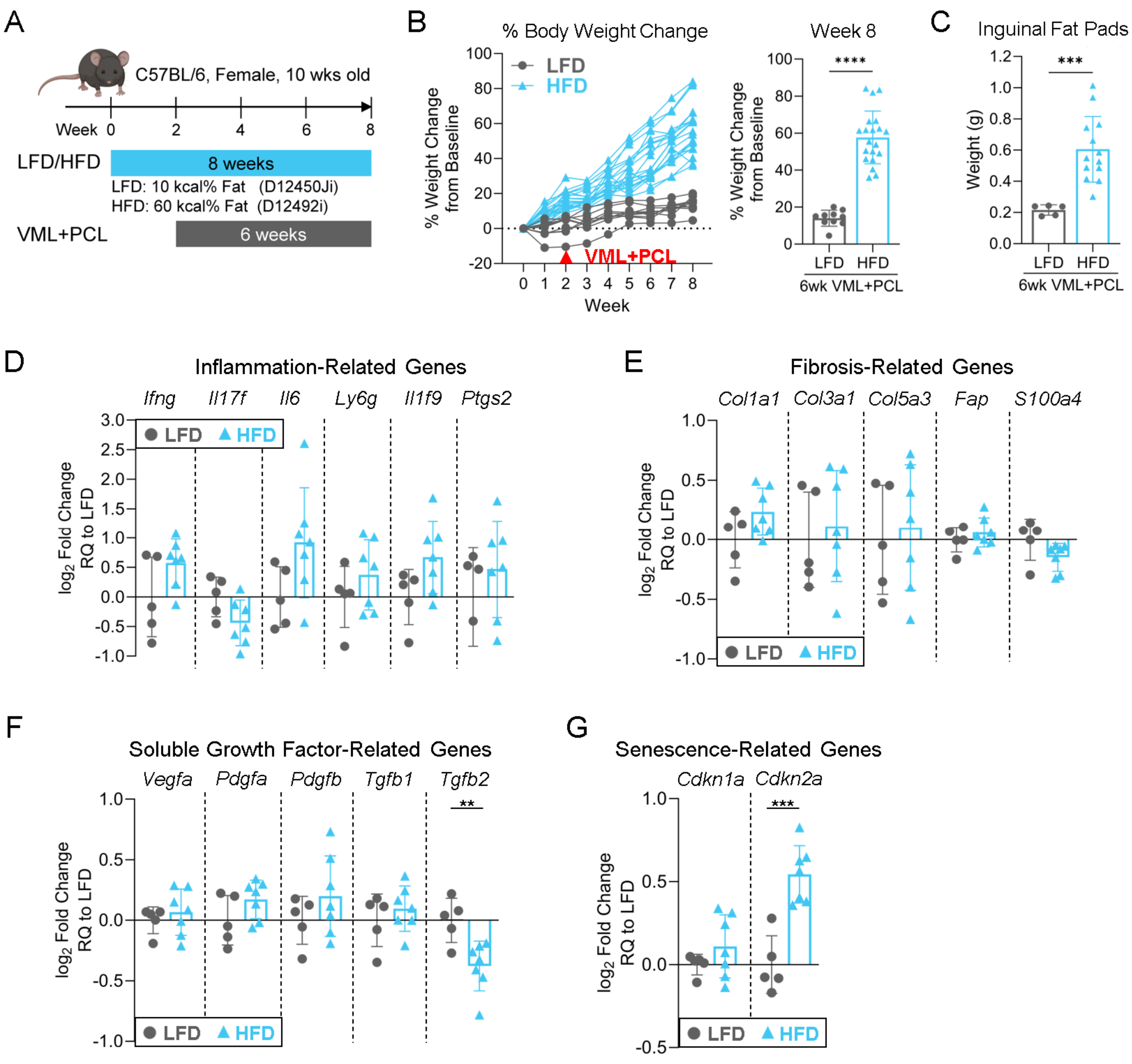

**Suppl Fig 12. Impact of high fat diet (HFD) on mouse weight and gene expression in implant fibrosis. (A)** Experiment scheme. Mice were randomized between low fat diet (LFD, control) and high fat diet (HFD) groups, with new *ad libitum* diet beginning 2 wks prior to VML injury with PCL implantation (VML+PCL) and continuing for the remaining 6 wks of the experiment. **(B)** Percent change in mouse body weight relative to baseline measured at 0 wk (before start of new diets) represented as longitudinal spider plot (left) and at final experiment time point (8 wk) (right). **(C)** Weight of inguinal fat pads from LFD- and HFD-fed mice at 8 wk time point from separate independent experiment. **(D)** Expression of inflammation-related genes (*Ifng, Il17f, Il6, Ly6g, Il1f9, Ptgs2*) in bulk VML+PCL tissue of HFD-fed mice relative to LFD-fed controls using RT-qPCR. **(E)** Expression of fibrosis-related genes (*Col1a1, Col31, Col5a3, Fap, S100a4*) in bulk VML+PCL tissue of HFD-fed mice relative to LFD-fed controls using RT-qPCR. **(F)** Expression of growth factor-related genes (*Vegfa, Pdgfa, Pdgfb, Tgfb1, Tgfb2*) in bulk VML+PCL tissue of HFD-fed mice relative to LFD-fed controls using RT-qPCR. **(G)** Expression of senescence-related genes (*Cdkn1a, Cdkn2a*) in bulk VML+PCL tissue of HFD-fed mice relative to LFD-fed controls using RT-qPCR. **(Statistics)** Bar graphs: mean±SD. Data analyzed using unpaired two-tailed student t-test (B-G). NS: Not significant $p>0.05$, * $p<0.05$, ** $p<0.01$, *** $p<0.001$, **** $p<0.0001$.

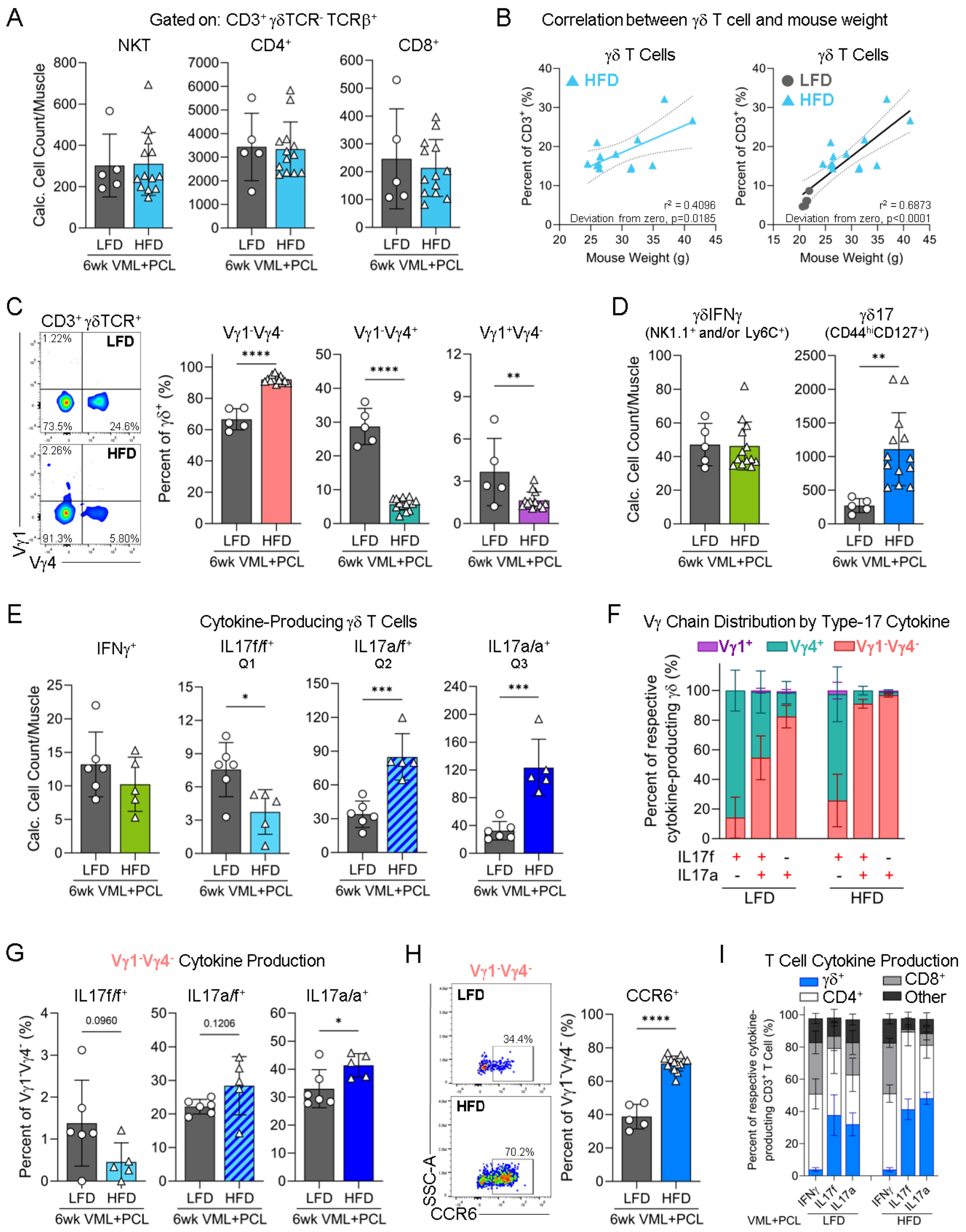

**Suppl Fig 13. Flow cytometric profiling of high fat diet (HFD) impact on γδ T cells in implant fibrosis. (A)** Quantification of non-γδ T cell subsets in VML injured muscle with PCL implants (VML+PCL) with HFD. **(B)** Linear regression to correlate percentage of γδ T cells and mouse body weight in only HFD-fed mice (left) and both LFD- and HFD-fed mice (right). **(C)** Representative flow plots (left) and quantification (right) of Vγ chain expression by γδ T cells in VML+PCL with HFD. **(D)** Quantification of γδIFNγ (NK1.1⁺ and/or Ly6C⁺) and γδ17 (CD44hiCD127⁺) in VML+PCL with HFD. **(E)** Quantification of γδ T cell cytokine production (IFNγ, IL17f/f, IL17a/f, IL17a/a) in VML+PCL with HFD. **(F)** Distribution of Vγ chain expression by IL17-producing γδ T cells in VML+PCL with HFD. **(G)** Quantification of Vγ1⁻Vγ4⁻ IL17 production in VML+PCL with HFD. **(H)** Representative flow plots (left) and quantification of CCR6 expression by Vγ1⁻Vγ4⁻. **(I)** Distribution of CD3⁺ T cell subsets that produce IFNγ, IL17f, or IL17a in VML+PCL with HFD. **(Statistics)** Bar graphs: mean±SD. Data analyzed using unpaired two-tailed student t-test (A, C-E, G, H) and simple linear regression (B). NS: Not significant $p>0.05$, * $p<0.05$, ** $p<0.01$, *** $p<0.001$, **** $p<0.0001$.

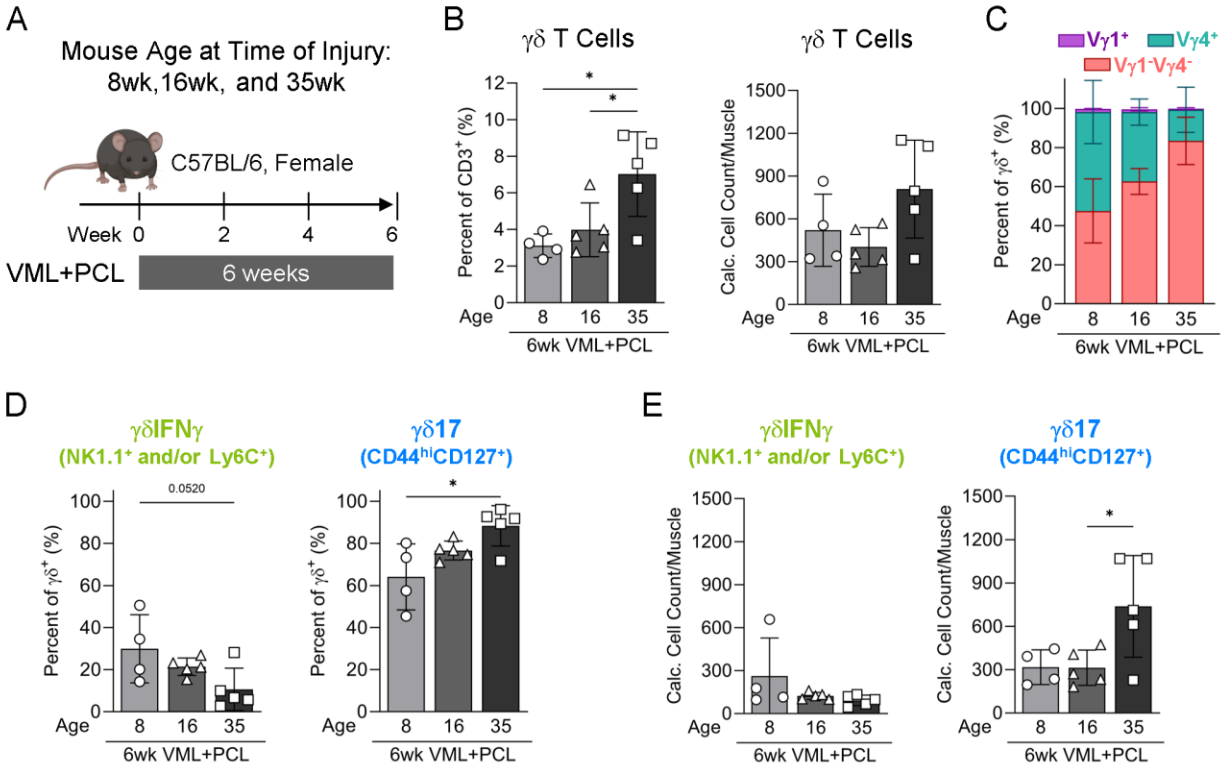

**Suppl Fig 14. Flow cytometric profiling of aging impact on γδ T cells in implant fibrosis.**
**(A)** Experiment scheme. VML injury with PCL implantation (VML+PCL) was performed on mice of varying age (8, 16, and 35 wk). **(B)** Quantification of γδ T cell percentage (left) and counts (right) in VML+PCL at 6 wks with varying mouse age. **(C)** Distribution of Vγ chain expression by γδ T cells in VML+PCL with varying mice age. **(D)** Quantification of γδIFNγ (NK1.1$^+$ and/or Ly6C$^+$) and γδ17 (CD44$^{hi}$CD127$^+$) in VML+PCL with varying mouse age. **(E)** Calculated cell counts of γδIFNγ and γδ17 in VML+PCL with varying mouse age. **(Statistics)** Bar graphs: mean±SD. Data analyzed using one-way ANOVA with Tukey's multiple comparisons test (B, D, E). NS: Not significant p>0.05, * p<0.05, ** p<0.01, *** p<0.001, **** p<0.0001.

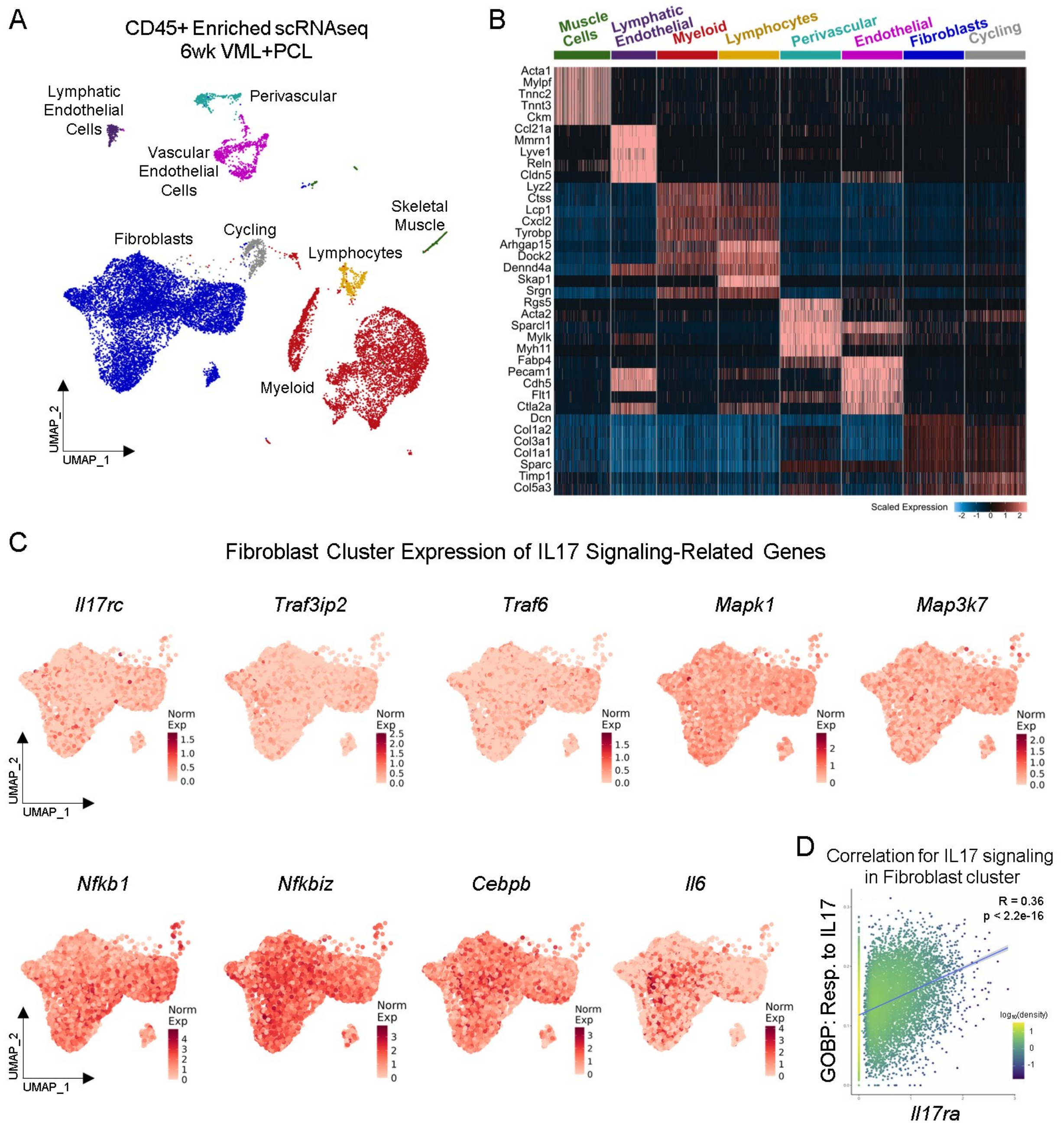

**<u>Suppl Fig 15</u>. Fibroblast scRNAseq cluster in implant fibrosis expresses IL17 signaling genes.** **(A)** Both CD45$^{+}$ immune and CD45$^{-}$ stromal cell fractions were collected from VML injured muscles with PCL implants (VML+PCL) at 6 wks for scRNAseq analysis (n = 3 independent replicates). UMAP of broad scRNAseq clusters including immune (myeloid, lymphocytes) and stromal (fibroblasts, vascular endothelial cells, perivascular cells, lymphatic endothelial cells). **(B)** Scaled expression heatmap of top differentially expressed genes by immune and stromal cell clusters present at 6 wks in VML+PCL. **(C)** Cropped feature plots of normalized expression of IL17 signaling related genes (*Il17rc, Traf3ip2, Traf6, Mapk1, Map3k7, Nfbk1, Nfkbiz, Cebpb, Il6*) by the fibroblast cluster. **(D)** Correlation of *Il17ra* gene expression and "Response to Interleukin-17" (GO:0097396) score by the fibroblast scRNAseq cluster. **(Statistics)** R and p-value from Pearson correlation test (D).

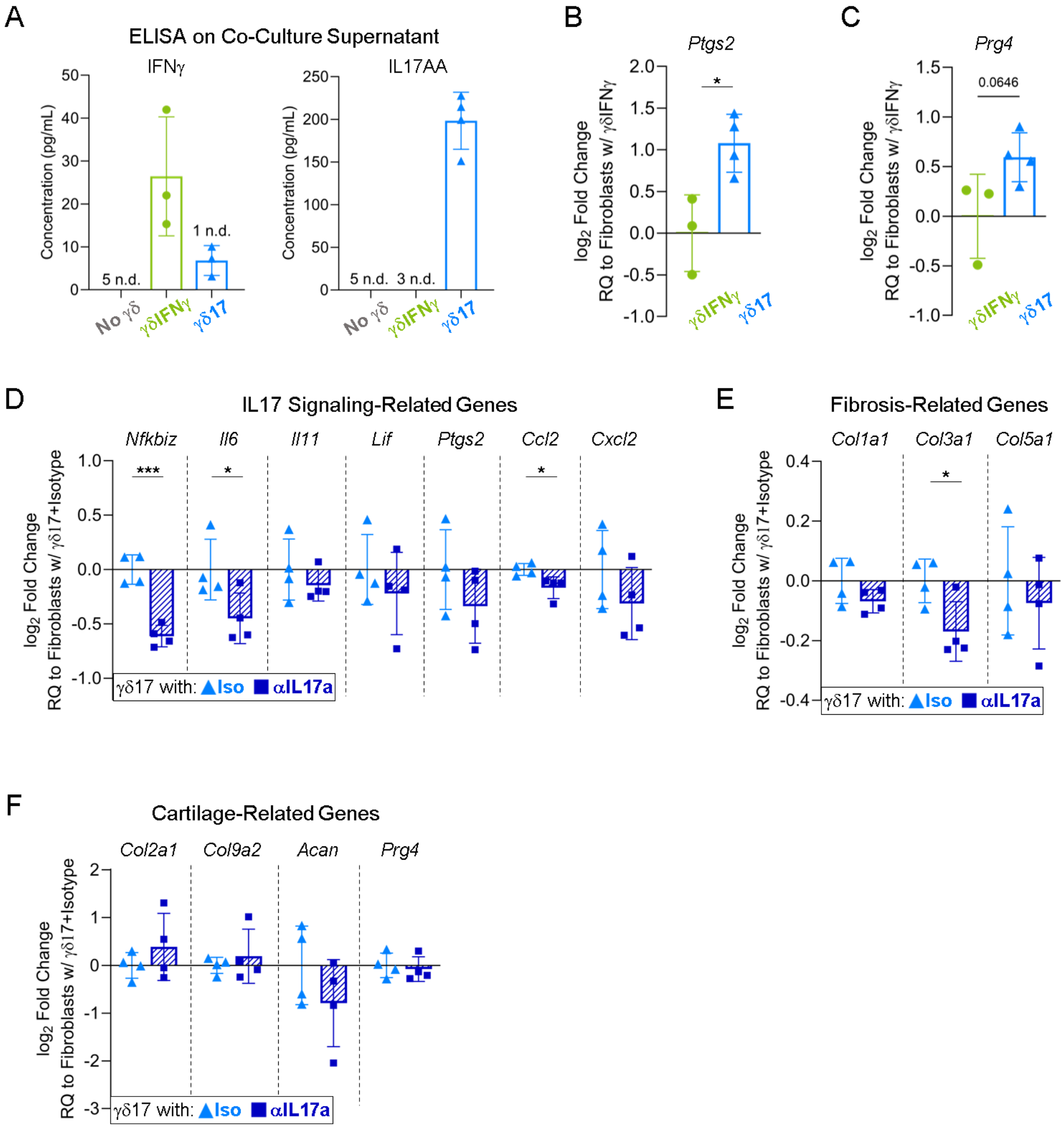

**Suppl Fig 16.** **γδ T cell secretome affects fibroblast gene expression in Transwell co-culture system.** **(A)** Murine dermal fibroblasts were co-cultured for 48 hours with murine splenic γδ T cells skewed towards either γδIFNγ or γδ17 effector phenotypes. γδ T cell production for IFNγ or IL17A homodimer (IL17AA) by respective effector subtype was confirmed via enzyme linked immunosorbent assay (ELISA) on co-culture supernatant. N.D.: not detected. **(B)** Fibroblast expression of *Ptgs2* after co-culture with γδ17 relative to γδIFNγ control using RT-qPCR. **(C)** Fibroblast expression of *Prg4* after co-culture with γδ17 relative to γδIFNγ control using RT-qPCR. **(D)** To block IL17a-mediated signaling, anti-IL17a antibody (or isotype control) was introduced into media for entire co-culture duration. Fibroblast expression of IL17 signaling-related genes (*Nfkbiz, Il6, Il11, Lif, Ptgs2, Ccl2, Cxcl2)* after γδ17 co-culture with αIL17a relative to isotype control using RT-qPCR. **(E)** Fibroblast expression of fibrosis-related genes (*Col1a1, Col3a1, Col5a1)* after γδ17 co-culture with αIL17a relative to isotype control using RT-qPCR. **(F)** Fibroblast expression of cartilage-related genes (*Col2a1, Col9a2, Acan, Prg4)* after γδ17 co-culture with αIL17a relative to isotype control using RT-qPCR. **(Statistics)** Bar graphs: mean±SD. Data analyzed using unpaired two-tailed student t-test (B-F). NS: Not significant $p>0.05$, * $p<0.05$, ** $p<0.01$, *** $p<0.001$, **** $p<0.0001$.

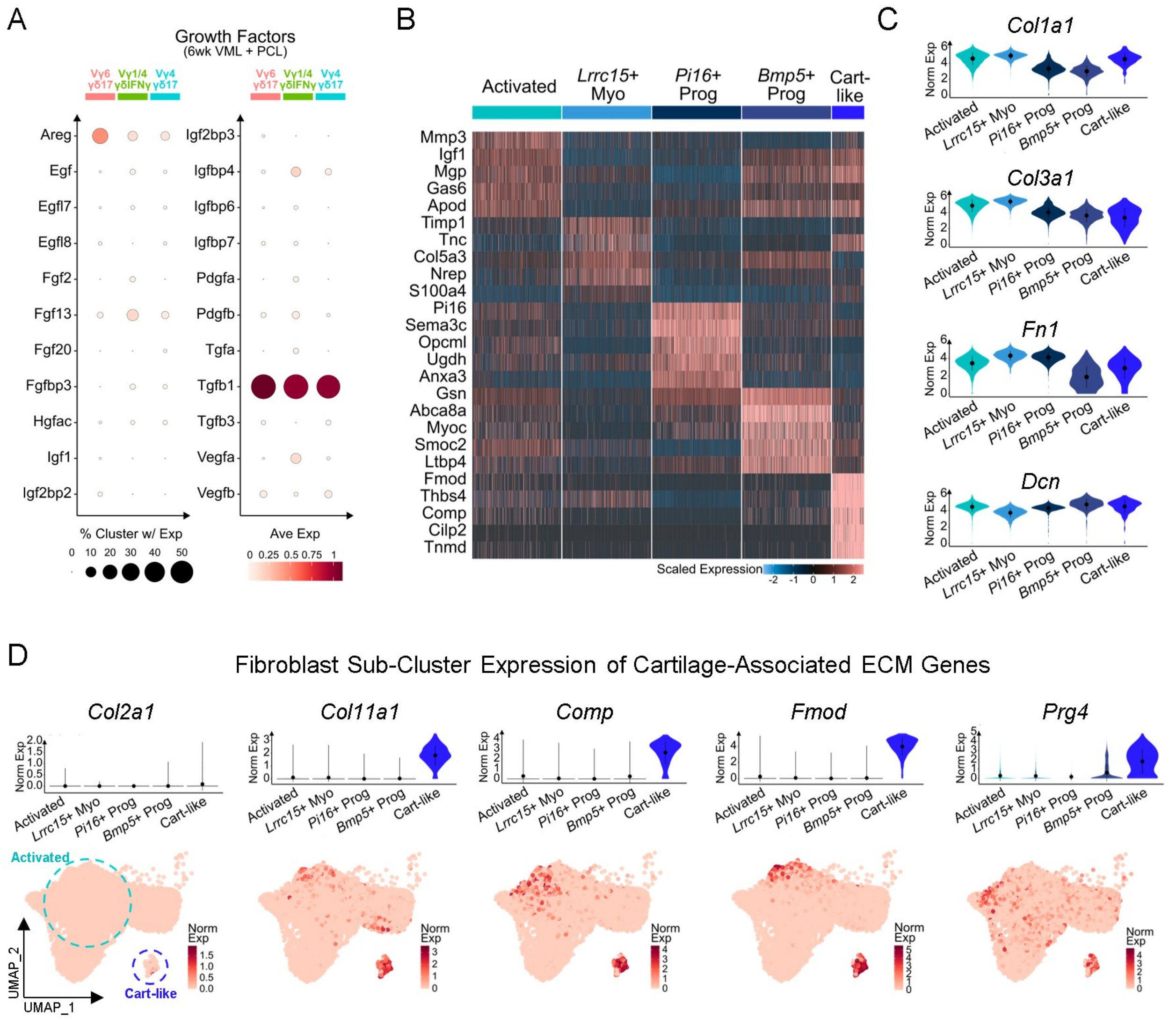

**<u>Suppl Fig 17</u>. Fibroblast scRNAseq sub-clusters in implant fibrosis express ECM components.**

**(A)** Average expression normalized to maximum expression (avg. exp.) of secreted growth factors by scRNAseq $\gamma\delta$ T cell clusters in VML+PCL at 6 wks (no thresholds for percentage of cells with detected expression or for avg. exp.). **(B)** Scaled expression heatmap of top differentially expressed genes by fibroblast sub-clusters present in VML+PCL at 6 wks. **(C)** Violin plots of normalized expression (norm. exp.) of common ECM components (*Col1a1, Col3a1, Fn1, Dcn*) by fibroblast sub-clusters in VML+PCL at 6 wks. **(D)** Violin (top) and cropped feature (bottom) plots of norm. exp. of canonical cartilage-associated ECM components (*Col2a1, Col11a1, Comp, Fmod, Prg4*) by fibroblast sub-clusters in VML+PCL at 6 wks. Violin plots depict data distribution; overlaid point and line indicate mean±SD.

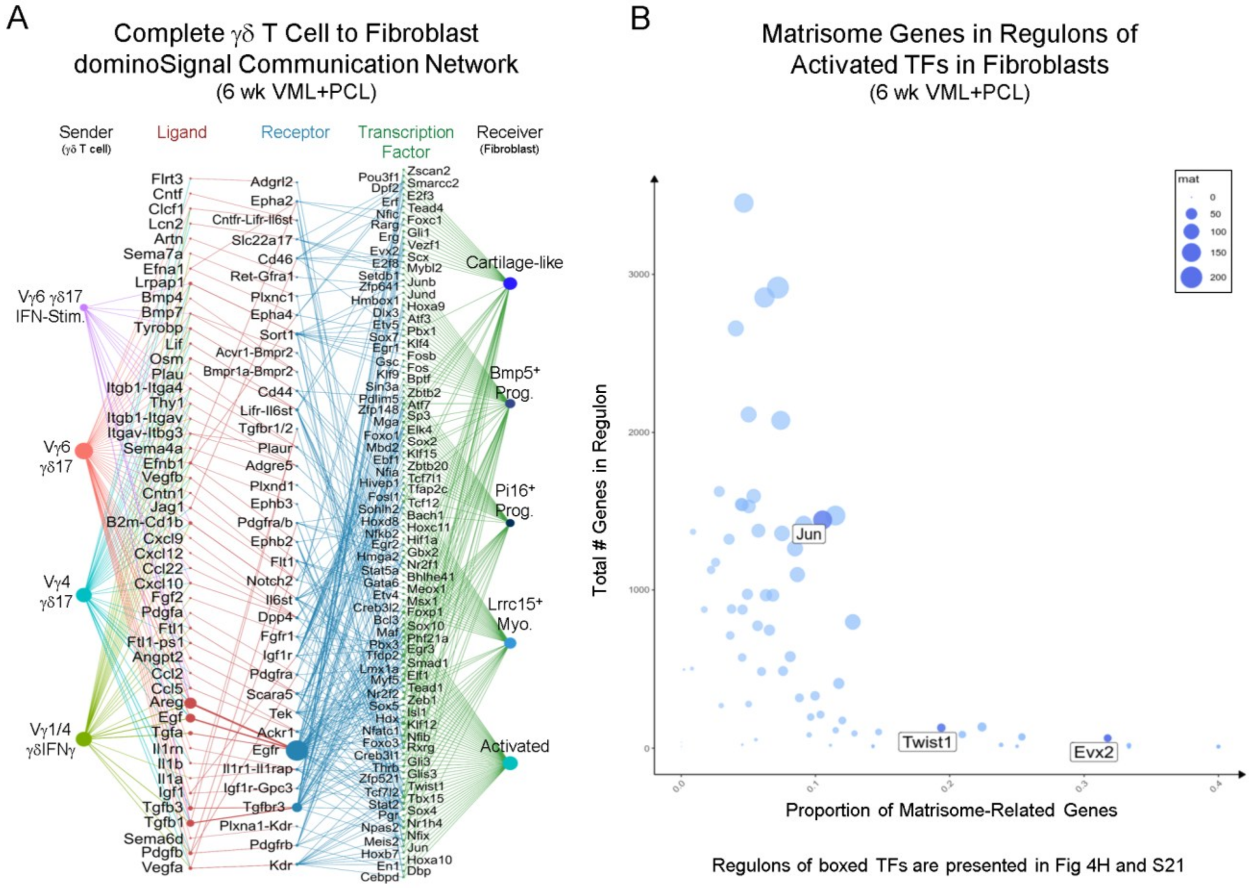

**Suppl Fig 18. Inferred communication from γδ T cells to fibroblasts in implant fibrosis.**
**(A)** Complete inferred dominoSignal network connecting γδ T cell ligands to fibroblast receptors and activated TFs in VML+PCL at 6 wks (No thresholds of ligand/receptor average expression or TF activation scores). **(B)** Scatter plot displaying the total number of genes in the regulon (y axis ) and proportion of matrisome-related genes in regulon (x axis and point size) for TFs activated in the γδ T cell to fibroblast communication network.

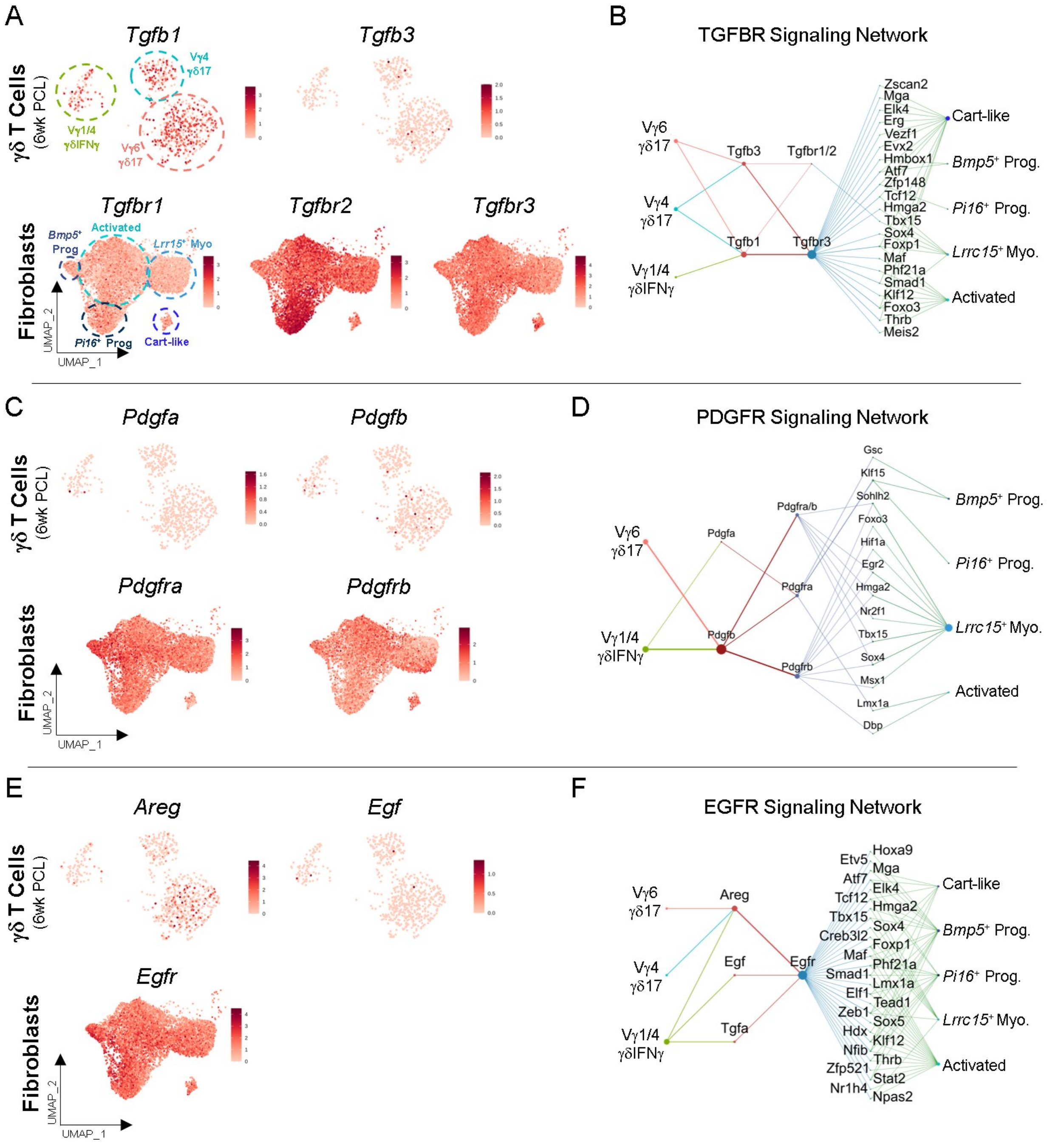

**Suppl Fig 19. Inferred signaling networks between γδ T cells and fibroblasts mediated by growth factors in implant fibrosis.**

**(A)** Feature plots displaying normalized expression (norm. exp.) of TGFB ligands (*Tgfb1, Tgfb3*) by γδ T cells (top) and TGFB receptors (*Tgfbr1, Tgfbr2, Tgfbr3*) by fibroblasts in VML+PCL at 6 wks. **(B)** Complete TGFBR signaling network. **(C)** Feature plots displaying norm. exp. of PDGF ligands (*Pdgfa, Pdgfb*) by γδ T cells (top) and PDGF receptors (*Pdgfra, Pdgfrb*) by fibroblasts in VML+PCL at 6 wks. **(D)** Complete PDGFR signaling network. **(E)** Feature plots displaying norm. exp. of EGF ligands (*Areg, Egf*) by γδ T cells (top) and EGF receptor (*Egfr*) by fibroblasts in VML+PCL at 6 wks. **(F)** Complete EGFR signaling network.

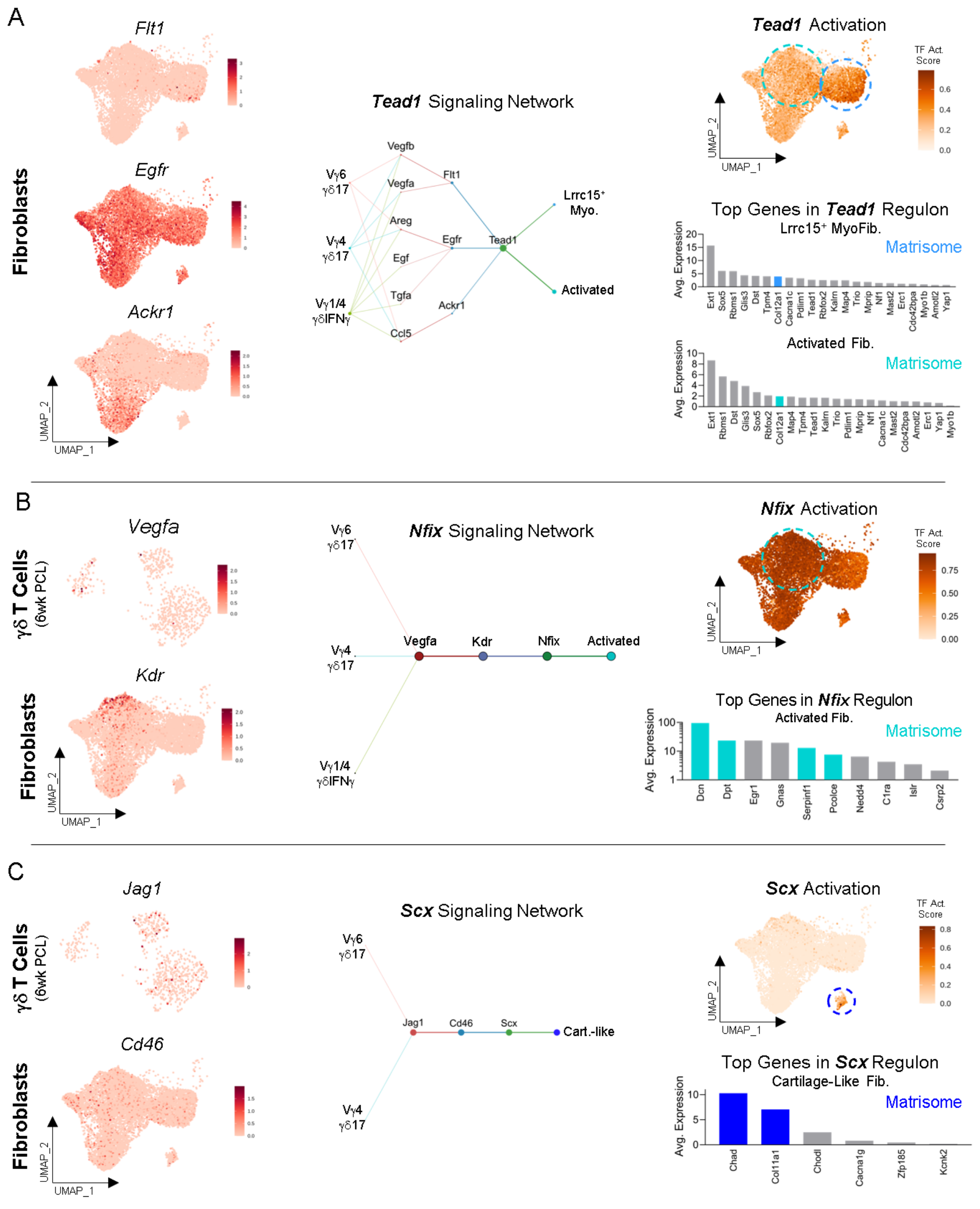

**<u>Suppl Fig 20</u>. Inferred networks for the most activated fibroblast TFs involved in γδ T cell communication.**

**(A)** Feature plots displaying normalized expression (norm. exp.) of receptors (*Flt1, Egfr, Ackr1*) that correlate with fibroblast *Tead1* activation (left). Complete *Tead1* signaling network (middle). Cropped feature plot of *Tead1* activation score (right, top) and average expression (avg. exp.) of top genes in *Tead1* regulon by fibroblast sub-clusters with colored bars indicating matrisome association (right, bottom). **(B)** Feature plots displaying norm. exp. of *Vegfa* by γδ T cells (left, top) and receptor (*Kdr*) that correlates with fibroblast *Nfix* activation (left, bottom). Complete *Nfix* signaling network (middle). Cropped feature plot of *Nfix* activation score (right, top) and avg. exp. of all genes in *Nfix* regulon by 'activated fibroblasts' with colored bars indicating matrisome association (right, bottom). **(C)** Feature plots displaying norm. exp. of *Jag1* by γδ T cells (left, top) and receptor (*Cd46*) that correlates with fibroblast *Scx* activation (left, bottom). Complete *Scx* signaling network (middle). Cropped feature plot of *Scx* activation score (right, top) and avg. exp. of all genes in *Scx* regulon by 'cartilage-like' fibroblasts with colored bars indicating matrisome association (right, bottom).

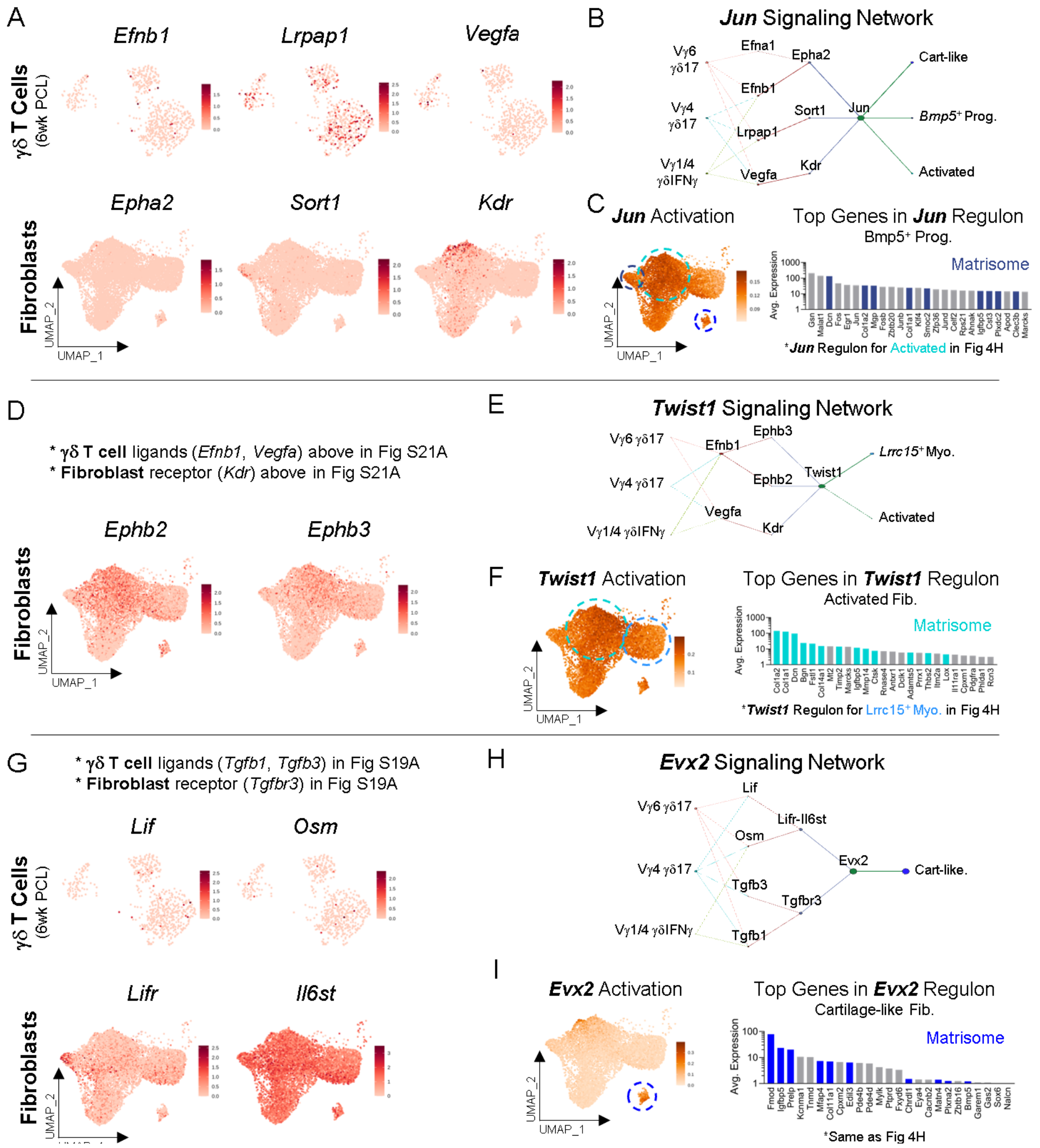

**Suppl Fig 21. Networks for select activated fibroblast TFs involved in γδ T cell communication with matrisome-related genes in TF regulons.**

**(A)** Feature plots displaying normalized expression (norm. exp.) of ligands (*Efnb1, Lrpap1, Vegfa*) by γδ T cells (top) and receptors (*Epha2, Sort1, Kdr*) by fibroblasts (bottom) involved in *Jun* activation. **(B)** Complete *Jun* signaling network. **(C)** Cropped feature plot of *Jun* activation score (left) and average expression (avg. exp.) of top genes in *Jun* regulon by '*Bmp5*+ Prog.' Cluster (right). Colored bars indicate matrisome association. **(D)** Feature plots displaying norm. exp. of receptors (*Ephb2, Ephb3*) by fibroblasts correlated with *Twist1* activation. **(E)** Complete *Twist1* signaling network. **(F)** Cropped feature plot of *Twist1* activation score (left) and avg. exp. of top genes in *Twist1* regulon by 'activated fibroblast' cluster (right). Colored bars indicate matrisome association. **(G)** Feature plots displaying norm. exp. of ligands (*Lif, Osm*) by γδ T cells (top) and receptors (*Lifr, Il6st*) by fibroblasts (bottom) involved in *Evx2* activation. **(H)** Complete *Evx2* signaling network. **(I)** Cropped feature plot of *Evx2* activation score (left) and avg. exp. of top genes in *Evx2* regulon by 'cartilage-like' cluster (right). Colored bars indicate matrisome association.

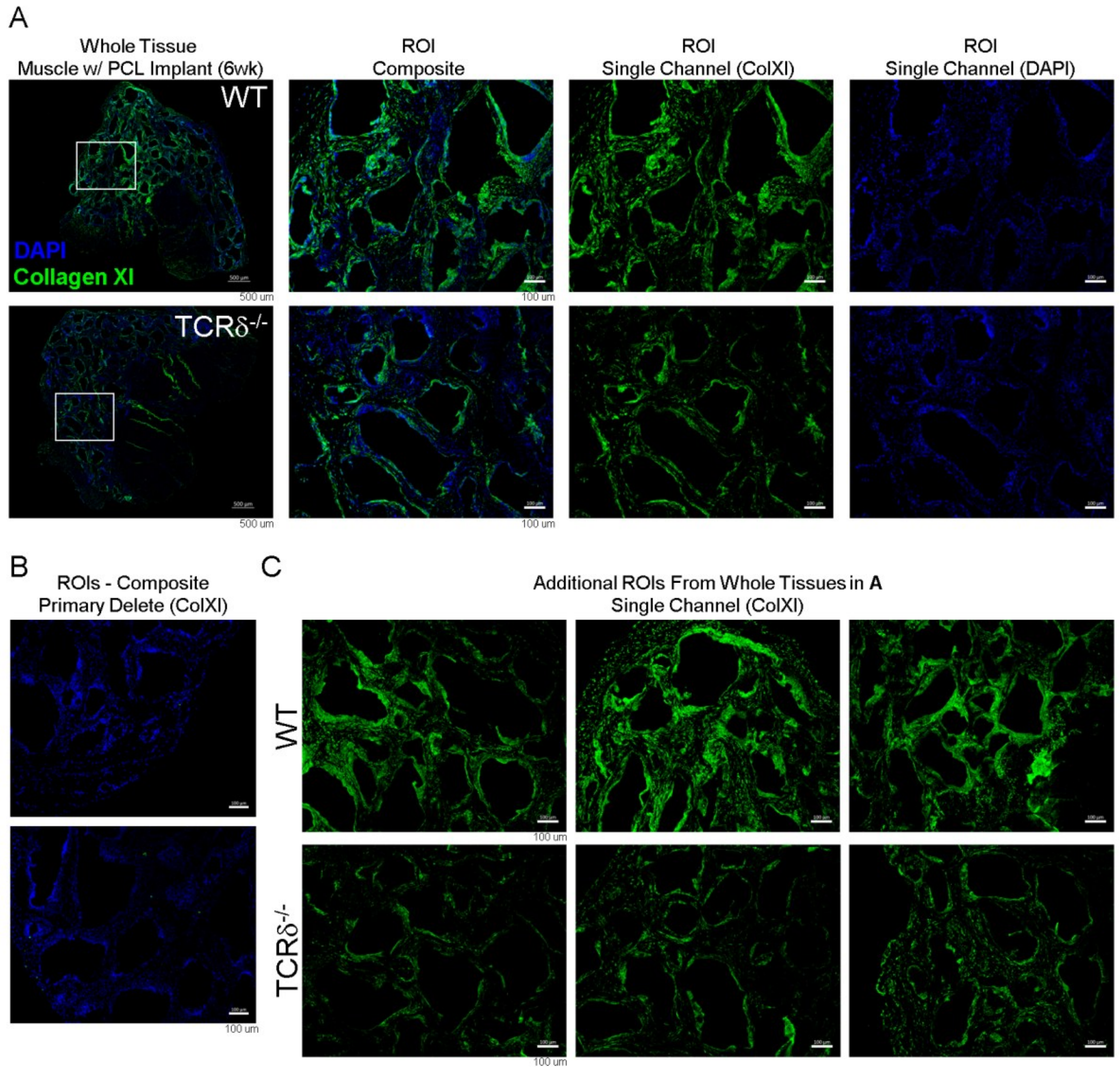

**<u>Suppl Fig 22</u>. Collagen XI in fibrotic tissue surrounding synthetic biomaterial implant.**
**(A)** Representative immunofluorescence staining of collagen XI (green) in fibrotic tissue surrounding PCL implants in WT (top) and TCRδ$^{-/-}$ (bottom) mice after 6 wks. Whole tissue with selected region of interest (ROI) outlined in white box (left) (scale bar: 500μm). Composite and single channel images of selected ROI (middle, right), same as presented in **Fig 5C** (scale bar: 100μm). **(B)** Composite images of primary delete controls for collagen XI antibody (scale bar: 100μm). **(C)** Additional single channel (only collagen XI) ROIs selected from the whole tissue sections of WT (top) and TCRδ$^{-/-}$ (bottom) mice (scale bar: 100μm).

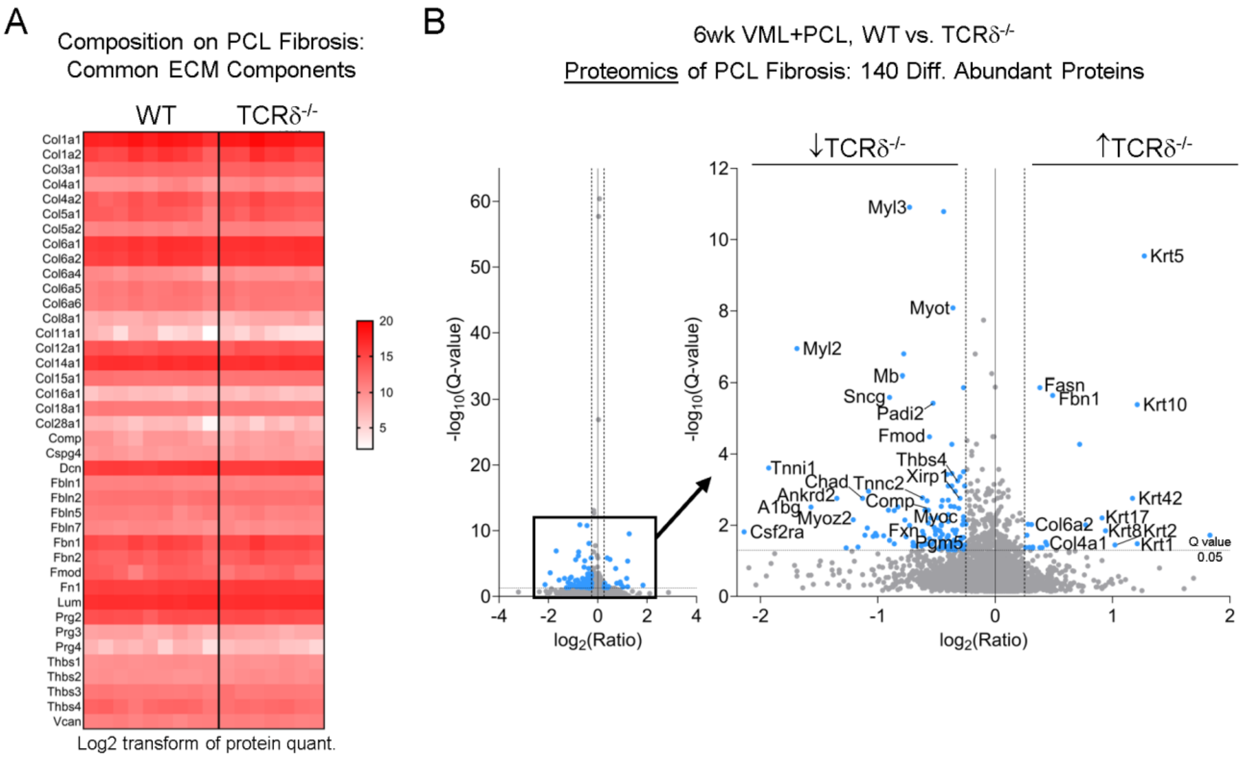

**Suppl Fig 23. Proteomic analysis of fibrotic tissue surrounding synthetic biomaterial implant.**
**(A)** Heatmap of common extracellular matrix (ECM) components (e.g., collagens, proteoglycans) detected in fibrotic tissue surrounding PCL implants after 6 wks in C57BL/6 wildtype (WT) and TCRδ$^{-/-}$ mice. Protein quantification was $\log_2$ transformed prior to heatmap graphing due to large scale range. **(B)** Differential protein abundance (Q val. <0.05, $|\log_2(FC)| \geq 0.25$ indicated in blue) from PCL-associated fibrosis in WT versus TCRδ$^{-/-}$ mice using LC-MS/MS.

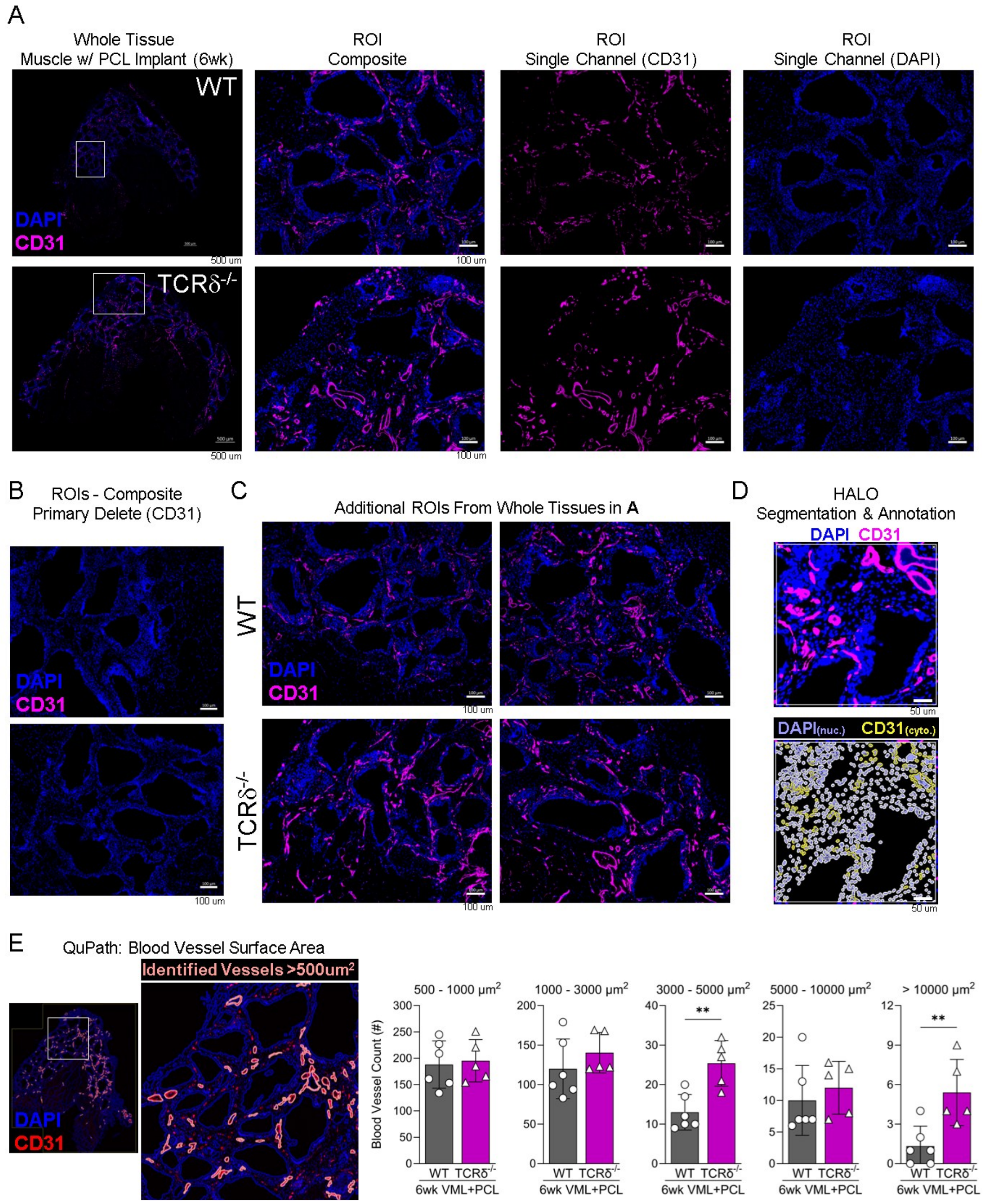

**Suppl Fig 24. Profiling of blood vessels in fibrotic tissue surrounding synthetic biomaterial implant. (A)** Representative immunofluorescence staining of $CD31^+$ blood vessels (pink) in fibrotic tissue surrounding PCL implants in WT (top) and $TCR\delta^{-/-}$ (bottom) mice after 6 wks. Whole tissue with selected region of interest (ROI) outlined in white box (left) (scale bar: 500 μm). Composite and single channel images of selected ROI (middle, right), same as presented in **Fig 6B** (scale bar: 100 μm). **(B)** Composite images of primary delete controls for CD31 (scale bar: 100 μm). **(C)** Additional ROIs selected from whole tissue sections of WT (top) and $TCR\delta^{-/-}$ (bottom) mice (scale bar: 100μm). **(D)** Representative cell segmentation and annotation using HALO for histological quantification of endothelial cell (EC) presented in **Fig 6B**. **(E)** Representative blood vessel annotation using QuPath for lumen surface area measurement (left). Histological quantification of blood vessel lumen area (right). **(Statistics)** Bar graphs: mean±SD. Data analyzed using unpaired two-tailed student t-test (E). NS: Not significant $p>0.05$, * $p<0.05$, ** $p<0.01$.

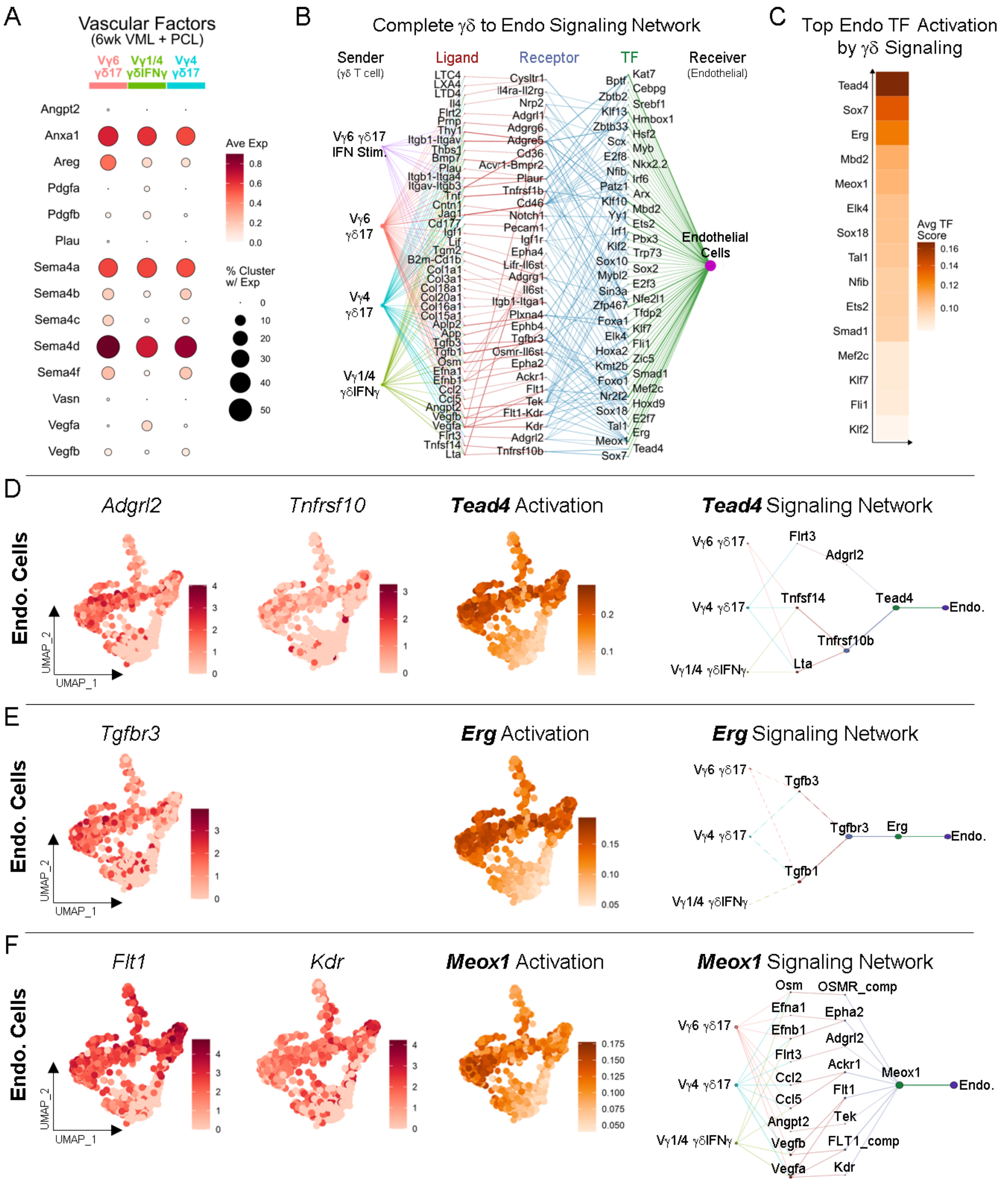

**<u>Suppl Fig 25</u>. Inferred communication from γδ T cells to endothelial cells (ECs) in implant fibrosis.** **(A)** Average expression normalized to maximum expression (Avg. exp.) of secreted vascular factors by scRNAseq γδ T cell clusters in VML+PCL at 6 wks (no thresholds for percentage of cells with detected expression or for avg. exp.). **(B)** Complete dominoSignal network connecting γδ T cell ligands to EC receptors and activated TFs in VML+PCL at 6 wks (no thresholds of ligand/receptor avg. exp. or TF activation scores). **(C)** Top 15 activated EC TFs present in γδ T cell signaling network ranked by avg. TF activation score. **(D)** Cropped feature plots displaying normalized expression (norm. exp.) of receptors (*Adgrl2, Tnfrsf10*) and *Tead4* activation score by ECs (left, middle). Complete *Tead4* signaling network (right). **(E)** Cropped feature plots displaying norm. exp. of receptor (*Tgfbr3*) and *Erg* activation score by ECs (left, middle). Complete *Erg* signaling network (right). **(F)** Feature plots displaying norm. exp. of receptors (*Flt1, Kdr*) and *Meox1* activation score by ECs (left, middle). Complete *Meox1* signaling network (right).

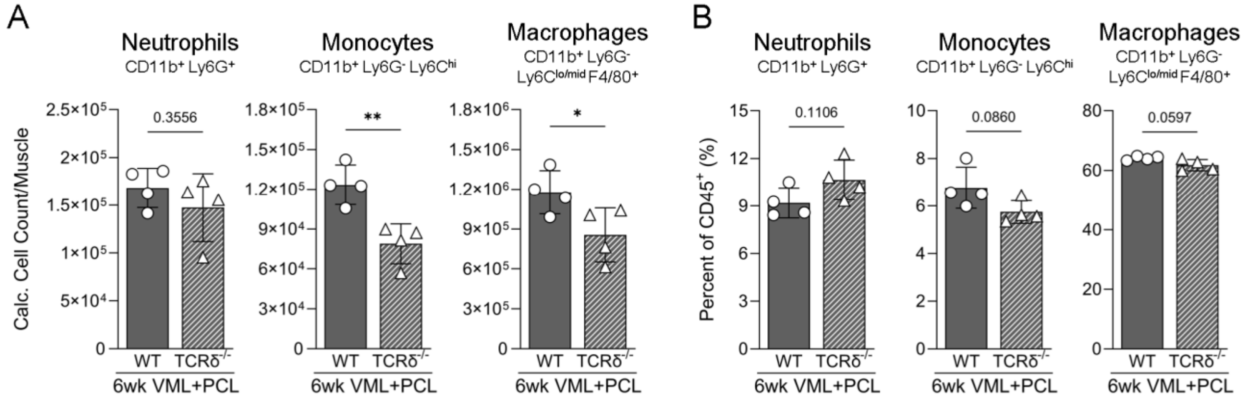

**<u>Suppl Fig 26</u>. Flow cytometric quantification of myeloid cells in TCRδ$^{-/-}$ mice in implant fibrosis.**
**(A)** Quantification of neutrophil (Live CD45$^+$ CD11b$^+$ SiglecF$^-$ Ly6G$^+$), monocyte (Live CD45$^+$ CD11b$^+$ SiglecF$^-$ Ly6G$^-$ Ly6C$^{hi}$), and macrophage (Live CD45$^+$ CD11b$^+$ SiglecF$^-$ Ly6G$^-$ Ly6C$^{lo/mid}$ F4/80$^+$) cell counts in VML-injured muscle with PCL implants (VML+PCL) after 6 wks in WT and TCRδ$^{-/-}$ mice. **(B)** Percentage of neutrophils, monocytes, and macrophages out of CD45$^+$ immune cells in VML+PCL after 6 wks in WT and TCRδ$^{-/-}$ mice. **(Statistics)** Bar graphs: mean±SD. Data analyzed using unpaired two-tailed student t-test (E). NS: Not significant $p>0.05$, * $p<0.05$, ** $p<0.01$.

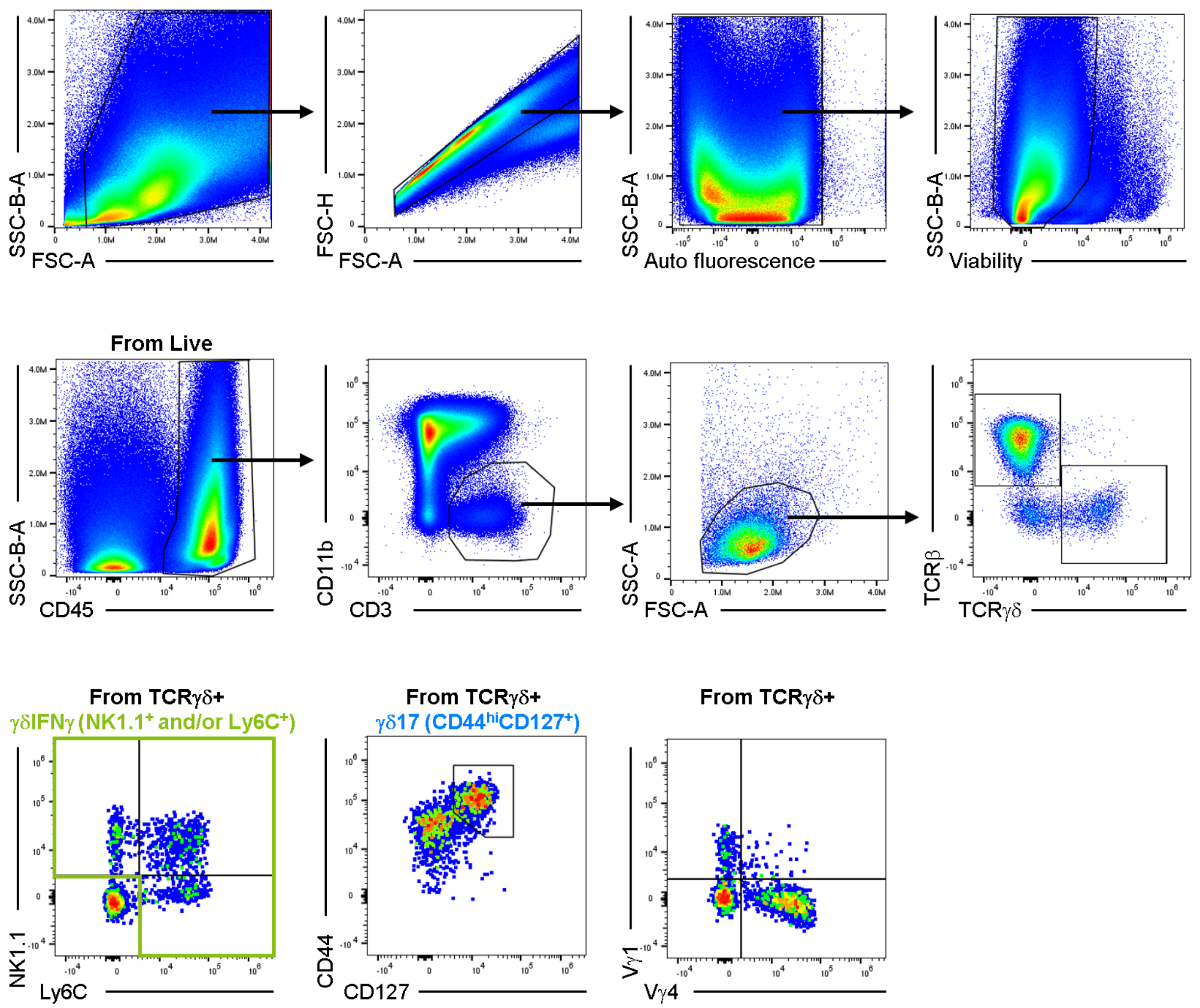

**Gate placement confirmed with FMOs (presented in Suppl Fig 7A)**

**Suppl Fig 27. Flow cytometry gating scheme for γδ T Cell Effector panel (surface markers) in VML-injured muscle with PCL implant.**

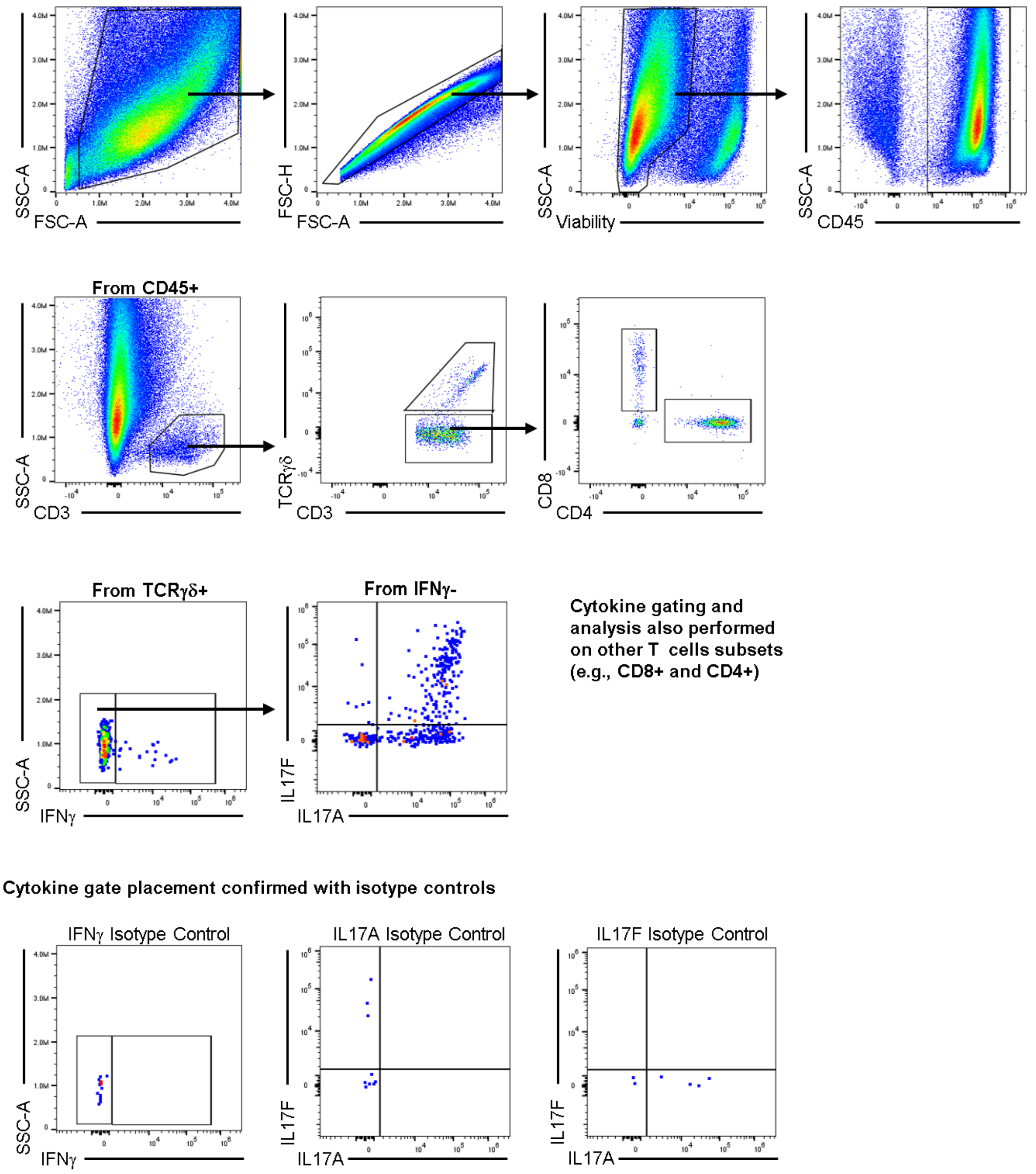

**<u>Suppl Fig 28</u>. Flow cytometry gating scheme for γδ T Cell Intracellular Cytokine panel in VML-injured muscle with PCL implant.**

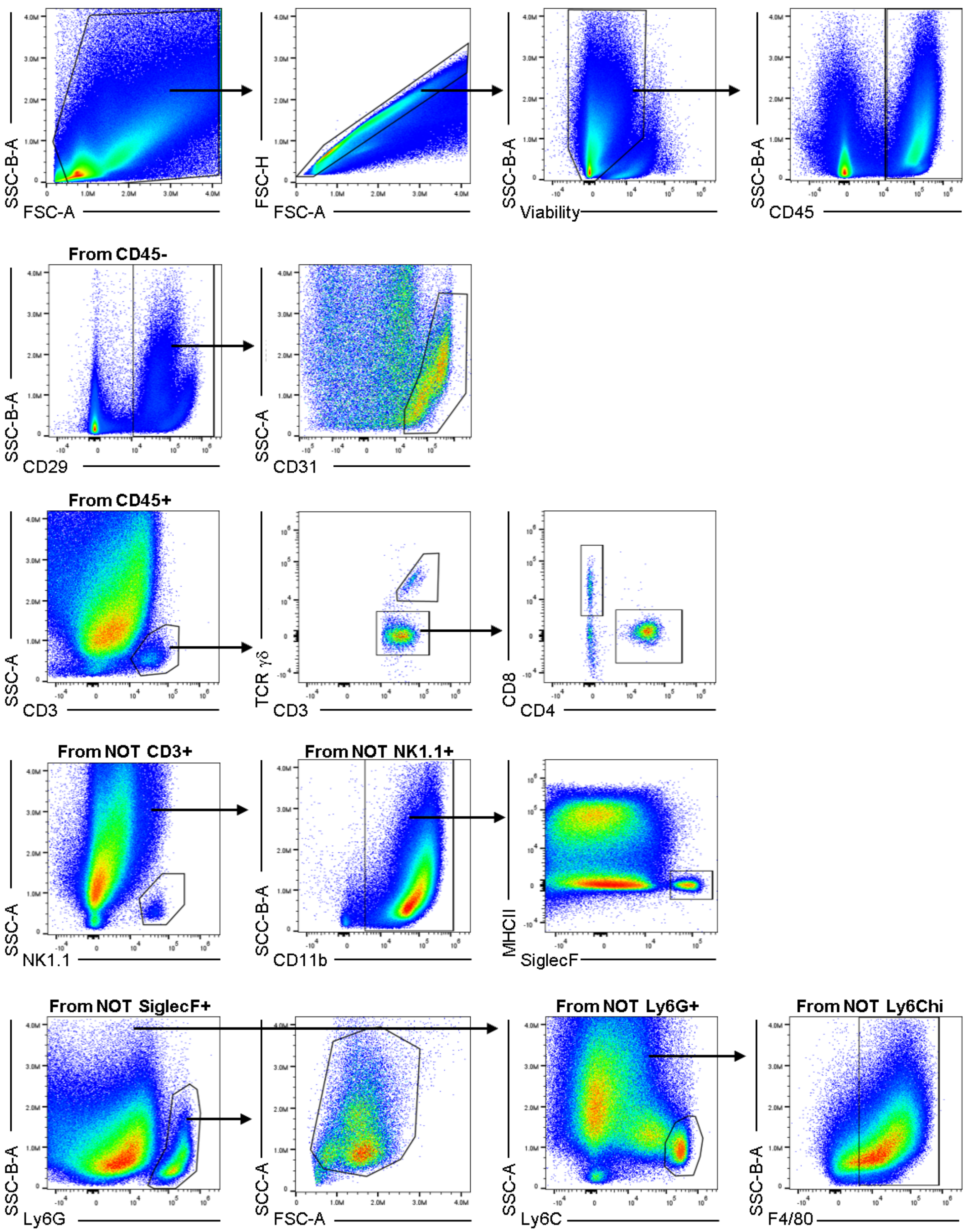

**<u>Suppl Fig 29</u>. Flow cytometry gating scheme for Pan Stroma-Immune panel in VML-injured muscle with PCL implant.**

### Material Tables

| Fluorophore | Antigen | Clone | Dilution | Manufacturer | Catalog # |
|---|---|---|---|---|---|
| BV421 | TCRβ | H57-197 | 100 | BioLegend | 109230 |
| V450 | CD27 | LG.3A10 | 100 | BD Biosciences | 561245 |
| BV480 | KLRG1 | 2F1 | 150 | BD Biosciences | 746353 |
| BV510 | CD45 | 30-F11 | 200 | BioLegend | 103138 |
| BV605 | TCR γ/δ | GL3 | 100 | BioLegend | 118129 |
| BV650 | CD44 | IM7 | 250 | BioLegend | 103049 |
| BV711 | NKG2D | CX5 | 100 | BD Biosciences | 563694 |
| BV750 | B220 | RA3-6B2 | 200 | BioLegend | 103261 |
| BV786 | Vγ1.1 (Vγ1) | 2.11 | 100 | BD Biosciences | 744612 |
| KIRAVIA Blue 520 | CD69 | H1.2F3 | 100 | BioLegend | 104554 |
| Spark Blue 550 | CD3 | 17A2 | 100 | BioLegend | 100260 |
| PerCP | Ly6C | HK1.4 | 250 | BioLegend | 128028 |
| BB700 | CD8α | 53-6.7 | 200 | BD Biosciences | 566409 |
| PerCP-eFluor710 | Vγ2 (Vγ4) | UC3-10A6 | 100 | Thermo Fisher | 46-5828-82 |
| PE | CCR6 | 292L17 | 100 | BioLegend | 129804 |
| PE/Dazzle 594 | CD62L | MEL-14 | 150 | BioLegend | 104448 |
| AF647 | NK1.1 | PK136 | 200 | BioLegend | 108720 |
| AF700 | CD11b | M1/70 | 400 | BioLegend | 101222 |
| Zombie NIR | Viability | - | 5000 | BioLegend | 423106 |
| APC-eFluor780 | CD127 | A7R34 | 150 | Thermo Fisher | 47-1271-82 |
| APC/Fire 810 | CD4 | GK1.5 | 100 | BioLegend | 100480 |
| PE | Vγ6 | 1C10-1F7 | 100 | BD Biosciences | 570217 |

* Swapped in after release of commercial antibody

**Material Table 1**: γδ T Cell Effector Flow Panel

| Fluorophore | Antigen | Clone | Dilution | Manufacturer | Catalog # |
|---|---|---|---|---|---|
| BV421 | SiglecF | E502440 | 150 | BD Biosciences | 562681 |
| Super Bright 436 | CD19 | 1D3 | 100 | Thermo Fisher | 62-0193-82 |
| Pacific Blue | Ly6G | 1A8 | 250 | BioLegend | 127612 |
| BV510 | CD31 | MEC 13.3 | 200 | BD Biosciences | 563089 |
| BV570 | CD45 | 30-F11 | 150 | BioLegend | 103136 |
| BV605 | CD90.2 | 30-H12 | 1500 | BioLegend | 105343 |
| BV650 | Ly6C | HK1.4 | 1200 | BioLegend | 128049 |
| BV711 | TCR γ/δ | GL3 | 200 | BD Biosciences | 563994 |
| BV750 | B220 | RA3-6B2 | 200 | BioLegend | 103261 |
| BV785 | F4/80 | BM8 | 300 | BioLegend | 123141 |
| BB515 | cKit | 2B8 | 500 | BD Biosciences | 564481 |
| Spark Blue 550 | CD3 | 17A2 | 100 | BioLegend | 100260 |
| PerCP | I-A/I-E | M5/114.15.2 | 200 | BioLegend | 107624 |
| BB700 | CD8α | 53-6.7 | 100 | BD Biosciences | 566409 |
| PerCP-eFluor710 | CD29 | HMB1-1 | 800 | Thermo Fisher | 46-0291-82 |
| PE | CD115 | AFS98 | 100 | BioLegend | 135506 |
| PE/Dazzle 594 | CD11c | N418 | 500 | BioLegend | 117348 |
| PE/Cy7 | CD200R3 | Ba13 | 400 | BioLegend | 142212 |
| AF647 | NK1.1 | PK136 | 200 | BioLegend | 108720 |
| AF700 | CD11b | M1/70 | 400 | BioLegend | 101222 |
| Zombie NIR | Viability | - | 3000 | BioLegend | 423106 |
| APC/Fire 810 | CD4 | GK1.5 | 100 | BioLegend | 100480 |

**Material Table 2**: Pan Stroma-Immune Flow Panel

| Fluorophore | Antigen | Clone | Dilution | Manufacturer | Catalog # |
|---|---|---|---|---|---|
| BV421 | Vγ1.1 (Vγ1) | 2.11 | 100 | BD Biosciences | 566308 |
| BV510 | CD45 | 30-F11 | 200 | BioLegend | 103138 |
| BV605 | TCR γ/δ | GL3 | 100 | BioLegend | 118129 |
| BV785 | CD4 | GK1.5 | 200 | BioLegend | 100453 |
| Spark Blue 550 | CD3 | 17A2 | 100 | BioLegend | 100260 |
| BB700 | CD8α | 53-6.7 | 200 | BD Biosciences | 566409 |
| PerCP-eFluor710 | Vγ2 (Vγ4) | UC3-10A6 | 100 | Thermo Fisher | 46-5828-82 |
| PE | IL17F | 9D3.1C8 | 200 | BioLegend | 517008 |
| APC | IFNγ | XMG1.2 | 150 | BioLegend | 505810 |
| AF700 | IL17A | TC11-18H10.1 | 200 | BioLegend | 506914 |
| Zombie NIR | Viability | - | 5000 | BioLegend | 423106 |

| | | | | | |
|---|---|---|---|---|---|
| PE | Mouse IgG1 | MOPC-21 | 200 | BioLegend | 400112 |
| APC | Rat IgG1 | RTK2071 | 150 | BioLegend | 400411 |
| AF700 | Rat IgG1 | RTK2071 | 200 | BioLegend | 400420 |

Red text indicates intracellular antibody staining

**Material Table 3**: Intracellular Cytokine Flow Panel

| Fluorophore | Antigen | Clone | Dilution | Manufacturer | Catalog # |
|---|---|---|---|---|---|
| BV605 | CD45 | 30-F11 | 150 | BioLegend | 103140 |
| AF488 | CD3 | 17A2 | 200 | BioLegend | 100210 |
| Zombie NIR | Viability | - | | BioLegend | 423106 |

| Hashing Oligo. | Antigen | Clone | Dilution | Manufacturer | Catalog # |
|---|---|---|---|---|---|
| TotalSeq-C0301 | MHC class I; CD45 | M1/42; 30-F11 | 100 | BioLegend | 155861 |
| TotalSeq-C0302 | MHC class I; CD45 | M1/42; 30-F11 | 100 | BioLegend | 155863 |
| TotalSeq-C0303 | MHC class I; CD45 | M1/42; 30-F11 | 100 | BioLegend | 155865 |
| TotalSeq-C0304 | MHC class I; CD45 | M1/42; 30-F11 | 100 | BioLegend | 155867 |
| TotalSeq-C0305 | MHC class I; CD45 | M1/42; 30-F11 | 100 | BioLegend | 155869 |

**Material Table 4**: CD3 T Cell FACS Panel for Single-cell RNAseq

| Fluorophore | Antigen | Clone | Dilution | Manufacturer | Catalog # |
|---|---|---|---|---|---|
| BV421 | CD31 | 390 | 400 | BioLegend | 102424 |
| BV510 (Fixable Aqua) | Viability | - | 500 | Thermo Fisher | L34957 |
| BV605 | CD45 | 30-F11 | 250 | BioLegend | 103140 |
| BV711 | CD11b | M1/70 | 600 | BioLegend | 101241 |
| AF488 | Ly6G | 1A8 | 250 | BioLegend | 127626 |
| PE/Cy7 | F4/80 | BM8 | 300 | BioLegend | 123114 |
| APC/Cy7 | CD29 | HMB1-1 | 400 | BioLegend | 102226 |

**Material Table 5**: Stromal and Endothelial Cell FACS Panel for Bulk RNAseq

| Primer Name | Sequence |
|---|---|
| 10X-FP-1 | 5’ AATGATACGGCGACCACCGAGATCTACACTCTTTCCCTACACGACGCTC |
| GDM-RP-1-1 | 5’ TCGAATCTCCATACTGACCAAGCTTGAC |
| GDM-RP-1-2 | 5’ GTCTTCAGCGTATCCCCTTCCTGG |
| GDM-RP-1-3 | 5’ CTTTCAGGCACAGTAAGCCAGC |
| GDM-RP-1-4 | 5’ TCTTCAGTCACCGTCAGCCAACTAA |
| 10X-FP-2 | 5’ AATGATACGGCGACCACCGAGATCT |
| GDM-RP-2-1 | 5’ CCACAATCTTCTTGGATGATCTGAGACT |
| GDM-RP-2-2 | 5’ GTCCCAGTCTTATGGAGATTTGTTTCAGC |

**Material Table 6**: Custom primers for $\gamma\delta$ TCR library prep for paired scRNA/TCRseq.
Originally from Lee et al. (*Nature Commun.*, 2020).

Fibroblasts from Co-Culture

| Gene | TaqMan Probe ID |
|---|---|
| Acan | Mm00545794_m1 |
| Ccl2 | Mm00441242_m1 |
| Col1a1 | Mm00801666_g1 |
| Col2a1 | Mm01309565_m1 |
| Col3a1 | Mm00802300_m1 |
| Col5a1 | Mm00489299_m1 |
| Col9a2 | Mm00483875_m1 |
| Cxcl2 | Mm00436450_m1 |
| Il6 | Mm00446190_m1 |
| Il11 | Mm00434162_m1 |
| Lif | Mm00434762_g1 |
| Nfkbiz | Mm00600522_m1 |
| Prg4 | Mm01284582_m1 |
| Ptgs2 | Mm00478374_m1 |
| Rer1 | Mm00471276_m1 |

VML+PCL in LFD versus HFD

| Gene | TaqMan Probe ID |
|---|---|
| Cdkn1a | Mm00432448_m1 |
| Cdkn2a | Mm00494449_m1 |
| Col1a1 | Mm00801666_g1 |
| Col3a1 | Mm01254477_m1 |
| Col5a3 | Mm00489842_m1 |
| Fap | Mm01329177_m1 |
| Ifng | Mm01168134_m1 |
| Il1b | Mm00434228_m1 |
| Il1f9 | Mm00463327_m1 |
| Il6 | Mm00446190_m1 |
| IL17f | Mm00521423_m1 |
| Ly6g | Mm04934123_m1 |
| Nos2 | Mm00440502_m1 |
| Pdgfa | Mm01205760_m1 |
| Pdgfb | Mm00440677_m1 |
| Ptgs2 | Mm00478374_m1 |
| Rer1 | Mm00471276_m1 |
| S100a4 | Mm00803372_g1 |
| Tgfb1 | Mm01178820_m1 |
| Tgfb2 | Mm00436955_m1 |
| Tnf | Mm00443258_m1 |

**Material Table 7**: TaqMan Probes for RT-qPCR
